# Genomic plasticity of the mating-type loci underlies reproductive strategy transitions in *Rhodotorula* yeasts

**DOI:** 10.1101/2025.09.11.675505

**Authors:** Xin-Zhan Liu, Cheng-Hung Tsai, Marco A. Coelho, Eva Ottum, Cene Gostinčar, Benedetta Turchetti, Claudia Coleine, Laura Selbmann, Ian Wheeldon, Nina Gunde-Cimerman, Feng-Yan Bai, Jason E. Stajich

## Abstract

Transitions from canonical outcrossing to rare, cryptic, or noncanonical reproduction are widespread in fungi, but how mating-type (*MAT*) systems are reorganized at the genomic and population levels during these shifts remains unclear. The coexistence of sexually reproducing species and putatively asexual lineages makes *Rhodotorula* a natural comparative system for addressing this question. We analyzed 249 globally sampled strains using a combination of long-read and short-read sequencing. Phylogenomics resolved three major clades and several closely related species complexes. Chromosome-scale assemblies confirmed physical separation of the pheromone/receptor (*P/R*) and homeodomain (*HD*) loci, whereas non-random *P/R*–*HD* associations revealed constrained tetrapolar inheritance. The *P/R* locus showed conserved synteny within mating types but extensive structural divergence between A1 and A2 alleles, whereas the *HD* locus remained compact and conserved. Mating-type distributions varied across clades, with both A1 and A2 present in Clade C, lineage-structured partitioning in Clade B, and pronounced A2 dominance in Clade A. Genome-wide linkage disequilibrium (LD) decay in putatively asexual *R*. *mucilaginosa*, although slower than in sexual *R*. *diobovata*, was consistent with historical recombination in a species without an observed sexual cycle. Clade A further exhibited deep trans-specific polymorphism of *STE3*.*A2* and relaxed purifying selection, mosaic A2 *P*/*R* configurations retaining *RHA.A1*, A1/A2 and A2/A2 hybrid genomes, and A2-associated gene content beyond *MAT* loci. In conclusion, our findings underscore the dynamic nature of *MAT*-locus remodeling in *Rhodotorula* and establish this genus as a valuable model for investigating transitions from heterothallism toward reproductive strategies less dependent on opposite-type partners.

## Introduction

Sexual reproduction is a pervasive feature of eukaryotic life, likely present in the last eukaryotic common ancestor and broadly conserved across major lineages as a major evolutionary innovation (Heitman et al. 2013; Heitman 2015). Meiotic recombination reshuffles genetic variation and enhances the efficacy of natural selection by combining beneficial mutations together and purging deleterious alleles, thereby promoting adaptation (Yadav et al. 2023). In fungi, sexual reproduction has contributed substantially to morphological, biological, and ecological diversification (Virágh et al. 2022) and can promote speciation through reproductive isolation mediated by the evolution of mating compatibility systems and reproductive genes (Idnurm 2011). In pathogenic fungi, sexual reproduction can promote virulence evolution by generating infectious spores with increased pathogenicity or fitness (Hsueh and Heitman 2008). In *Ustilago maydis*, the mating-induced dimorphic switch is required for plant infection (Lanver et al. 2017). Knowledge of fungal sexual systems also provides a basis for controlled crossing and strain improvement in industrial and biotechnological applications (Kück and Böhm 2013). Despite these evolutionary and practical advantages, transitions toward infrequent, cryptic, or apparently absent sexual reproduction have occurred repeatedly across eukaryotic lineages (Heitman 2015). How the genetic and genomic machinery of sexual reproduction is remodeled during such transitions remains a central question in evolutionary biology.

In fungi, reproductive mode cannot be inferred reliably from the presence or absence of observed sexual structures. Sexual identity in fungi is governed by mating-type (*MAT*) loci rather than differentiated male and female sex chromosomes as observed in animals and plants, and many lineages without detectable sexual stage retain conserved *MAT* genes and core meiotic machinery (Kües et al. 2011; Hofstatter and Lahr 2019). These genomic features indicate the potential for sex but do not demonstrate whether, how, or how often it occurs (Nieuwenhuis and James 2016). Population genetic and genomic studies instead reveal a continuum of reproductive strategies, from regular outcrossing to predominantly clonal propagation punctuated by rare, cryptic, selfing, or other noncanonical sexual cycles (Dyer and O’Gorman 2011; Beukeboom and Perrin 2014; Nieuwenhuis and James 2016). While clonal reproduction enables rapid expansion in favorable environments, these intermittent sexual cycles preserve opportunities for recombination, the removal of deleterious mutations, and the generation of adaptive diversity under specific ecological conditions. Apparent asexuality may therefore reflect a change in the frequency, genetic control, ecological context, or genomics consequences of sexual reproduction rather than its complete loss. Fungal mating systems thus provide a useful framework for studying how reproductive strategies evolve through changes in mating compatibility, *MAT* architecture, and recombination.

In tetrapolar basidiomycetes, canonical heterothallism requires compatible partners carrying distinct alleles at both the pheromone/receptor (*P/R*) and homeodomain (*HD*) loci. The *P/R* locus governs premating recognition and cell–cell fusion, while the *HD* locus regulates post-fusion sexual development, together ensuring that sexual reproduction occurs strictly between compatible non-self lineages (Kües et al. 2011). Strong mating-type bias or loss of a compatible mating type can therefore constrain conventional outcrossing without necessarily eliminating sex. Fungi have repeatedly evolved alternative reproductive strategies that reduce dependence on encounters with compatible partners while retaining ploidy transitions, meiosis, recombination, and the genetic diversity (Heitman et al. 2013). These strategies include several forms of self-fertile or mate-independent reproduction. Homothallism can arise either through the coexistence of compatible mating-type determinants within the same genome or mating-type switching (Krassowski et al. 2019). In pseudo-homothallism, nuclei of opposite mating-types are packaged within a single spore (Wilson et al. 2015). Same-sex mating, also known as unisexual reproduction, allows the sexual cycle between cells of the same mating type, as exemplified by *Neurospora africana*, *Cryptococcus deneoformans*, and *C. neoformans* (Glass and Smith 1994; Lin et al. 2005; Sun et al. 2025). These reproductive modes can preserve evolutionary benefits of sex while reducing the costs of finding a compatible mate, especially in host-associated or otherwise mate-limited environments (Fu et al. 2015).

Changes in mating compatibility are often accompanied by remodeling *MAT*-locus architecture and linkage relationships. *Basidiomycete* mating systems range from tetrapolar architectures, in which *P/R* and *HD* loci are physically unlinked and assort independently, to bipolar systems, in which linkage between the two loci reduces mating-type diversity and simplifies compatibility (Kües et al. 2011; Maia et al. 2015). Multiple independent transitions between tetrapolar and bipolar systems have occurred in *Trichosporonales*, *Cryptococcus*, *Kwoniella*, *Malassezia*, *Ustilago*, and anther-smut fungi through recombination suppression, chromosomal rearrangements, and the formation of mating-type supergenes (Lee et al. 1999; Fraser et al. 2004; Branco et al. 2018; Sun et al. 2019; Coelho et al. 2023, 2025). Intermediate configurations have also been described. In pseudo-bipolar systems, the *P/R* and *HD* loci are linked on the same chromosome but retain occasional recombination (Coelho et al. 2023), whereas the physically unlinked *MAT* loci of *Leucosporidium scottii* show non-random associations, demonstrating that a structurally tetrapolar system can exhibit constrained inheritance (Maia et al. 2015). These changes are frequently associated with ecological or life-history shifts in which limited partner availability favors increased mating efficiency and the preservation of favorable *MAT* combinations (Hsueh and Heitman 2008; Heitman et al. 2013; Coelho et al. 2023). *MAT*-locus architecture, inheritance, and recombination therefore provide a genomic framework for tracing reproductive transitions.

Transitions between canonical sex, cryptic recombination, and asexual propagation leave distinct, complementary footprints across the genome. In a species without an obvious sexual stage, meiotic recombination is regarded as one of the most robust pieces of evidence for cryptic sex (Nieuwenhuis and James 2016). Recombination in natural populations has been detected in fungi historically regarded as asexual, or lacking any known morphological evidence of sex, including *Aspergillus* and *Fusarium* (Dyer and O’Gorman 2011; Drott et al. 2020). Genome-wide linkage disequilibrium (LD) decay offers a comparative measure of recombination, with shorter half-decay distances generally associated with frequent outcrossing and slower decay with predominantly clonal reproduction (Taylor et al. 2015; Nieuwenhuis and James 2016). Trans-specific polymorphisms at *MAT* loci reflect ancient allele divergence and long-term maintenance of mating-recognition diversity, whereas shifts toward relaxed selection point to weakened constraints on canonical opposite-sex recognition (Fraser et al. 2004; Branco et al. 2017; Peris et al. 2022; Coelho et al. 2025). Inversions, transposable element accumulation, and pheromone-gene duplication can further reshape *MAT*-locus structure and local recombination (Branco et al. 2018; Coelho et al. 2023, 2025). Hybrid genomes provide another line of evidence, of which A1/A2 diploids are consistent with canonical mating between compatible types, whereas A2/A2 diploids may preserve genomic footprints of noncanonical or same-sex-like compatibility (Lin et al. 2007, 2009; Takashima et al. 2018; Steenwyk et al. 2020; Theelen et al. 2022).

Species of the red yeast genus *Rhodotorula* (*Sporidiobolales*, *Microbotryomycetes*) inhabit remarkably diverse and extreme environments, exhibiting broad adaptive capacity alongside significant biotechnological and clinical relevance. Strains have been isolated from acidic mine waters, high-salinity soils, Antarctic permafrost, glaciers, and even the International Space Station (Sampaio 2011; Buzzini et al. 2018; Coleine et al. 2020; Daudu et al. 2020). Beyond their ecological versatility, *Rhodotorula* species are also of industrial interest due to their ability to produce carotenoids and intracellular lipids that confer tolerance against cold, UV radiation, and oxidative stress (Moliné et al. 2009; Rossi et al. 2009; Park et al. 2018). Furthermore, while they constitute key components of human intestinal or skin mycobiota (Raimondi et al. 2019; Vial et al. 2024), they are also increasingly reported as opportunistic human pathogens, particularly in immunocompromised patients (Ioannou et al. 2019; Gil et al. 2023). Of the 19 currently recognized species, however, sexual reproduction has been observed in only six (Sampaio 2011; Wang et al. 2015; Crous et al. 2017; Tiwari et al. 2021; Touchette et al. 2022). The coexistence of sexually characterized species and lineages lacking an observed sexual cycle makes *Rhodotorula* an ideal model for investigating how *MAT* locus plasticity and associated genomic signatures relate to transitions in reproductive systems.

Previous studies showed that mating-type determinants remain structurally intact across both sexual and putatively asexual *Rhodotorula* lineages. PCR and genome-walking analyses identified *MAT*-linked genes and suggested a pseudo-bipolar system, wherein restricted recombination between mating-type determinants maintains bipolar-like inheritance (Coelho et al. 2008, 2010, 2011). However, the lack of chromosomal-scale assemblies left the physical relationship between the *P/R* and *HD* loci unresolved. Genomic studies of mating-system evolution in *Microbotryomycetes* have also focused on a limited number of taxa, particularly *Leucosporidiales* and *Microbotryum* (Maia et al. 2015; Branco et al. 2017, 2018; Duhamel et al. 2023). Consequently, it remains unclear whether the *P/R* and *HD* loci are physically linked in *Rhodotorula*, how mating-type alleles are distributed within natural populations, and how *MAT*-locus evolution, local recombination, and hybridization differ between sexual and putatively asexual species.

Here, we analyzed 249 globally sampled *Rhodotorula* strains using Oxford Nanopore, PacBio HiFi, and Illumina sequencing to test how *MAT*-locus plasticity and associated genomic differentiation accompany transitions in mating compatibility and reproductive strategy. We established a genus-wide phylogenomic framework, localized the *P/R* and *HD* loci on chromosome-scale assemblies, compared their organization across species and mating types, quantified mating-type allele distributions, and assessed genome-wide linkage disequilibrium. We also examined the evolutionary dynamics and selective regimes of *STE3* alleles, reconstructed subgenomes of representative diploid hybrids to infer their parental origins and tested whether A2 dominance in Clade A was accompanied by broader genomic differentiation. Our analyses revealed that *Rhodotorula* retains physically tetrapolar *MAT* loci but their inheritance is constrained by non-random *P/R*–*HD* associations and a structurally plastic *P/R* region. In A2-dominant Clade A, mating-type imbalance, ancient trans-specific divergence and relaxed purifying selection of *STE3.A2*, mosaic A2 *P/R* configurations, and A1/A2 and A2/A2 hybrid genomes together suggest a stepwise transition from canonical tetrapolar sexuality toward constrained reassortment and lower dependence on opposite-type partners.

## Results

### A genus-wide genomic framework resolved three major *Rhodotorula* lineages and recently diverged species complexes

Genus-wide phylogenetic inference on Rhodotorula yielded a fully congruent topology resolving three major clades, designated Clades A, B, and C. Clades A and B forming sister groups, with Clade C branching as the sister lineage to their common ancestor (Fig. 1), congruent with recent phylogenomic analyses of Sporidiobolales (Kobayashi et al. 2025). Several recently diverged groups nevertheless exhibited unstable or method-dependent placements, most notably the Clade A complex containing *R. mucilaginosa* Y-2510, *R. frigidialcoholis* JG-1b, and *R.* aff. *mucilaginosa* JY1105 (Fig. 1 and Fig. S1). Additional genome comparisons and supporting statistics are provided in Supplementary text S1, Fig. S2, and Table S3.

**Figure 1.**
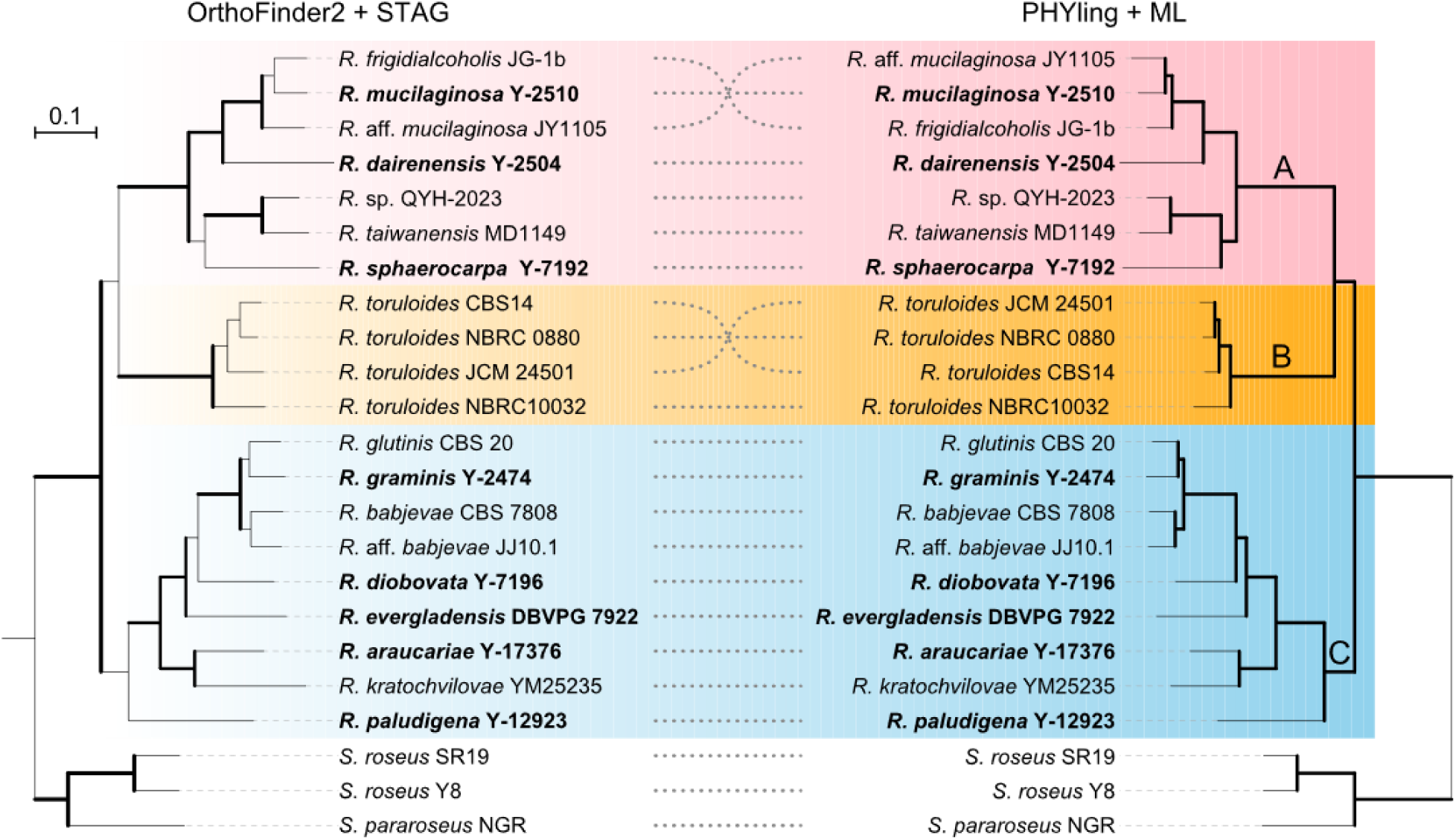
Species tree of the genus *Rhodotorula*. The coalescent species tree on the left was inferred by OrthoFinder (STAG), with bold branches indicating support values above 0.75 (gene tree proportion supporting the corresponding bipartition). The concatenated maximum-likelihood species tree on the right was inferred by Phying, with bold branches indicating bootstrap support above 96% (1,000 replicates). Strains sequenced in this study are shown in bold type. The tree is rooted using three outgroup species (*Sporobolomyces pararoseus* NGR, *S. roseus* SR19, and *S. roseus* Y8). Clades A, B, and C are color shaded and labeled on the right. Underlying data are available in Table S1.

Interspecific comparisons revealed that most species pairs exhibited average nucleotide identity (ANI) values below 90%. Three species groups showed elevated sequence conservation with ANI values of 90–95%: (i) *R. mucilaginosa*, *R.* aff*. mucilaginosa*, and *R. frigidialcoholis*, (ii) *R.* aff. *babjevae*, *R. babjevae*, *R. graminis*, and *R. glutinis*, and (iii) the *R. toruloides* species complex (Fig. S3). Synteny comparisons by NUCmer within these groups revealed small-scale interchromosomal translocations and inversions, with genome-wide sequence identity below 90% (Fig. S4). Within the *R. toruloides* species complex, genome-wide divergence closely paralleled mating type differentiation, consistent with previous observations of sequences and gene-order differences between compatible mating types (Fig. 2 and Fig. S5) (Brink et al. 2023). These analyses establish an enhanced genus-wide phylogenomic and taxonomic framework used for subsequent comparison of *MAT* locus architecture and mating-type allele inheritance.

**Figure 2.**
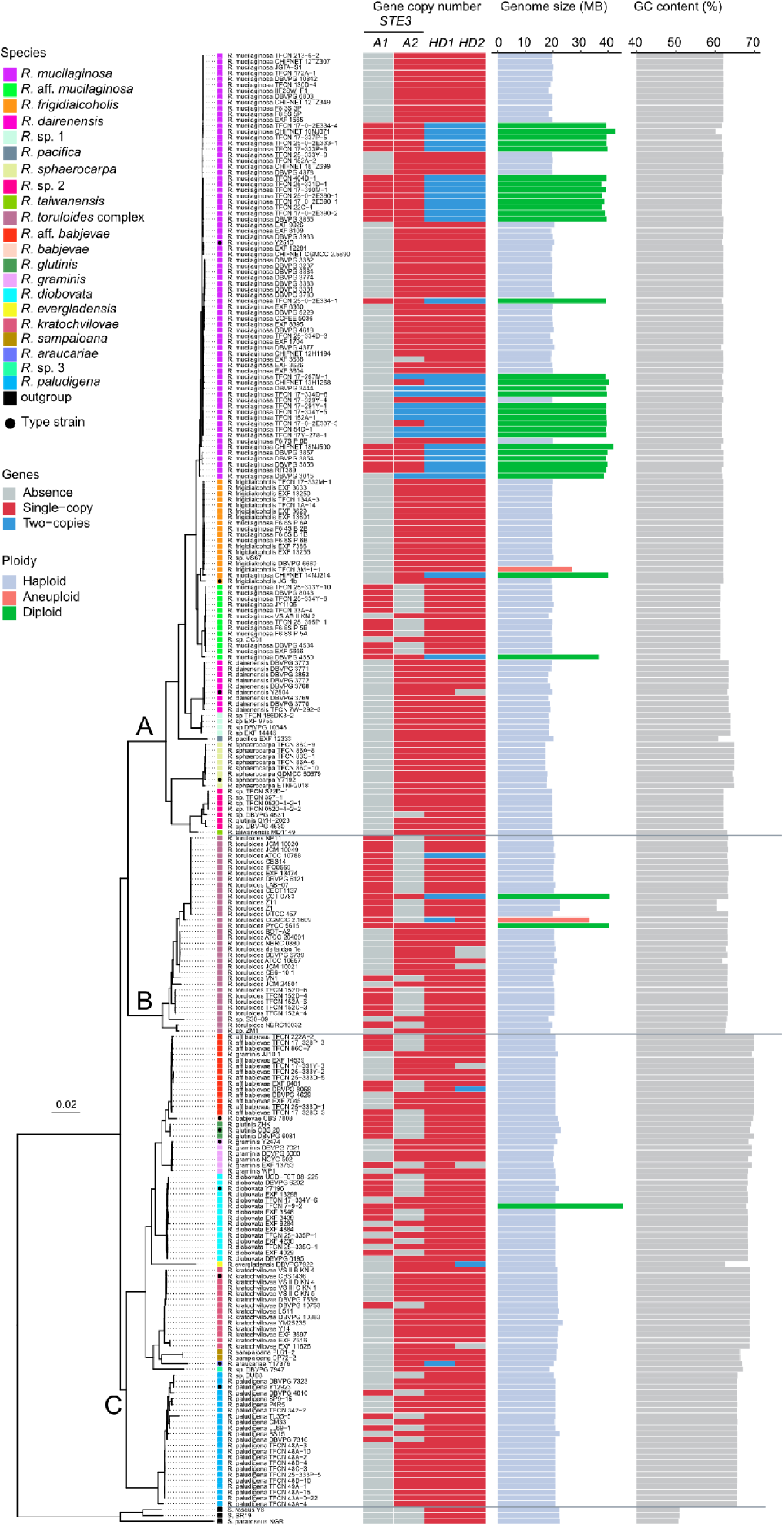
Phylogenetic relationships and genomic features of 249 global *Rhodotorula* strains. The maximum likelihood tree on the left was constructed from a concatenated five-gene alignment (ITS, LSU D1/D2, *RPB1*, *RPB2*, and *TEF1*). Clades A, B, and C are indicated. Tips are color-coded by species/complex. Type strains are marked with circles. Genomic features on the right include: (1) gene copy numbers of key mating-type genes (*STE3.A1*, *STE3.A2*, and *HD1*/*HD2*), indicating mating allele combinations; (2) genome size (Mb) and predicted ploidy level; (3) genomic GC content (%). Underlying data are available in Table S2.

### The *P*/*R* and *HD* loci are physically unlinked but segregate non-randomly in *Rhodotorula*

Previous studies in *Sporidiobolales*, particularly in *Sporidiobolus salmonicolor*, proposed a pseudo-bipolar mating system characterized by bipolar-like mating behavior despite occasional recombination between the *P/R* and *HD* loci (Coelho et al. 2010). Lacking chromosome-scale assemblies, earlier work hypothesized that these two *MAT* loci were partially linked across a distance of at least 800 kb (Coelho et al. 2008). To resolve their physical organization, we examined the chromosomal positions of both loci across our representative genome dataset. We established that the *P/R* and *HD* loci reside on separate chromosomes using two complementary lines of evidence. First, in nine of the eleven newly sequenced species, the *P/R* and *HD* loci mapped to distinct chromosome-level scaffolds, both of which were end-to-end complete with telomeres detected at both ends (Table S1). Second, for assemblies lacking complete telomeric caps, putative centromeric regions suggested that the two loci reside on separate linkage groups across *R.* aff. *babjevae*, *R. glutinis* and members of the *R. toruloides* complex (Fig. S6). Consequently, we found no evidence for physical linkage of the *P/R* and *HD* loci in the representative genomes, indicating that *Rhodotorula* retains a structurally tetrapolar *MAT* organization.

We next evaluated whether the physically unlinked *P/R* and *HD* loci exhibit independent assortment in natural populations, as expected under a canonical tetrapolar system. Under free reassortment, each *P/R* allele should pair with multiple *HD* alleles, and conversely, each *HD* allele should occur across both *P/R* A1 and A2 genetic backgrounds. We examined associations between *STE3* (representing the *P/R* locus) and *HD1*/*HD2* across the 249 strains (Table S4). As expected, the *P/R* locus was biallelic (comprising A1 and A2 types), whereas the *HD* locus was multiallelic. While each *STE3* allele was indeed associated with several *HD* variants, indicating that the two loci are not completely locked into a single fixed combination, most *HD* alleles were restricted to a single *STE3* allele. This generated an asymmetric “one-to-many” association pattern between the two loci (Fig. S7). Reassorted *MAT* allele combinations were detected in only a small subset of strains, including *R*. aff. *mucilaginosa* (e.g., CC01 vs. EXF 5666 or VS_AB_II_KN_2 vs. JY1105), *R.* aff. *babjevae* (EXF 6481 vs. TFCN 17-331Y-3), and *R. diobovata* (e.g., EXF 9284 vs. EXF 4884) (Table S4). Thus, although the *P/R* and *HD* loci are physically unlinked, they do not reassort freely in the natural populations—a pattern we defined as constrained tetrapolar inheritance.

### Contrasting structural dynamics of the *P*/*R* and *HD* loci across *Rhodotorula*

Comparative synteny revealed a conserved core gene set within the *P/R* locus across *Rhodotorula*. This core comprises mating-related genes encoding the pheromone receptor *STE3*, the pheromone precursor *RHA*, *STE20*, and a *STE12*-like protein, alongside a recurrently associated gene cluster, including *KAP95*, *LSM7*, *RIBL6*, *RPAC1*, *DDOST*, *RIBOSOMAL_S19*, *RRM*, *RIBL18AE* (Supplementary text S1 and Table S5). Clade A species and *R. graminis* in Clade C retained the most compact *P/R* gene repertoire, with the core region flanked by *APO* on the left and *MITCARR1* on the right. In contrast, *R. toruloides* complex of Clade B and Clade C species (*R.* aff. *babjevae*, *R. diobovata*, *R. kratochvilovae*, and *R. paludigena*) exhibited an extended locus boundary, incorporating up to seven additional genes on the right side: *MITCARR1*, *ABC1*, *DCN1*, *MRD1*, *SNC*, *SRP9*, and *PREFOLDIN3* (Fig. 4, Fig. S8B, C). Several genes residing within or flanking the expanded *P/R* locus of *Rhodotorula* also occur at or near the *P/R* locus of *Leucosporidium scottii*, a more basally diverging lineage (Maia et al. 2015). In *Rhodosporidiobolus fluvialis*, a sister genus to *Rhodotorula*, this region displays variable architecture, ranging from a collinear *P/R*-adjacent configuration in the heterozygous strain TZ579 to gene deletion and inversion in the A2 strain NJ103 (Huang et al. 2024). Thus, differences in *P/R* gene content among *Rhodotorula* clades reflect variation in locus boundaries and the differential inclusion of flanking genes rather than replacement of the conserved core.

Within each mating type, gene order and organization were generally conserved across species, indicating that A1 and A2 alleles have each been maintained as relatively stable haplotypic configurations (Fig. 3). Nevertheless, a small number of species- and strain-specific genomic rearrangements were detected within these otherwise conserved *P/R* allele classes. Among A2 alleles, structural variations included changes in the position or orientation of *STE20* as well as an inversion of the *LSM7*-*KAP95* module (Supplementary text S1). In A2-dominated Clade A species, strain-specific rearrangements typically involved one or two inverted regions (Fig. S8A and Supplementary text S1). Among A1 alleles, lineage-specific rearrangements included inversion of *STE20* in several Clade C species and a larger module containing *MRD1*, *SRP9*, *SNC1*, *DCN1* and *ABC1* in selected taxa (Supplementary text S1). Overall, the *P/R* locus architecture exhibits strong synteny within each mating-type, with limited structural variations across specific lineages and strain.

**Figure 3.**
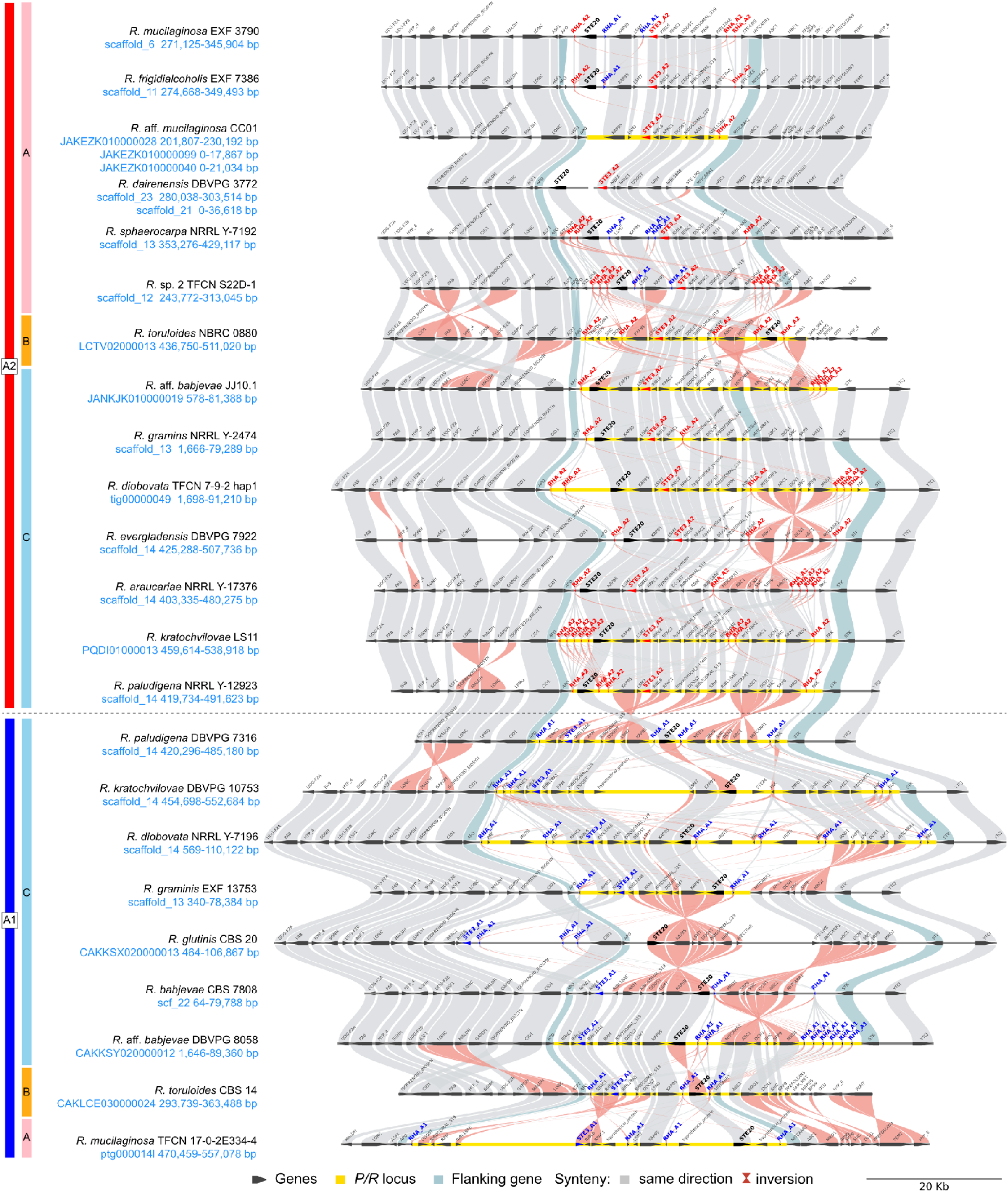
Synteny and gene organization of the *P/R* locus across *Rhodotorula* species and mating types. Comparative synteny map highlights species- and mating-type-specific rearrangements. A1 and A2 *MAT* alleles are labeled on the left. Canonical alleles, *STE3.A2* and *RHA.A2* (red), versus *STE3.A1* and *RHA.A1* (blue) are highlighted. *STE20* is shown in bold black. Homologous genes between adjacent strains are connected by colored shades, where gray indicates collinearity, red indicates inversion, and cyan-blue delineates the predicted *P/R* locus boundaries.

**Figure 4.**
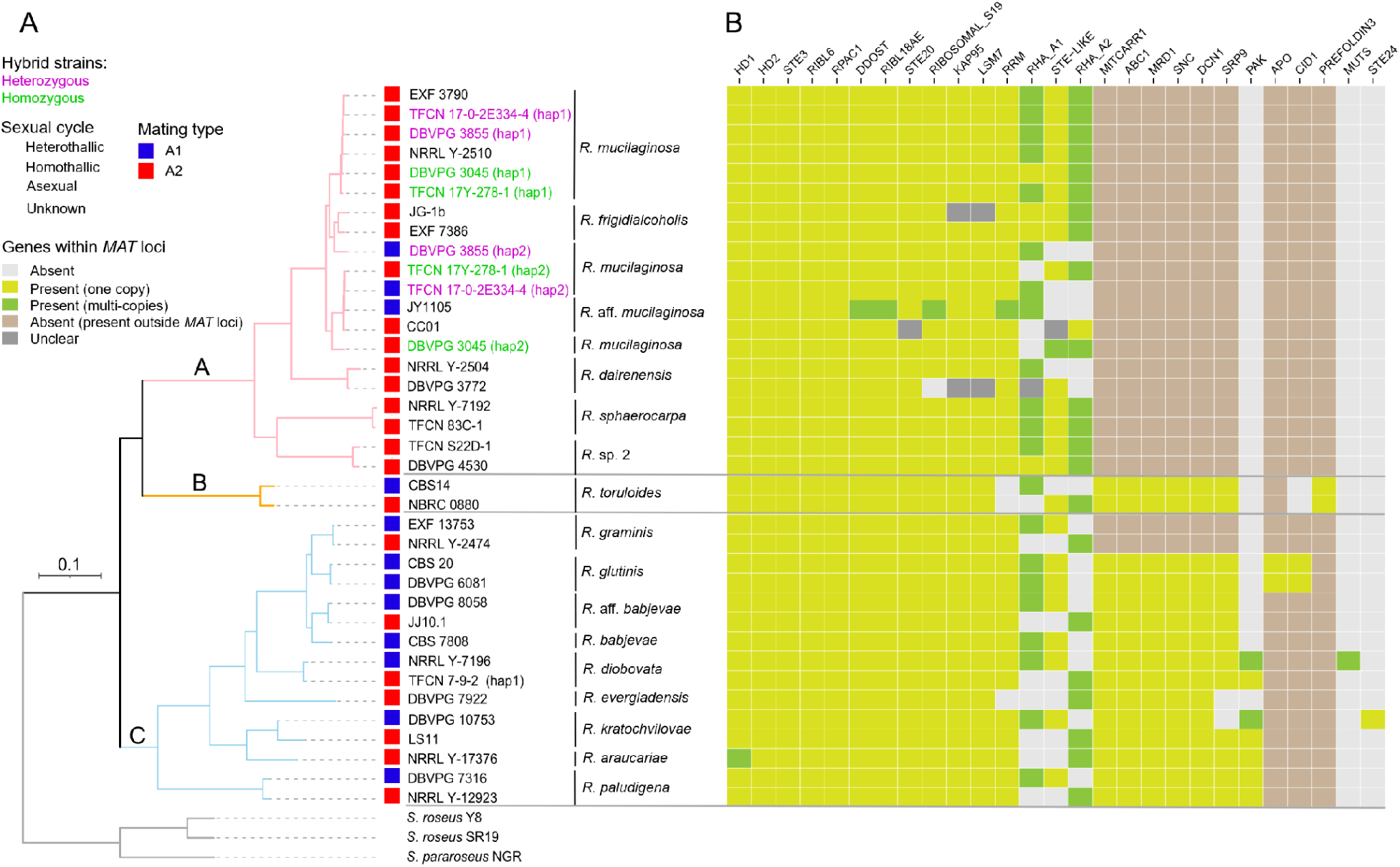
Phylogeny and gene composition of the *MAT* locus in *Rhodotorula*. **(A)** Phylogenomic species tree of *Rhodotorula* including both mating type per species, as well as phased subgenomes (hap1 and hap2) from heterozygous (*A1*+*A2*; purple) and homozygous (*A2*+*A2*; green) *R. mucilaginosa* hybrids; **(B)** Heatmap of gene presence-absence, and copy number within inferred *MAT* loci, ordered by presence frequency across species (most conserved on the left). ‘Absent’ indicates genes missing from the genome or localized outside the *MAT* locus. ‘Unclear’ indicates ambiguous locus assignment due to assembly fragmentation.

By contrast, comparisons between A1 and A2 *P/R* alleles revealed extensive structural divergence across the entire locus. For example, the A1 and A2 *P/R* alleles of *R. paludigena* differed by four inversions and rearranged gene modules (Fig. 3 and Supplementary text S1). Across species, *STE20* located near one boundary of the *P/R* locus in A2 strains, whereas it resided toward the center or opposite boundary in A1 strains. This positional shift may be caused by multiple overlapping inversions that collectively produce a translocation-like rearrangement. Together, the combination of broad within-type synteny and extensive between-type rearrangement indicates the long-term maintenance of structurally distinct, mating-type-specific haplotypes.

Multi-copy pheromone precursor *RHA* genes and transposable elements (TEs) were frequently associated with structurally variable regions of the *P/R* locus. *RHA* copies commonly flanked rearranged modules, whereas TEs were detected across nearly all *P/R* loci—with the sole exception of *Rhodotorula* sp. 1 DBVPG 10348, where an assembly gap within the locus likely hindered TE detection (Table S6). A large number of unclassified TE families were identified within the *P/R* locus and frequently occurred within *STE20* or accumulated near *RHA.A1* genes. Notably, Long Terminal Repeat (LTR) retrotransposons from the *Gypsy* and *Copia* families were also detected in low-gene-density intergenic regions of several A1 *P/R* loci, including those of *R*. aff. *mucilaginosa* JY1105, TFCN 17-0-2E334-4 hap2, and *R*. *kratochvilovae* DBVPG 10753. Their co-occurrence with structurally variable regions suggests *RHA* gene duplications and TE insertions as key drivers of *P/R* locus structural diversification.

The *HD* locus showed substantially less structural variation than the *P/R* locus. Across most species and mating types, it contained only the divergently transcribed *HD1* and *HD2* genes, with allelic differences largely confined to the N-terminal regions of their encoded proteins (Fig. S8D to F). Gene order and the surrounding flank architecture were highly conserved across species and mating types (Fig. 5). A notable exception was identified in *R. araucariae* NRRL Y-17376, whose *HD* locus contained an additional *HD1* copy together with an extra *MITCARR2* gene and a gene encoding a pectin lyase fold/virulence factor protein. Given the presence of a second *HD1* copy, we hypothesized that the accompanying pectin lyase gene originated from a heavily diverged *HD2* paralog. To test this, we compared its sequence against known *HD2* alleles; however, we detected no sequence homology, arguing against an *HD2* origin. Consequently, how these extra genes arose and whether they function within the *HD* locus remain unresolved.

**Figure 5.**
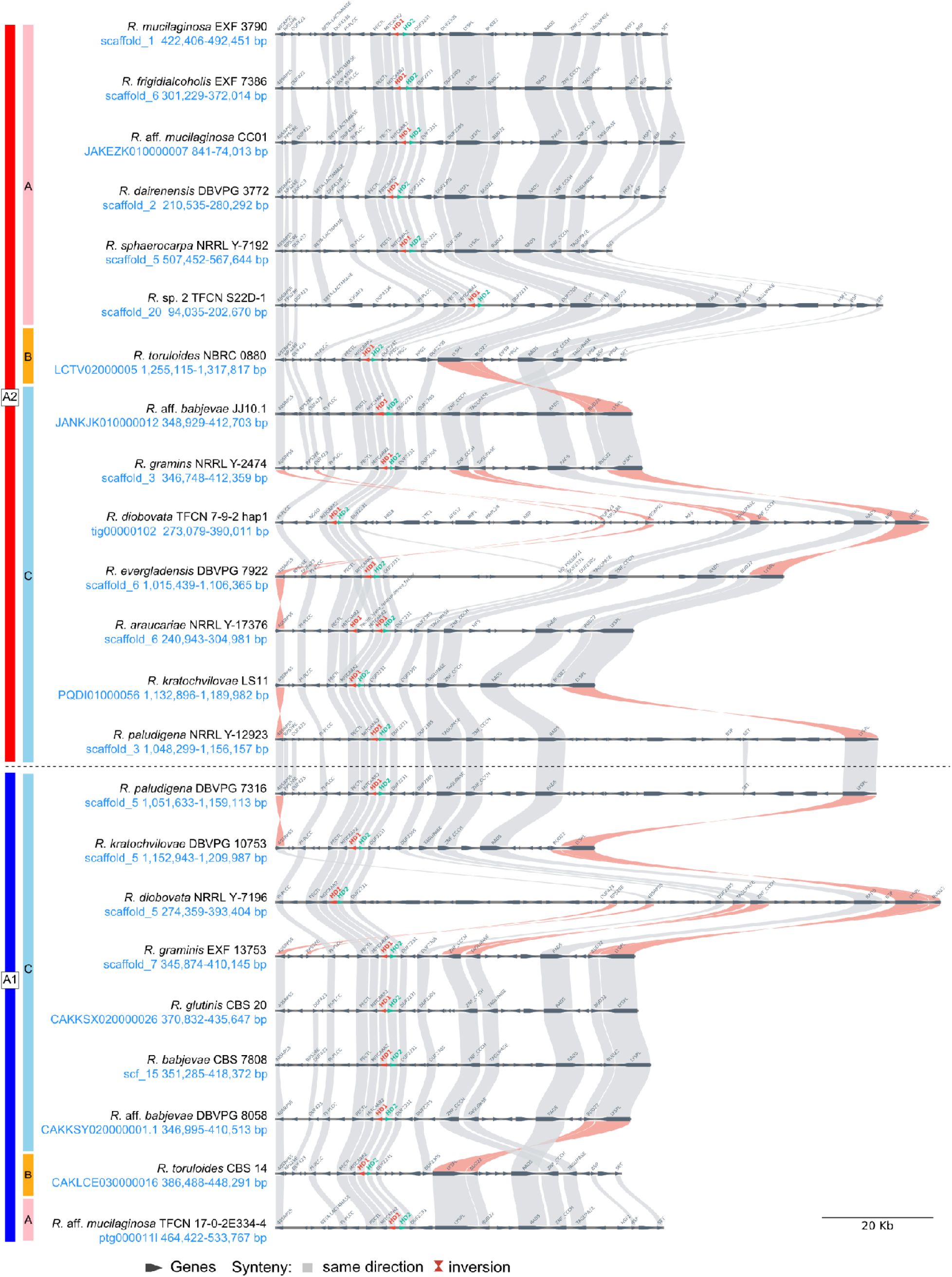
Synteny and gene organization of the *HD* locus across *Rhodotorula* species and mating types. Comparative synteny map reveals largely conserved locus architecture across species and mating types, with a notable *HD1* duplication in *R. araucariae*. A1 and A2 *MAT* alleles are labeled on the left. Canonical mating genes, *HD1* (red) and *HD2* (green), are highlighted. Homologous genes between adjacent strains are connected by colored shades, where gray indicates collinearity and red indicates inversion.

### Clade-specific mating-type distributions and genome-wide LD decay patterns

Phylogenetic analyses of genes within and flanking the *MAT* loci revealed that only the *STE3* gene showed a deep trans-specific polymorphism (Fig. 6A and Fig. S9), consistent with previous analyses of fungal pheromone receptors (Fraser et al. 2004; Coelho et al. 2010; Branco et al. 2018). We used *STE3* alleles to assign mating identity of *Rhodotorula* strains within each species population. The distributions of mating types differed markedly among the three major clades (Fig. 6B). In Clade C, both *P/R* A1 and A2 were presented in most species represented by multiple strains. Sampling remained limited for *R. glutinis*, *R. babjevae*, *R. evergladensis*, and *R. araucariae*, precluding a comprehensive assessment of their natural population-level mating-type composition. Previous surveys identified only A2 strains of *R. graminis* and A1 strains of *R. glutinis*, whereas no comparable bias was detected in the *R. babjevae* species complex (Coelho et al. 2011). Clade B, corresponding to the *R. toruloides* species complex, also harbored both mating types, but A1 and A2 strains segregated into distinct phylogenetic subclades, with pairwise ANI values below 95% between representative lineages (Fig. S5B). In contrast, Clade A showed a pronounced mating-type skew: most species contained only A2 strains, whereas *R.* aff. *mucilaginosa* was strongly biased towards A1.

**Figure 6.**
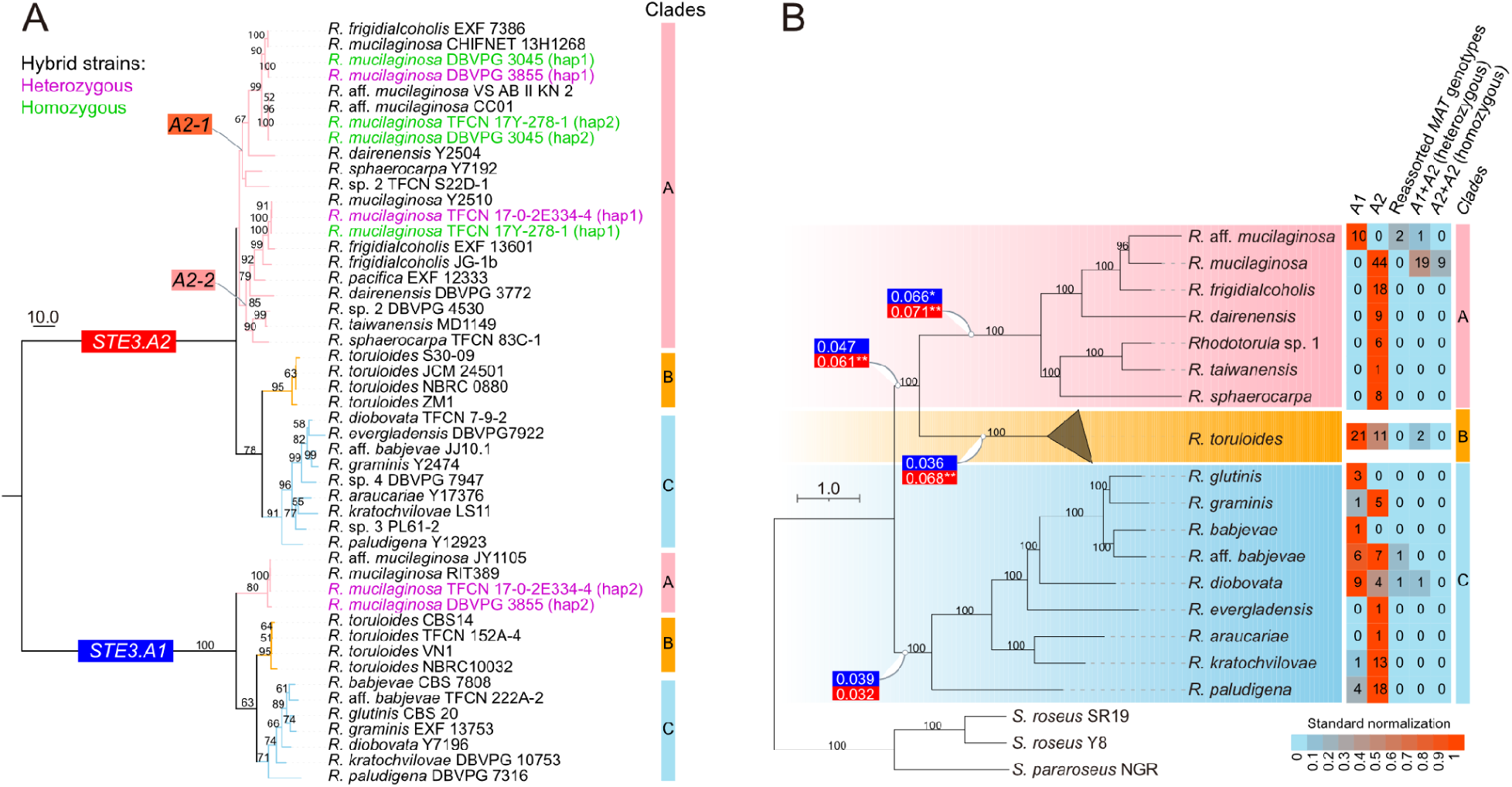
Mating-type allele diversity, evolution, and population distribution in *Rhodotorula*. **(A)** Maximum likelihood phylogeny of *STE3* showing alternate alleles *STE3.A1* and *STE3.A2*. Within clade A, *STE3.A2* splits into two divergent subclades (*A2-1* and *A2-2*), reflecting trans-specific polymorphism. Diploid hybrids are highlighted by heterozygous (A1+A2; purple) and homozygous (A2+A2; green); **(B)** *Rhodotorula* species tree (left) paired with a *MAT* genotype heatmap across 249 global strains (right). Major Clades A, B, and C are shaded and labeled. Branch labels indicate lineage-specific *STE3* evolutionary rates (ω = *d_N_*/*d_S_*) for *STE3.A1* (blue) and *STE3.A2* (red). Heatmap categories include *P*/*R* A1, *P*/*R* A2, reassorted *MAT* alleles, A1+A2 heterozygous diploids, and A2+A2 homozygous diploids, with color intensity showing row-normalized strain counts (0 = blue, min; 1 = red, max). Underlying data are available in Table S4 and S7.

To assess whether the A2-dominant *R. mucilaginosa* population retains signatures of historical recombination, we compared its genome-wide LD decay against that of the sexual species *R*. *diobovata*. The non-*MAT* genomic background in both species showed clear distance-dependent LD decay, with *r*^2^ declining rapidly over short physical distances before reaching a baseline plateau (Fig. S10). However, genome-wide LD decayed more gradually in *R. mucilaginosa*, with an estimated LD50 value of 20.21 kb, compared with 10.09 kb in *R*. *diobovata*. Mean *r*^2^ over the 0-50 kb interval was likewise higher in *R. mucilaginosa*, whereas LD values converge at the intermediate distances of 50-150 kb. These genome-wide patterns are consistent with historical recombination in both species, although recombination rates appear to have been higher in *R. diobovata*, driving its faster short-range decay.

### Deep *STE3.A2* divergence and mosaic A2 *P/R* architectures in Clade A

The A1 and A2 classes of *STE3* alleles showed contrasting patterns of sequence diversity. In Clade A, *STE3.A2* sequences exhibited pronounced diversification and segregated into two deeply divergent subclades, designated *A2-1* and *A2-2*, with representatives of both lineages co-occurring within multiple species (Fig. 6A). Branch-model analyses indicated overall purifying selection across both *STE3.A1* and *STE3.A2*, with *d_N_/d_S_* value below one across all tested branches (Fig. 6 and Table S7). However, evolutionary rates varied among lineages. For *STE3.A1*, clade A showed a significantly higher *d_N_/d_S_* than Clades B and C or their common ancestor branches (likelihood ratio test, LRT, *p* < 0.05) (Fig. 6B), whereas no rate difference was observed between Clades B and C and their common ancestor. For *STE3.A2*, the common ancestor of Clades A and B exhibited a significantly higher *d_N_/d_S_*than Clade C (*p* < 0.01), suggesting accelerated evolution prior to the divergence of Clade A and B. Since all branch *d_N_/d_S_* estimates remained below one, these elevated ratios were more consistent with relaxed purifying selection rather than positive selection. RELAX analyses supported this interpretation: *STE3.A1* showed evidence of relaxed selection in the common ancestor of Clades A and B (*k* = 0.34, *p* = 0), while *STE3.A2* showed stronger relaxation both in the ancestors of Clade A and B (*k* = 0.40, *p* = 0) and within Clade A (*k* = 0.12, *p* = 0) (Table S8). Branch-site-model analyses detected no evidence of positive selection for either *STE3* alleles (Table S9), with only a few sites assigned to positive-selection categories (Table S10). Together, these results demonstrate that the accelerated evolution of *STE3.A2* in Clade A was primarily driven by relaxation of purifying selection.

Sequence divergence of *STE3.A2* was coupled with structural changes in *P*/*R* locus organization. In Clade A, strains carrying divergent *STE3.A2* alleles exhibited altered local synteny across the *P*/*R* locus (Fig. S8A). In addition, A2 strains in Clade A frequently carried *RHA.A1*, a pheromone gene typically restricted to A1 backgrounds (Fig. S8A and Table S11). These features define a recurrent, mosaic A2 *P*/*R* architecture characterized by divergent *STE3.A2* receptor alleles, the unexpected retention of A1-associated pheromone genes, and syntenic rearrangements. Whether these mosaic configurations alter pheromone recognition or impact mating compatibility remain to be tested.

### Subgenome-resolved analyses reveal A1/A2 and A2/A2 hybrid *MAT* configurations

Genomes carrying a single *STE3* allele alongside single copies of *HD1* and *HD2* were classified as haploid, with genome sizes ranging from 17.4 to 23.7 Mb. In contrast, genomes larger than 35 Mb were inferred to be diploid or hybrid. We identified 35 such diploid strains, including 30 *R. mucilaginosa*, two *R. toruloides*, one *R. frigidialcoholis*, one *R.* aff. *mucilaginosa*, and one *R. diobovata*, with sizes ranging from 36.6 to 45.5 Mb (Fig. 2). These strains spanned diverse habitats: 21 strains were isolated from tidal flat zones, five from plants, four from clinical sources, two from food, one from soil, one from conjugation, and one from an unknown source. At the mating-type loci, 23 of these strains harbored both *STE3.A1* and *STE3.A2* alleles, whereas nine strains carried two copies of *STE3.A2*. The remaining three strains—two *R. mucilaginosa* and one *R. frigidialcoholis*—carried only a single detectable *STE3.A2* allele despite retaining two copies of *HD1* and *HD2*, suggesting possible loss of heterozygosity at the *P/R* locus.

To determine whether these diploid genomes arose through whole-genome duplication or interspecies hybridization, we generated PacBio HiFi assembled for two A1/A2 heterozygotes *R. mucilaginosa* strains (DBVPG 3855 and TFCN 17-0-2E334-4) and two A2/A2 homozygotes strain (DBVPG 3045 and TFCN 17Y-278-1). Subgenome-resolved phylogenetic analyses revealed that the hap1 subgenomes of all four diploids were closely related to *R. mucilaginosa* Y-2510, whereas their hap2 subgenomes showed different affinities. Specifically, the hap2 subgenomes of TFCN 17-0-2E334-4 and TFCN 17Y-278-1 were closely related to *R.* aff. *mucilaginosa* JY1105, whereas DBVPG 3045 formed a weakly supported sister group to this subclade, and DBVPG 3855 clustered with *R. frigidialcoholis* JG-1b (Fig. 7A). Homeologous subgenomes within each diploid share less than 92% ANI—well below the commonly used 95% species threshold—suggeting their interspecific hybrid origins (Fig. 7B). All hap1 subgenomes shared more than 98% ANI with *R. mucilaginosa*, suggesting this species as one parental lineage. Similarly, the hap2 subgenomes of TFCN 17-0-2E334-4 and TFCN 17Y-278-1 shared more than 98% ANI with *R.* aff. *mucilaginosa* JY1105, indicating that they are conspecific with this lineage. By contrast, the hap2 subgenome of DBVPG 3045 and DBVPG 3855 showed below 95% ANI to their closest relatives, suggesting origins from more divergent or unsampled parental lineages.

**Figure 7.**
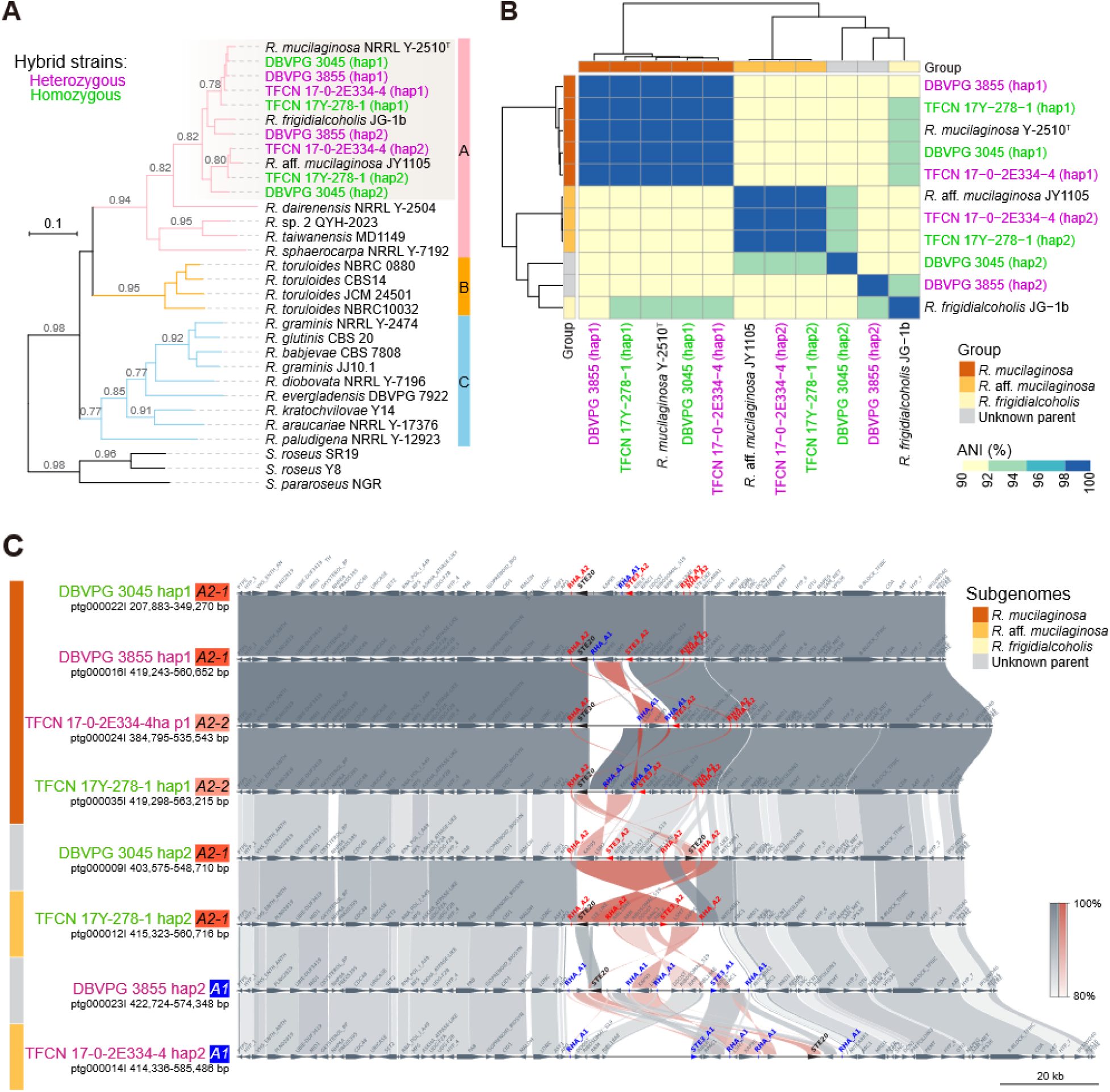
Phylogenetic placement, genome divergence, and the *P/R* locus architecture in *R. mucilaginosa* hybrid strains. **(A)** Species tree of *Rhodotorula* showing phased subgenomes (hap1 and hap2) from heterozygous (*A1*+*A2*; purple) and homozygous (*A2*+*A2*; green) *R. mucilaginosa* hybrids; **(B)** ANI heatmap comparing hybrid subgenomes and their close relatives (*R. mucilaginosa*, *R.* aff*. mucilaginosa*, and *R. frigidialcoholis*); **(C)** Synteny comparisons of the *P/R* locus across heterozygous and homozygous hybrid subgenomes. Colored blocks on the left denote inferred parental origin.

Homeologous *P/R* loci between subgenomes also exhibited substantial structural divergence (Fig. 7C and Fig. S11). In the A1/A2 hybrid DBVPG 3855, the *STE3***–***RIBL6*–*RPAC1* module was relocated from the left flank of *DDOST* to the right of *RIBL18AE*. In the A1/A2 hybrid TFCN 17-0-2E334-4, three structural rearrangements involved *STE20*, *LSM7***–***KAP95*and *DDOST*–*RIBOSOMAL*_*S19*–*RRM*–*RIBL18AE*. Structural differences between subgenomes were also characterized in the A2/A2 hybrids: DBVPG 3045 differed between subgenomes by inversion of *STE20*, whereas TFCN 17Y-278-1 exhibited a larger inversion spanning the *STE–LIKE*–*RIBL18AE*–*RRM*–*RIBOSOMAL_S19*–*DDOST*–*RPAC1*–*RIBL6*–*STE3* module, together with a relocation of *KAP95*–*LSM7* module.

The two *STE3.A2* copies in the A2/A2 hybrids were also divergent. In TFCN 17Y-278-1, the *STE3.A2* alleles from hap1 and hap2 belonged to *A2-2* and *A2-1* subclades, respectively, clustering with sequences from *R. mucilaginosa* CHIFNET 13H1268 and Y-2510 (Fig. 6A). In DBVPG 3045, both *STE3.A2* alleles belonged to subclade *A2-1* but shared less than 90% sequence identity—a level of divergence higher than typical within-subclade comparison (Fig. S12). This divergence likely reflects their origin from different parental species and was accompanied by structural differences across their corresponding *P/R* loci (Fig. 7C). Notably, the *P/R* locus in the hap1 subgenomes of these A2/A2 hybrids retained at least one copy of the *RHA.A1* gene. These results support interspecific hybrid origins for the examined diploid strains: A1/A2 diploids combined opposite mating-type haplotypes, whereas the A2/A2 diploids integrated two distinct A2 haplotypes from separate parental species within the *R. mucilaginosa* species complex.

### A2-associated genomic differentiation extends beyond the *MAT* loci

We next examined whether A2 dominance in Clade A was associated with broader genomic differentiation outside the *MAT* regions. Orthogroup presence-absence analysis incorporating representative *Rhodotorula* species and haplotype-resolved subgenomes from *R*. *mucilaginosa* hybrids identified an UpSet plot core set: 49 orthogroups were specifically shared by the A2-type haploid genome *R. mucilaginosa* Y-2510 and the A2-type hap1 subgenomes of four hybrid strains (DBVPG 3855, DBVPG 3045, TFCN 17-0-2E334-4, and TFCN 17Y-278-1), but were absent from the corresponding A1-type haplotypes and other *Rhodotorula* species (Fig. S13A). Although most of the 49 orthogroups encoded hypothetical proteins, analysis of public RNA-seq datasets confirmed active transcription for the majority of these genes under at least one condition, with a subset showing moderate to high expression across multiple conditions (Fig. S13B). Furthermore, predicted proteins within this set had length distributions comparable to standard annotated proteins, with most exceeding 100 amino acids (Fig. S13C, D). These observations support that this A2-associated gene set represents functional loci rather than gene prediction artifacts.

Among the 49 A2-associated orthogroups, 11 formed a physically clustered genomic block. Both gene order and orientation within this block were conserved across A2-type *R. mucilaginosa* hap1 subgenomes; in contrast, the corresponding regions in A1-type subgenomes and related Clade A species completely lacked this gene cluster (Fig. S14). Its A2-restricted distribution, physical clustering, and conserved synteny identify this region as a candidate A2-associated accessory genomic island. Although most genes within this block encoded hypothetical proteins, several encoded recognizable domains, including a C2H2-type zinc finger, a Cryptic loci regulator 2, and fungal fatty acid synthase subunit. Genomic differentiation associated with the A2 background in Clade A therefore extends beyond the *MAT* loci to include a conserved, lineage-specific accessory gene island.

## Discussion

Comparative genomic analyses position *Rhodotorula* as a useful genus-level system for investigating mating-system evolution in basidiomycetous fungi. Across the genus, we uncovered a spectrum of reproductive architectures—from Clade C lineages that retain both mating types and the potential for canonical heterothallism, to lineage-structured mating-type segregation in Clade B, and ultimately to A2-dominated populations in Clade A. This diversity was supported by multiple layers of genomic evidence, including constrained *P*/*R*–*HD* allele combinations, clade-specific mating-type distributions, allele- and lineage-specific remodeling of the plastic *P*/*R* locus, and deep trans-specific polymorphism coupled with heterogeneous evolutionary dynamics at *STE3*. Within A2-dominant Clade A, this evolutionary trajectory is marked by deep *STE3.A2* divergence, local *P/R* locus rearrangement, the discovery of A2/A2 hybrid genomes, and an A2-associated accessory gene island beyond the *MAT* loci. Furthermore, Genome-wide LD decay in *R*. *mucilaginosa* indicates historical recombination despite the absence of an observed sexual cycle. These findings reveal clade-level differences in mating-type availability, *P*/*R*–*HD* allele combinations, and the extent of *P*/*R* locus remodeling across *Rhodotorula*. In Clade A, strongly skewed mating-type distributions, sequence and structural divergence among A2 *P*/*R* haplotypes, A2/A2 hybrid, and genome-wide evidence of historical recombination collectively suggests that alternative reproductive routes may persist despite the absence of an observed sexual cycle.

### Taxogenomic discordance reveals recent divergence and incomplete reproductive isolation in *Rhodotorula* species complexes

Interpreting reproductive diversity in *Rhodotorula* requires a taxogenomic framework because species boundaries, genome divergence, and mating compatibility frequently decouple in fungi, particularly in yeasts with hidden diversity (Gostinčar 2020; Libkind et al. 2020). In this study, we resolved three major clades and identified several closely related species complexes, including the Clade A group containing *R. mucilaginosa*, *R.* aff. *mucilaginosa*, and *R. frigidialcoholis*, and the Clade B *R. toruloides* species complex. Intermediate ANI values, unstable phylogenetic placement, and local chromosomal rearrangements across these groups are consistent with recent divergence, incomplete lineage sorting, ongoing or historical gene flow, and hybridization (James et al. 2020; Libkind et al. 2020). Accounting for this taxogenomic complexity is critical, because mating-type variation may reflect polymorphism within a species, lineage sorting among recently diverged taxa, or introgression across incompletely isolated lineages.

The *R. toruloides* complex in Clade B provides a classic example of this discordance. Strains carrying opposite mating types formed distinct phylogenetic subclades and exhibited 90–95% ANI between representative pairs (e.g., ATCC 10788 and ATCC 10657, or CBS 14 and NBRC 0880), consistent with previous observations (Hu and Ji 2016; Brink et al. 2023). Genome-scale interchromosomal rearrangements, including gene relocations and inversions, further contribute to structural divergence among these closely related species or mating-compatible strains, as exemplified by NBRC 0880 and CBS 14 (Fig. S4C). Nevertheless, genome-wide sequence divergence has not driven complete reproductive isolation: certain strains either share a common diploid ancestor (such as ATCC 90781; Hu and Ji 2016), or remain sexually compatible, as evidenced by their ability to produce viable offspring with the type strain CBS 6016 (synonym PYCC 5015) (Banno 1967). Thus, *MAT*-locus differentiation and structural divergence appear to have preceded the establishment of complete reproductive isolation. More broadly, because reproductive isolation in fungi may arise from changes at a limited number of key loci, fixed genomic thresholds for species delimitation should be applied in a lineage-specific context rather than used as universal criteria (Hart et al. 2025).

### Constrained tetrapolarity and differential *MAT*-locus plasticity shape the *Rhodotorula* mating system

Chromosome-scale assemblies show that *Rhodotorula* retains physically tetrapolar *MAT* loci, with the *P*/*R* and *HD* loci residing on separate chromosomes. This arrangement distinguishes it from pseudo-bipolar systems described in red yeasts such as *Sporobolomyces* and in some *Malassezia* species, where both loci occupy the same chromosome with limited recombination (Coelho et al. 2010; Coelho et al. 2023). This physical separation in *Rhodotorula*, however, does not lead to free allelic reassortment: most *HD* alleles remain strictly associated with a single *P/R* background. We interpret *Rhodotorula* as exhibiting constrained tetrapolarity, in which physically unlinked *P*/*R* and *HD* loci segregate in restricted allele combinations.

Several evolutionary and genetic mechanisms could generate this pattern. Transmission distortion or differential survival of meiotic products could enforce the co-segregation of parental allele combinations without requiring physical linkage (Lindholm et al. 2016). For example, in the tetrapolar basidiomycete *Leucosporidium scottii*, reassorted *MAT* allele combinations are absent from natural populations but readily recovered among laboratory-generated F1 progeny without reduced fitness, suggesting that parental *MAT* combinations are favored in nature (Maia et al. 2015). Alternatively, centromere-associated segregation biases, as proposed in the predominantly selfing fungi *Microbotryum lagerheimii* and *M. saponariae* (Carpentier et al. 2019), or *MAT*-linked haplo-lethal alleles, as reported in *Ustilago* and *M. violaceum* (Hood and Antonovics 2000; Rabe et al. 2016), could actively restrict meiotic survival to specific *MAT* combinations. Repeated transitions from tetrapolar to bipolar organization via physical locus linkage represent one known route to limit *MAT* reassortment, an adaptive strategy often favored in selfing or same-sex-like pathogenic species to maximize gamete compatibility and stabilize genetic content (Nieuwenhuis et al. 2013; Coelho et al. 2023). *Rhodotorula* expands the known spectrum of fungal mating systems by showing that restricted *MAT* inheritance can evolve and persist independent of physical linkage between the *P*/*R* and *HD* loci.

The *P/R* locus exhibits high syntenic conservation within each mating type but extensive structural divergence between A1 and A2 alleles, indicating long-term maintenance of mating-type-specific haplotypes. Similar tightly linked *MAT* regions have evolved independently in *Cryptococcus*, *Trichosporonales*, and anther-smut fungi (Branco et al. 2018; Sun et al. 2019; Coelho et al. 2025). In *Rhodotorula*, the conserved *P*/*R* core includes *STE3*, *STE20*, and a *STE12*-like gene, which encode key components of the pheromone-signaling MAPK cascade (Fraser et al. 2004). The locus also contains genes lacking known mating functions, including ribosomal proteins (*RIBL6*, *RIBOSOMAL_S19*, *RIBL18AE*), RNA recognition and splicing factor (*RRM*, *LSM7*), karyopherin beta (*KAP95*), dolichyl-diphosphooligosaccharide transferase (*DDOST*), and RNA polymerase subunits (*RPAC1*), in the related taxon *L. scottii* (Maia et al. 2015). Co-selection on linked mating and cellular functions, together with combination suppression between alternative *P*/*R* alleles, likely maintains the structural integrity of these locus haplotypes.

The genomic extent of the *P*/*R* locus varies across *Rhodotorula* lineages. Clade A species and *R*. *graminis* retain a relatively compact locus configuration, whereas species in Clade B and C possess a broader right-flanking region encompassing up to seven additional genes that reside outside the locus defined in Clade A. Because these genes also present within or adjacent to the *P*/*R* locus of the early-diverging relative *L. scottii* (Maia et al. 2015), their phylogenetic distribution suggests an ancestral association with the *P*/*R* region, followed by lineage-specific rearrangements or boundary shifts.

Multiple copies of pheromone gene *RHA* with inverted orientations represent another recurrent feature of the *Rhodotorula P/R* locus, mirroring patterns observed in *Cryptococcus* and other *Microbotryomycetes* yeasts (Maia et al. 2015; Huang et al. 2024; Coelho et al. 2025; Lucotte et al. 2025). Increased *RHA* copy number may enhance partner recognition, as proposed in the dosage-based model for *C*. *deneoformans* (Shen et al. 2002). Inverted *RHA* gene pairs may also drive structural plasticity by serving as potential sites for recombination and inversion loops, thereby expanding recombination-suppressed regions within the *P/R* locus (Coelho et al. 2025). In Clade C, broader *P*/*R* locus architectures are characterized with wider physical spacing among *RHA* gene pairs, a greater number of intervening genes, extensive structural rearrangements, and increased TE site density near *RHA* genes. These associations highlight pheromone-gene duplication and TE accumulation as potential drivers to *P*/*R*-locus plasticity.

In contrast, the *HD* locus is compact and largely conserved across species and mating types. In basidiomycetes, the *HD1*/*HD*2 complex regulates post-fusion sexual development, although functional requirements for these genes vary across reproductive modes. In the related species *L. scottii* and *Sporobolomyces salmonicolor*, successful mating requires compatible alleles at both *P*/*R* and *HD* loci (Coelho et al. 2010; Sun et al. 2019). In *Cryptococcus*, *HD* genes *Sxi1α* and *Sxi2a* can be dispensable for hyphal morphogenesis and sporulation during unisexual reproduction, leaving vegetative growth and virulence-related traits unaffected (Lin et al. 2010; Huang et al. 2025). The conserved *HD* architecture in *Rhodotorula*, together with its non-random co-occurrence with *P*/*R* alleles could reflect constraints on particular allele combinations, although reduced or lost mating-compatibility functions in some *HD* haplotypes cannot be excluded (Lucotte et al. 2025). Taken together, the contrasting evolutionary dynamics of these two *MAT* components highlight the structural plasticity of the *P*/*R* locus, whereas the *HD* locus maintains a conserved, stable genomic core.

### Mating-type asymmetry place Clade A in a distinct reproductive context

Mating-type distributions differed markedly among the three major clades. In Clade C, both compatible mating types were present in most sampled species, consistent with the retention of heterothallic potential and laboratory evidence for sexual reproduction (Sampaio 2011). Clade B also contained both mating types, but A1 and A2 strains were more strongly partitioned into distinct phylogenetic subclades. Clade A showed the greatest asymmetry, with most sampled species represented predominantly by A2 strains. Within the *R*. *mucilaginosa* species complex, sampled *R*. *mucilaginosa* and *R*. *frigidialcoholis* strains were A2, whereas *R*. aff. *mucilaginosa* strains were exclusively A1. This lineage-partioned pattern points to the presence of both mating types in their common ancestor, followed by lineage sorting, demographic expansion, founder effects or ecological differentiation.

Mating-type biases are common in fungi and frequently reflect population history, niche specialization, or selection on mating-type-linked physiological traits. In *Cryptococcus neoformans* var. *grubii*, *MAT*a strains are largely restricted to the mopane-associated sub-Saharan African VNB lineage (Litvintseva and Mitchell 2012). Anthropophilic dermatophytes frequently comprise single mating-type lineages and lack a known sexual cycle, in contrast to their sexually active geophilic relatives (Metin and Heitman 2017). Soil and corn populations of *Aspergillus flavus* are enriched for sclerotium-producing S-type and prolific conidium-producing L-type morphotypes, respectively, which exhibit distinct maternal and paternal reproductive roles during sexual crossing (Sweany et al. 2024). Beyond cell-type specification, *MAT* genes are embedded within broader developmental and physiological regulatory networks that influence carbon utilization, stress tolerance, and virulence (Zhang et al. 2024). In *Saccharomyces eubayanus*, *MAT*-dependent haploid-like transcriptional programs interact with carbon metabolism networks to confer condition-dependent fitness advantages during adaptation to maltose utilization (Crandall et al. 2023). Therefore, A2 dominance in Clade A does not necessarily signify a degenerated *MAT* system; rather, it likely represents a shift in the ecological niche and evolutionary context that reduces dependence on canonical A1/A2 encounters.

The expansion of A2 lineages may also involve genomic differentiation beyond the *MAT* loci. In this study, we discovered a conserved, physically linked accessory genomic island exclusive to A2-type *R. mucilaginosa* genomes as well as subgenomes isolated from hybrids. Several genes in this island encoded proteins containing domains such as a zinc finger, a Clr2-like chromatin regulator, a fatty acid synthase subunit. This tight physical linkage hints at potential functional coupling between transcriptional or epigenetic regulation, and lipid metabolism (Rossi et al. 2009; Arcangioli and Gangloff 2023; Son et al. 2025). While these predicted functions do not yet establish an integrated metabolic pathway or a direct mechanism for A2 dominance, this lineage-specific genomic block provides candidate loci for testing whether A2 expansion was accompanied by broader physiological adaptations. Similar accessory genomic islands in other microbial systems have been shown to confer distinct ecological benefits, such as enhanced stress tolerance, virulence, or nutrient utilization (Yuan et al. 2021).

Regardless of whether this genomic differentiation contributed to A2 lineage success, the resulting mating-type imbalance inevitably restricts canonical partner availability. Genome-wide LD decay in the A2-dominant *R*. *mucilaginosa* population nevertheless reveals a sign of historical recombination, arguing against strict clonality. Although its estimated LD50 of 20,206 bp is longer than that of the sexual species *R. diobovata* (10,095 bp) and the range reported for outcrossing *Aspergillus flavus* populations (1,000-12,300 bp; Drott et al. 2020), it is substantially shorter than the 162,100 bp observed in the predominantly clonal *Candida albicans* (Nieuwenhuis and James 2016) and and closely matches to the 20,010 bp reported for a sexual population of *Schizosaccharomyces pombe* (Jeffares et al. 2015). These cross-species comparisons should be interpreted qualitatively rather than as precise quantifications of sexual frequency, given that LD decay rate is sensitive to population structure, demographic history, sampling bias, and filtering criteria (Nieuwenhuis and James 2016).

Overall, the genomic decay profile of *R*. *mucilaginosa* aligns with a reproductive history involving both clonal propagation and recombination, though it does not resolve the specific frequency or mechanism of reproductive exchange within an A2-dominant population. Meiotic sex, unisexual or parasexual processes, or rare outcrossing events could each generate this recombination signal, as documented in mating-type-biased populations of *C*. *neoformans*, *C. gattii*, *Aspergillus*, and *Penicillium* species (Lin et al. 2005; Dyer and O’Gorman 2011; Yadav et al. 2023; Coelho et al. 2025).

### *MAT* remodeling and hybrid configurations suggest altered mating compatibility in Clade A

The evolution of *STE3.A2* highlights a potential mechanism by which mating recognition may have evolved in A2-dominant lineages. Persistent mating-type skews can weaken the balancing selection that typically preserves alternative mating-type alleles (Peris et al. 2022). In Clade A, *STE3.A2* sequences formed two deeply divergent trans-specific subclades, indicating that the polymorphism predates diversification of the sampled species. Their accelerated evolutionary rate compared to *STE3.A1*, together with the RELAX results, is more consistent with relaxed purifying selection than positive selection. This reduced selective constraint may have permitted sequence and functional divergence in the receptor, potentially altering pheromone recognition specificity or downstream signal transduction across A2 subtypes.

Mosaic *P/R* locus configurations provide a second mechanism for altered compatibility. In Clade A, the A2-associated receptor *STE3*.*A2* frequently co-occurs with *RHA.A1*, a pheromone precursor otherwise characteristic of the A1 background. These non-canonical arrangements departed from standard A1 and A2 haplotypes and likely arose through rare recombination or introgression events. Conserved local synteny and proximity of transposable elements to *RHA* genes could provide short homologous tracts that facilitate ectopic recombination, though the precise exchange events remain to be demonstrated. The incorporation of an opposite-type pheromone component into an A2 background could alter pheromone-receptor pairings and potentially generate novel self- or inter-lineage compatible haplotypes in A2-dominant populations. A relevant precedent occurs during α-α unisexual reproduction in *C. neoformans*, where rare crossovers between collinear *MAT*α haplotypes generate mosaic *MAT* configurations, despite overall recombination suppression within the locus (Sun et al. 2014). In *Rhodotorula*, subsequent inversions (e.g., *LSM7*-*KAP95* in *R. mucilaginosa*, and *STE20* alongside the multi-gene block containing *STE3* in *R. frigidialcoholis*) could stabilize these divergent A2 *P*/*R* configurations by preventing further homologous recombination.

Comparative analyses across budding yeasts provide a broader evolutionary context for this model, where repeated transitions from heterothallism to homothallism have occurred through the acquisition or relocation of opposite-type *MAT* determinants (Krassowski et al. 2019). While these specific reproductive transitions have not yet been demonstrated in *Rhodotorula*, they establish a clear evolutionary link between *MAT*-locus remodeling and shifts in reproductive strategy. Sequence divergence, mosaic gene combinations, and local structural rearrangements at the Clade A *P/R* locus may therefore serve as genomic substrates for altered mating compatibility, facilitating a transition toward reproductive strategies that bypass dependence on canonical A1/A2 partner availability.

In fungi, hybridization serves as a critical driver of speciation, evolution, and adaptation (Takashima et al. 2018; Steenwyk et al. 2020; Theelen et al. 2022). Within the *R*. *mucilaginosa* species complex, hybrid genomes record an additional outcome of mating-system reconfiguration. Diploid strains constituted over 40% of the sampled *R. mucilaginosa* population and comprised both canonical A1/A2 and non-canonical A2/A2 genotypes. Subgenome-resolved phylogenomic analyses identified *R. mucilaginosa* as one parent across four representative hybrids, while the opposing subgenomes aligned with *R.* aff. *mucilaginosa*, *R. frigidialcoholis*, or uncharacterized close lineages. The presence of A1/A2 hybrids indicates that reproductive isolation remains incomplete within the species complex despite genome-wide sequence divergence and lineage-partitioned mating types. Furthermore, the recovery of divergent second subgenomes points to hidden diversity or geographic restriction among compatible donor strains.

The discovery of A2/A2 hybrids directly informs our understanding of compatibility reconfiguration in Clade A. These strains reflect the combination of two distinct A2 parental lineages rather than whole-genome duplication. Moreover, one of these A2 *P/R* subgenomes contains *RHA.A1*, reinforcing the connection to mosaic *P/R* locus remodeling described above and suggesting that altered pheromone signaling may permit same-type mating. Although A2/A2 diploids provide compelling evidence for such mating strategy, alternative mechanisms—such as post-hybridization loss of heterozygosity following an initial A1/A2 hybridization—cannot be ruled out. Similar same-type hybrid configurations are documented in *Cryptococcus*, where aADα and αAAα hybrids originate via same-sex mating between *C. neoformans* and *C. deneoformans*—two lineages capable of cell fusion but constrained by post-zygotic barriers (Lin et al. 2007, 2009). Direct crossing assays, progeny viability analyses, and functional characterization of *STE3* and the *P*/*R* alleles will be required to determine whether *STE3.A2* divergence and mosaic *P*/*R* remodeling alter pheromone specificity, and whether the A2/A2 hybrids arose through same-sex mating or other noncanonical processes.

### Conclusion: a comparative framework for reproductive transitions in *Rhodotorula*

Across *Rhodotorula*, reproductive variation spans a broad spectrum—from lineages retaining balanced mating types and canonical heterothallic potential, through phylogenetic partitioning of mating types, to pronounced A2 dominance. The persistence of physically tetrapolar *MAT* loci, non-random *P*/*R*–*HD* allele associations, and genome-wide signatures of historical recombination demonstrates that the lack of an observed sexual cycle does not reflect a loss of genomic capacity for sex. In Clade A, deep *STE3.A2* divergence, mosaic *P*/*R* configurations, structural remodeling of the *P*/*R* locus, and the emergence of A2/A2 hybrids point toward a reduced dependence on canonical A1/A2 compatibility. Ultimately, these findings position *Rhodotorula* as a powerful model system for investigating how partner scarcity, constrained *MAT* assortment, and structural genome remodeling interact to reshape reproductive strategies in fungi.

## Materials and methods

### Isolate collection and dataset

We analyzed two datasets to resolve the taxogenomics, population structure, and mating-type variation within *Rhodotorula*. The first, designed for comparative genomic analyses, comprised 20 representative strains, including 14 described *Rhodotorula* species, three potential novel species, and three strains from the *R. toruloides* complex. The second dataset was used for broad-scale taxogenomic assessment, species delimitation, and mating-type distribution analysis. It comprised 249 isolates representing 15 described *Rhodotorula* species collected globally, combining 81 previously published strains from NCBI Sequence Read Archive (SRA) and GenBank databases with 168 newly sequenced strains. Newly sequenced isolates were obtained from the Industrial Yeasts Collection DBVPG, the Microbial Culture Collection Ex (University of Ljubljana, Slovenia), and tidal zones across China. Comprehensive strain metadata and database accession numbers for both datasets are detailed in Table S1 and S2.

### Culture growth conditions and DNA extraction

All strains sequenced in this project were purified via single-colony isolation on streak plates and stored in 25% glycerol at -80°C. For routine growth, cultures were maintained on yeast extract peptone dextrose medium (YPD, 1% yeast extract, 2% peptone, 2% glucose, 2% agar) at 25°C for three to five days. High-molecular-weight genomic DNA was extracted from yeast cells scraped from solid YPD medium using a modified cetyltrimethylammonium bromide (CTAB) protocol (Carter-House et al. 2020). To optimize extraction for *Rhodotorula*, two key modifications were introduced: (1) sequential wash steps using Phenol:Chloroform:Isoamyl alcohol (PCI) followed by Chloroform:Isoamyl alcohol (CI) to thoroughly remove cellular proteins, polysaccharides, lipids, and phenolic compounds; and (2) an extended one-hour DNA precipitation at -20°C following isopropanol addition. Purified DNA was dissolved in 60 µl of TE buffer and quantified using a Qubit 4 Fluorometer (Invitrogen, Thermo Fisher Scientific, Waltham, MA, USA) with the Qubit dsDNA Broad Range Assay kit. DNA purity was assessed by A260/A280 and A260/A230 absorbance ratios using a Nanodrop 2000c Spectrophotometer (Thermo Fisher Scientific, Waltham, MA, USA).

### Library preparation and sequencing

Illumina paired-end (PE) libraries for all newly collected strains in the second dataset were prepared using the purePlexTM DNA Library Preparation kit and sequenced on an Illumina NovaSeq 6000 platform (2 × 150 cycles) through the California Institute for Quantitative Biosciences at the University of California, Berkeley. Additionally, Oxford Nanopore Technologies (ONT) long-read sequencing was performed for eight selected strains. These included type strains for seven *Rhodotorula* species—*R. araucariae* Y-17376, *R. dairenensis* Y-2504, *R. diobovata* Y-7196, *R. graminis* Y-2474, *R. mucilaginosa* Y-2510, *R. paludigena* Y-12923, and *R. sphaerocarpa* Y-7192—alongside *R. evergladiensis* DBVPG 7922. For nanopore sequencing, barcoded libraries were prepared using the NBD104 barcoding expansion and LSK109 ligation sequencing kits according to the manufacturer’s protocols. Multiplexed libraries were loaded onto a MinION R9.4.1 flow cell and run on a MinION device for up to 48 hours. Raw signal data were basecalled and demultiplexed using Guppy 6.4.8. Finally, four diploid *R. mucilaginosa* isolates (DBVPG 3045, DBVPG 3855, TFCN 17-0-2E334-4, TFCN 17Y-278-1) were sequenced on the PacBio Revio system using HiFi mode (Annoroad Gene Technology Co. Ltd, Beijing, China). PacBio HiFi BAM files were converted to FASTA format using the bam2fasta for downstream analysis.

### Genome assembly and annotation

Illumina short-read data were assembled de novo using the Automatic Assembly For The Fungi (AAFTF) pipeline v. 0.5.0 (Palmer and Stajich 2026). Raw reads were first quality-filtered, trimmed, and cleaned using fastp (Chen et al. 2018) and bbduk (Bushnell 2014). *De novo* assembly was performed by SPAdes v3.15.5 (Bankevich et al. 2012; Prjibelski et al. 2020). Assembled contigs were screened for adaptor and vector contamination against the GenBank database using BLASTN (AAFTF *fcs* and *vecscreen*). Microbial contamination was removed using sourmash 4.6.1 (AAFTF sourpurge; Brown and Irber 2016), and duplicate contigs were eliminated with minimap2 2.24 (AAFTF rdump; Li 2018). Finally, consensus sequences were polished using Pilon 1.24 (AAFTF polish; Walker et al. 2014) by aligning reads back to the assembled contigs.

For the ONT long-read data, *de novo* assembly was conducted using Canu v2.2 (Koren et al. 2017) assuming an estimated genome size of 20 Mb. Long-read assembly accuracy was further improved by iterative polishing using Pilon v1.24 (up to 5 rounds) using corresponding Illumina short reads integrated via AAFTF. PacBio HiFi reads were assembled using Hifiasm 0.19.7 (Cheng et al. 2021). Assembly completeness was evaluated using the AAFTF *assess* module. All Raw sequencing reads and finalized genome assemblies have been deposited in NCBI GenBank/SRA under the BioProject accessions listed in Table S1 and S2. Assembly metrics and parameter settings for long-read (ONT and PacBio) and short-read (Illumina) datasets are summarized in Table S1 and Table S2, respectively.

Gene prediction and annotation were conducted using funnanotate v1.8.17 (Palmer and Stajich 2020), using public RNA-seq datasets retrieved from the NCBI SRA to train and refine gene models. First, assembled scaffolds were sorted by length using the AAFTF *sort* module and soft-masked for repetitive elements with RepeatMasker v4.1.4 (Smit and Hubley 2026) via funannotate *mask*. Training parameters for gene predictors were generated from RNA-seq evidence via funannotate *train*. Gene prediction was performed using funannotate *predict*, which incorporates AUGUSTUS, GeneMark-ES/ET, and EVidenceModeler to generate consensus gene models, supported by protein evidence from the curated UniProtKb/SwissProt database. Functional annotation of predicted proteins was performed with funannotate *annotate*, which integrates domain profiles, pathways, and family classifications from PFAM, InterProScan, Gene Ontology (GO), eggNOG, dbCAN2, UniprotDB, antiSMASH, and MEROPS. Finally, genome assembly and gene model completeness were evaluated by BUSCO v5.5.0 (Manni et al. 2021) in genome mode against the fungi_odb10 marker set.

### Identification of centromeric and telomeric regions

To evaluate chromosome completeness, canonical telomeric regions (TAAC[3-8] or its reverse complement) were located using the ‘find_telomeres.py’ script (https://github.com/markhilt/genome_analysis_tools/blob/master/find_telomeres.py; Hiltunen et al. 2021). Putative centromeric regions were inferred by mapping localized shifts in GC content and gene density along contigs. GC content was calculated with non-overlapping 0.25 kb windows using SeqKit v2.4.0 (Shen et al. 2016), following established parameters (Coelho et al. 2023). Gene counts per 1-kb non-overlapping window were obtained from GFF files using SAMtools v1.22.1 (Danecek et al. 2021) and BEDtools v2.30.0 (Quinlan and Hall 2010). Windowed GC content and gene density tracks were plotted along chromosome scaffolds using the R/Bioconductor package karyoploteR v1.22 (Gel and Serra 2017) to delineate putative centromeric boundaries.

### Subgenome separation for hybrid strains

Subgenomes were phased using a reference-guided approach that utilized the haploid non-hybrid genome *R. mucilaginosa* Y-2510 to partition homeologous regions across four diploid strains. Pairwise whole-genome alignments between the *R. mucilaginosa* Y-2510 and each hybrid assembly were performed using Minimap2 v2.24 (Li 2018) with the ‘-x asm5’ parameter preset. Alignment records in the resulting paf files were filtered to retain only high-confident alignments with a mapping quality score above 60. Synteny plots generated by D-GENIES (Cabanettes and Klopp 2018) were then used to partition filtered alignment into two distinct subsets corresponding to parental subgenomes. Contigs assigned to each subgenome were subsequently extracted using a custom Python script.

### Whole-genome pairwise identity

Pairwise average nucleotide identity (ANI) among genome assemblies was calculated using Mash 2.3 (Ondov et al. 2016). Representative sketches were generated with a k-mer length of 16 nt and a sketch size of 10,000, from which global mutation distances were estimated under default settings. Digital DNA-DNA hybridization (dDDH) values among representative strains across species were calculated by the Genome-to-Genome Distance Calculator 3.0 (GGDC) via the DSMZ website, employing Formula 2 (https://ggdc.dsmz.de/ggdc.php<u>#</u>; Meier-Kolthoff et al. 2013). To examine structural synteny and genome-wide variation across species, pairwise alignments were generated with MUMmer 4 (Marçais et al. 2018). Initial alignments were conducted using nucmer with the ‘--mum’ parameter. Alignments were then filtered using the delta-filter with parameters ‘-1’ (to retain 1-to-1 mapping) and ‘-l 5000’ (to enforce a minimal alignment length threshold of 5,000 bp).

### Multi-locus and phylogenomic tree reconstruction

To infer species-level relationships across *Rhodotorula*, a maximum likelihood (ML) tree was constructed from a concatenated five-gene alignment comprising the internal transcribed spacer (ITS) region, the D1/D2 domains of the large subunit (LSU) rRNA, two RNA polymerase II subunits (*RPB1* and *RPB2*), and translation elongation factor 1-α (*TEF1*). Nucleotide sequences of these loci were identified from genomes using fasta36 (Pearson 2016). Sequences of each locus were aligned individually using MAFFT v7.505 (Katoh and Standley 2013) and concatenated into a single alignment matrix using the ‘catfasta2phyml.pl’ (https://github.com/nylander/catfasta2phyml). ML tree was inferred using IQ-TREE v2.2.2.6 (Minh et al. 2020) under the GTR+I+G substitution model, with branch support evaluated using 1,000 bootstrap replicates.

To thoroughly cross-verify the topology of the five-gene tree against genome-scale signal, we conducted two independent phylogenomic analyses on both the 20-genome comparative and the 249-genome datasets. First, we used Phyling v2.0 (Tsai and Stajich 2026) to identify and extract single-copy orthologs from the BUSCO fungi_odb10 marker set across the 20 genomes. An ML species tree was reconstructed from the concatenated, partitioned alignment using IQ-TREE v2.2.2.6, with best-fit substitution models for each partition determined via ModelFinder. Second, we used OrthoFinder v2.5.5 to infer individual gene trees using the DendroBLAST algorithm (Emms and Kelly 2019). A coalescent-based species tree was inferred from the full set of all unrooted gene trees using STAG and rooted with STRIDE (Emms and Kelly 2018). Topological alignment and branch congruence across these methods were compared and visualized using Tanglegram in interactive tree of life (iTOL) v 7.1.1 (Letunic and Bork 2019).

Deduced protein sequences for genes residing within and adjacent to *MAT* loci were annotated using Miniprot (Li 2023) and parsed using a custom script. Sequence alignment and ML tree construction were conducted as described above. Consensus trees were visualized using iTOL. Gene presence-absence patterns and copy number variations for four key mating-related genes, namely *STE3*, *STE20*, *HD1* and *HD2*, were mapped onto the 249-genome phylogeny using ggtree (Yu et al. 2017).

### Evolutionary rate and selection analyses of *STE3*

To examine the evolutionary rate and selective constraints acting on the pheromone receptor gene *STE3*, we estimated nonsynonymous substitution (*d_N_*) and synonymous (*d_S_*) substitution rates and their ratio (ω = *d_N_*/*d_S_*), using *codeml* in PAML v4.9 (Yang 2007). We employed a ‘two-ratio’ branch model (model = 2, NSsites = 0) to test whether ω differed between foreground and background clades. This alternative model was evaluated against a null model assuming a uniform ω across all branches (model = 0). Statistical significance was assessed via likelihood ratio tests (LRTs), where the test statistic was calculated as ΔlnL = 2 × (lnL_model2 − lnL_model0) and compared against a chi-squared distribution with degrees of freedom equal to the difference in parameter counts between models.

To evaluate whether elevated ω in foreground branches were driven by positive selection or relaxed purifying selection, we tested it with both branch-site model in PAML package and RELAX model in HyPhy v2.5 (Wertheim et al. 2015). For branch-site tests, mode A (model = 2, NSsites = 2, fix_omega = 0, omega = 0.5) was compared to its null model (model = 2, NSsites = 2, fix_omega = 1, omega = 1). Sites under positive selection were identified via the Bayes Empirical Bayes approach when *p* < 0.05 (Álvarez-Carretero et al. 2023). For relaxed selection tests, the RELAX model was fitted to estimate the selection intensity parameter *k*, where *k* < 1 indicates relaxed selection and *k* > 1 suggests intensified selection. Significance for RELAX tests was assessed via LRTs against a null model of *k* = 1 (*p* < 0.05).

### Comparative orthogroup inference

Orthogroup inference was performed using OrthoFinder v2.5.5 on predicted proteomes from the 20-genome comparative dataset. Comparative genomic filtering was conducted to identify lineage-specific orthogroups retained exclusively in *R*. *mucilaginosa* A2-type genomes but absent from A1-type and other *Rhodotorula* species. Presence-absence patterns across lineages were visualized using UpsetR v1.4.0 (Conway et al. 2017).

To verify that A2-associated orthogroups represented transcriptionally supported genes rather than pseudogenes or assembly/annotation artifacts, 22 public *R*. *mucilaginosa* RNA-seq datasets were retrieved from the NCBI SRA. Raw RNA-seq reads were quality-filtered and adaptor-trimmed using fastp. Cleaned reads were aligned to the *R*. *mucilaginosa* Y-2510 reference genome using HISAT2 (Kim et al. 2019) and post-processed through SAMtools. Gene-level read abundances were quantified using featureCounts from the Subread package (Liao et al. 2014) based on the *R*. *mucilaginosa* Y-2510 genome annotation, and the expression matrix was visualized using the R package pheatmap (Kolde 2019). Genes were considered transcriptionally supported if at least one paired-end RNA-seq fragment was assigned to the annotated gene region in at least one RNA-seq dataset.

### *MAT* locus identification, synteny analysis, and transposable element annotation

Scaffolds encompassing putative *MAT* regions in the *R*. *mucilaginosa* Y-2510 genome were identified through BLASTP searches using well-annotated *MAT* locus from its relative *Leucosporidium scottii* CBS 5931 (Maia et al. 2015) as references. Retrieved scaffolds of *R. mucilaginosa* Y-2510 were manually re-annotated using tBLASTN queries against the NCBI nucleotide database. The finalized *MAT* genes from *R. mucilaginosa* Y-2510 were used to identify corresponding *MAT* genes across other *Rhodotorula* genomes by reciprocal blast search. In cases where short exon lengths, intron structures, or sequence hypervariability prevented initial detection of *MAT*-specific genes (*STE3*, *RHA*, *HD1* and *HD2*), protein-to-genome alignments were performed using Miniprot with verified homolog sequences from other species. Putative *RHA* genes identified by Miniprot were manually validated by confirming the presence of a conserved C-terminal CAAX motif characteristic of fungal mating pheromones (Akada et al. 1989; Coelho et al. 2008). To minimize artifacts, scaffolds that have fewer than five *MAT* genes were excluded from downstream comparisons. Syntenic organization of *MAT* regions was visualized using pyGenomeViz v1.0.0 (Shimoyama 2024). Species-level synteny maps were linked via BLASTN-derived collinear blocks between adjacent species, whereas genus-level synteny maps were anchored using reciprocal blast search and Miniprot hits against the *R. mucilaginosa* Y-2510.

To characterize transposable elements (TEs) within the *P/R* loci across *Rhodotorula*, we constructed a custom non-redundant TE library for target region annotation. First, *de novo* TE identification was performed on whole-genome assemblies of three representative stains, *R*. *mucilaginosa* Y-2510 from Clade A, *R*. *toruloides* CBS 14 from Clade B, and R. kratochvilovae DBVPG 10753 from Clade C, using EDTA package (Ou et al. 2019). The resulting TE sequences identified across these genomes were clustered with CD-HIT (Fu et al. 2012) to collapse redundant queries and isolate unique TE families. This curated TE library was supplied as a custom database to RepeatMasker to annotate TE distribution and density across the *P/R* locus in all *Rhodotorula* strains.

### Variant calling and LD decay assay

To investigate the sexuality of *R*. *mucilaginosa*, 41 *R*. *mucilaginosa* strains were selected for single-nucleotide polymorphism (SNP) calling and LD decay analysis, with 16 *R*. *diobovata* strains, which known to be sexual, serving as a comparative baseline. Sequencing reads were processed through a custom Nextflow (Di Tommaso et al. 2017) pipeline (https://github.com/chtsai0105/nf_snp_calling). Raw reads were quality-filtered using fastp (Chen et al. 2018) and aligned to the reference genome (Y-2510 for *R. mucilaginosa* and Y-7196 for *R. diobovata*) with bwa-mem2 (Vasimuddin et al. 2019). Alignment files were cleaned by SAMtools (*sort*, *fixmate*, and *markdup*) to keep only paired and unique hits.

A two-step variant calling strategy was performed via BCFtools (Danecek et al. 2021) to mimic GATK’s joint-calling pipeline with optimized computational efficiency. An initial round of variant discovery was performed individually across each deduplicated alignment via BCFtools *mpileup*. The resulting individual variant calls were merged, filtered, and chunked to facilitate parallelized joint variant calling in the second round. The final variant set was filtered to retain strictly high-quality, biallelic SNPs. Detailed parameters and tool flags for each pipeline execution step are documented in the ‘nextflow.config’ file within the linked repository.

The LD decay was evaluated on the filtered variant dataset using PopLDdecay (Zhang et al. 2019). To accommodate haploid variant calls within the diploid-centric framework of PopLDdecay, BCFtools was used to convert haploid genotypes to pseudo-diploid format. Variants located within the *MAT* locus were excluded, and subsequent LD decay assays were conducted exclusively across non-*MAT* genomic regions.

To estimate half-decay values (LD50, in base pairs), pairwise linkage disequilibrium *r*^2^ values were averaged across SNP pairs within each distance interval for each species. The initial *r*^2^ was calculated as the weighted mean *r*^2^ within the first 5 kb of physical distance, while the baseline *r*^2^ was estimated from the long-distance region representing background noise. The *r*^2^ midpoint corresponding to half decay was calculated as (baseline *r*^2^ + (initial *r*^2^ - baseline *r*^2^)/2). The physical distance at which the LD decay curve crossed this midpoint was calculated via linear interpolation using *approx* function in R.

## Acknowledgments

We are grateful to the ARS Culture Collection, as part of microbial strains used in this work were provided by the USDA-ARS Culture Collection (NRRL). We also appreciate the support from UC Riverside High Performance Computing Cluster for conduction of computational analyses, supported by grants from the National Science Foundation (DBI-1429826 & DBI-2215705) and the National Institutes of Health (NIH) (S10-OD016290). Special thanks go to Tania Kurbessoian, Mark Yacoub, Sadikshya Sharma, and Jessica Wu-Woods for their assistance with laboratory work. We also acknowledge Leila Shadmani, Kian Kelly, and Carolina Piña Páez for their valuable support during the research.

## Author contributions

J.E.S and X.Z.L conceptualized and supervised the study. B.T., C.G, N.G.C, X.Z.L, C.C., L.S established the resource collection. X.Z.L conducted strain purification, DNA extraction, and library preparation. J.E.S and X.Z.L performed data curation. X.Z.L, C.H.T, and J.E.S analyzed the genomes. X.Z.L, C.H.T, M.A.C produced the figures. X.Z.L, C.H.T, M.A.C, and J.E.S constructed the methodology. J.E.S, X.Z.L, C.G, N.G.C, I.W acquired funding. X.Z.L wrote the original draft with input from all coauthors. C.H.T contributed to writing – review & editing through substantial structural overhaul and critical revisions of the text. All authors read and approved the manuscript.

