## Supplementary figures S1 to S14; Supplementary text S1 for "Genomic plasticity of the mating-type loci underlies reproductive strategy transitions in *Rhodotorula* yeasts"

**This PDF file includes:**

Supplementary figures S1 to S14

Supplementary text S1

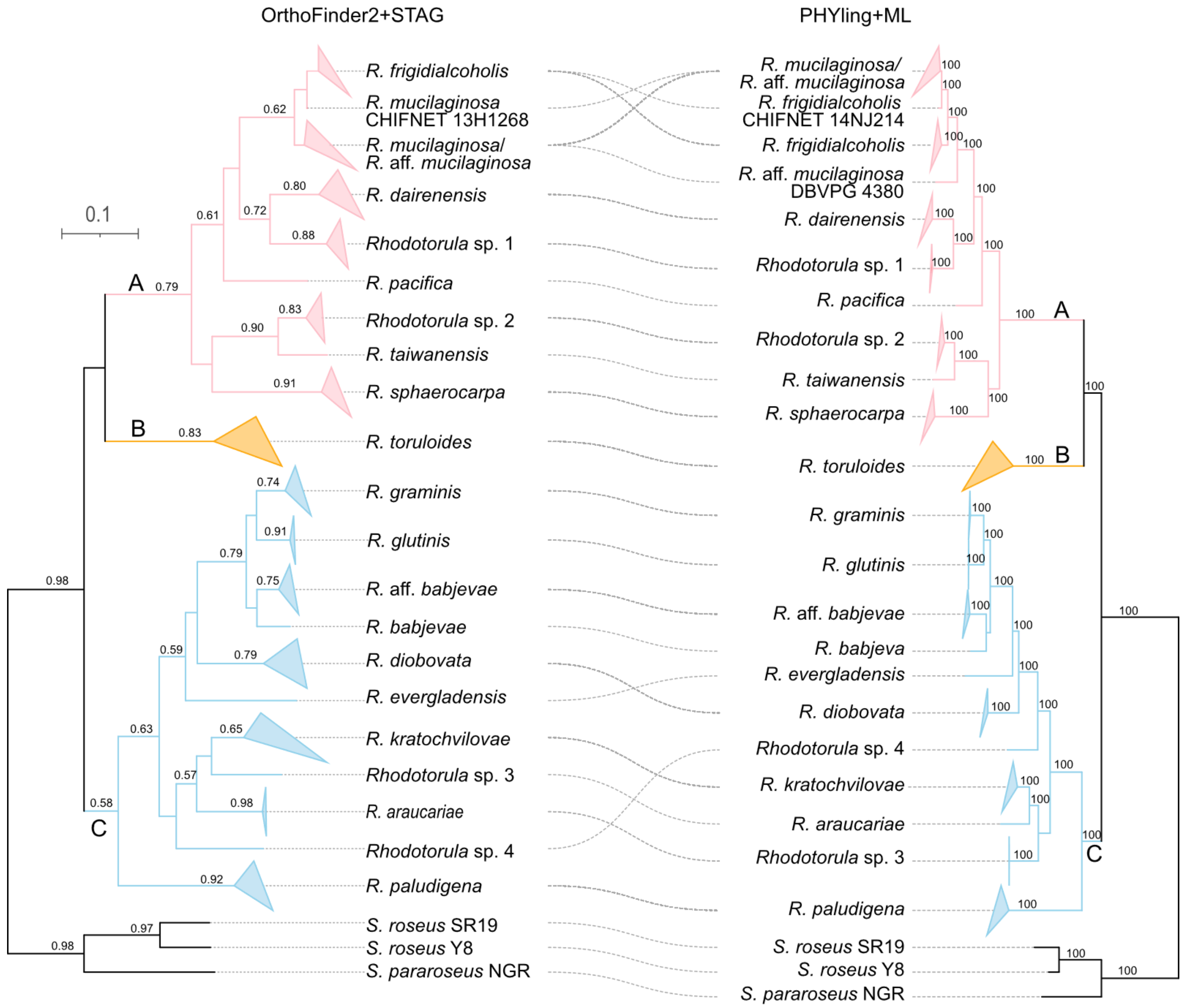


**Supplementary Figure 1. Species tree of the genus *Rhodotorula* inferred from the second dataset comprising 249 strains.** The coalescent species tree on the left was inferred by OrthoFinder (STAG), with bold branches indicating support values above 0.5 (gene tree proportion supporting the corresponding bipartition). The concatenated maximum-likelihood species tree on the right was inferred by Phying, with bold branches indicating bootstrap support above 96% (1,000 replicates). The tree is rooted using three outgroup species (*Sporobolomyces pararoseus* NGR, *S. roseus* SR19, and *S. roseus* Y8). Clades A, B, and C are color labeled on each tree.

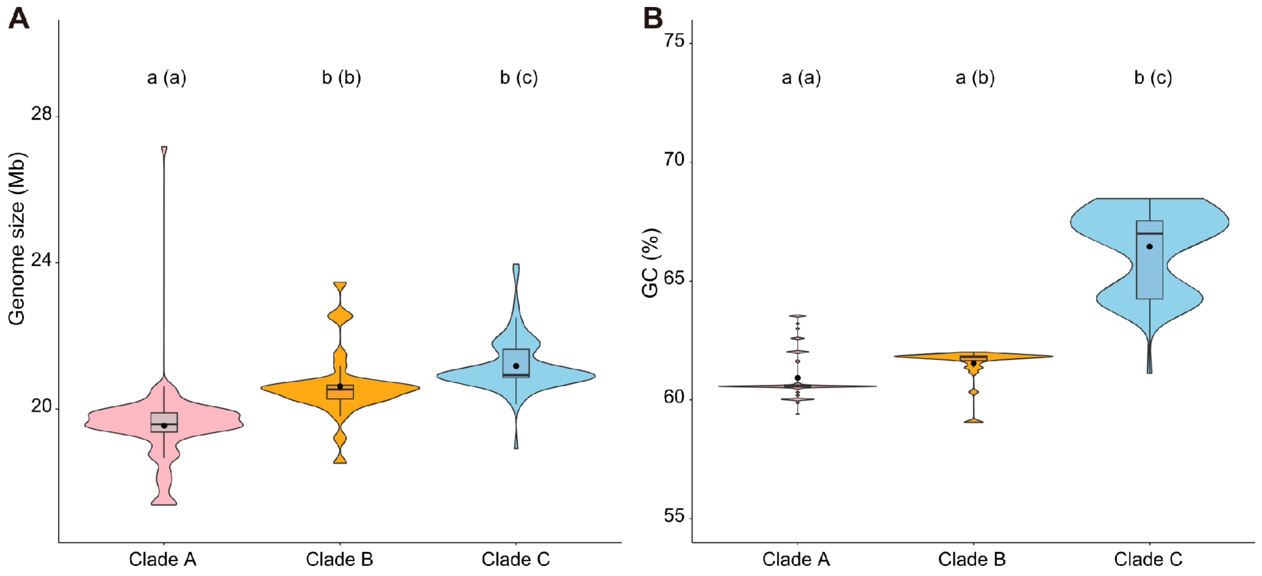


**Supplementary Figure 2. Genomic features across *Rhodotorula* clades. (A)** Genome size (Mb) and **(B)** GC content (%) variations across Clade A, B and C. Violin plots show distribution density, with embedded boxplots indicating the median (horizontal bar), mean (black dot) and interquartile range. Different letters indicate the significant differences among clades according to Tukey's HSD post hoc comparisons following a significant overall ANOVA, with letters outside and inside parentheses representing significance thresholds at $p < 0.01$ and $p < 0.05$, respectively).

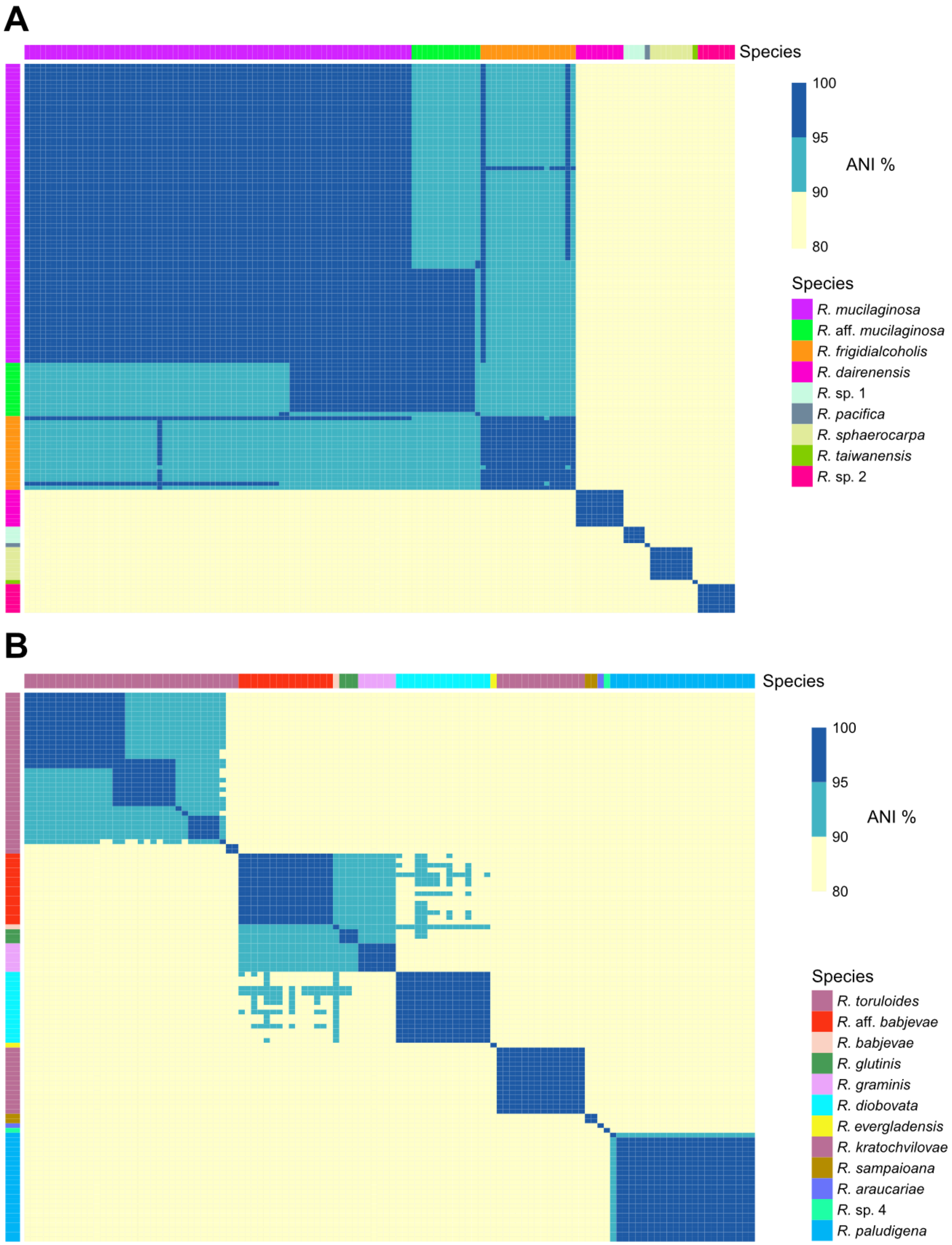
A
Species
100
95
90
80
ANI %
Species
R. mucilaginosa
R. aff. mucilaginosa
R. frigidialcoholis
R. dairenensis
R. sp. 1
R. pacifica
R. sphaerocarpa
R. taiwanensis
R. sp. 2
B
Species
100
95
90
80
ANI %
Species
R. toruloides
R. aff. babjevae
R. babjevae
R. glutinis
R. graminis
R. diobovata
R. evergladensis
R. kratochvilovae
R. sampaioana
R. araucariae
R. sp. 4
R. paludigena

**Supplementary Figure 3. Average Nucleotide Identity (ANI) comparisons across *Rhodotorula* strains.** Pairwise ANI values calculated using Mash for strains in **(A)** Clade A and **(B)** Clades B and C. Strains are color-coded by designated species along the top and left axes. Matrix cells are color-coded by ANI thresholds: dark blue (>95% ANI, standard species boundary); cyan (90-95% ANI); and yellow (< 90% ANI).

**B**

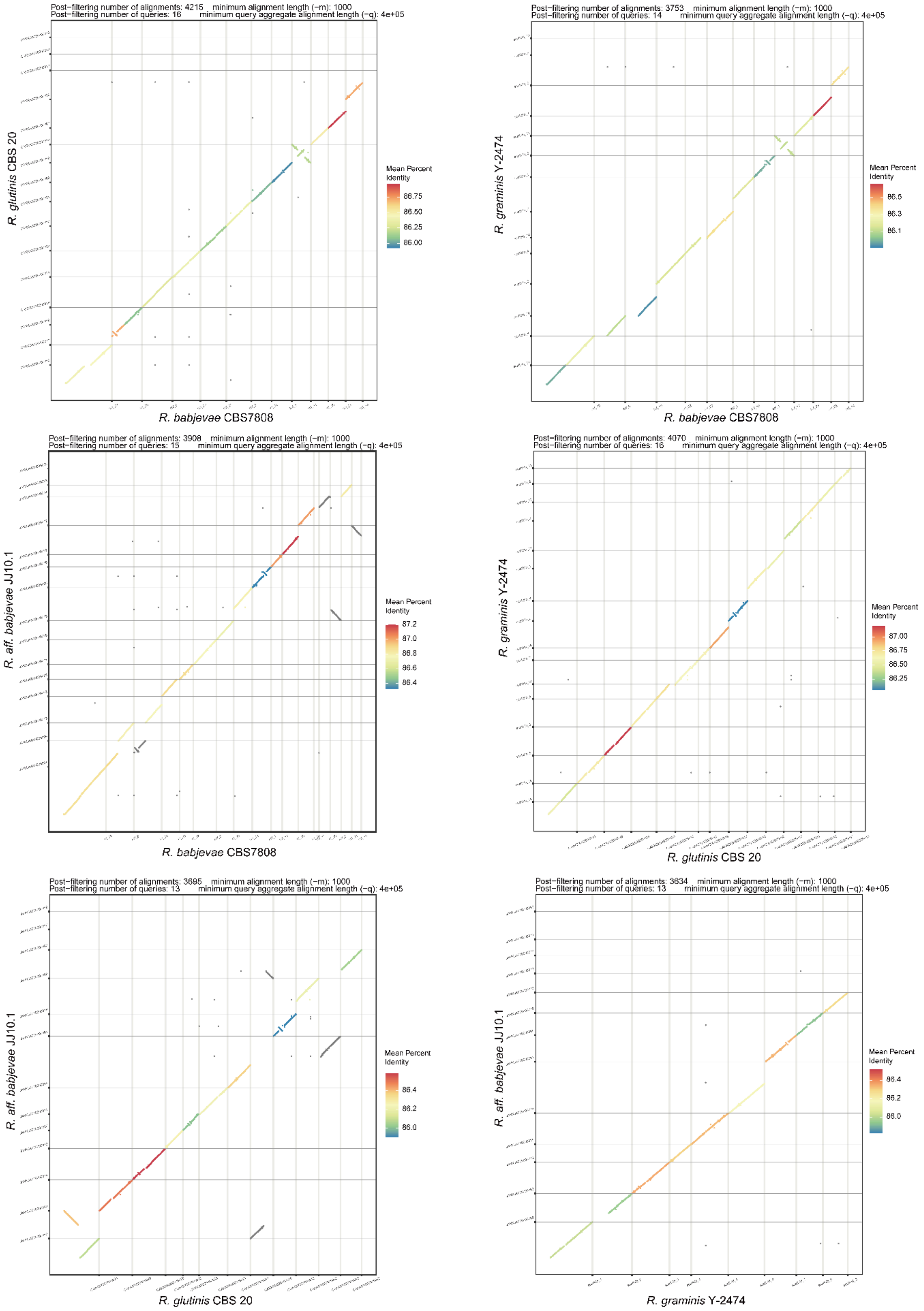
Post−filtering number of alignments: 4215 minimum alignment length (−m): 1000
Post−filtering number of queries: 16 minimum query aggregate alignment length (−q): 4e+05
R. glutinis CBS 20
R. babjevae CBS7808
Mean Percent Identity
86.75
86.50
86.25
86.00
Post−filtering number of alignments: 3753 minimum alignment length (−m): 1000
Post−filtering number of queries: 14 minimum query aggregate alignment length (−q): 4e+05
R. graminis Y-2474
R. babjevae CBS7808
Mean Percent Identity
86.5
86.3
86.1
Post−filtering number of alignments: 3908 minimum alignment length (−m): 1000
Post−filtering number of queries: 15 minimum query aggregate alignment length (−q): 4e+05
R. aff. babjevae JJ10.1
R. babjevae CBS7808
Mean Percent Identity
87.2
87.0
86.8
86.6
86.4
Post−filtering number of alignments: 4070 minimum alignment length (−m): 1000
Post−filtering number of queries: 16 minimum query aggregate alignment length (−q): 4e+05
R. graminis Y-2474
R. glutinis CBS 20
Mean Percent Identity
87.00
86.75
86.50
86.25
Post−filtering number of alignments: 3695 minimum alignment length (−m): 1000
Post−filtering number of queries: 13 minimum query aggregate alignment length (−q): 4e+05
R. aff. babjevae JJ10.1
R. glutinis CBS 20
Mean Percent Identity
86.4
86.2
86.0
Post−filtering number of alignments: 3634 minimum alignment length (−m): 1000
Post−filtering number of queries: 13 minimum query aggregate alignment length (−q): 4e+05
R. aff. babjevae JJ10.1
R. graminis Y-2474
Mean Percent Identity
86.4
86.2
86.0

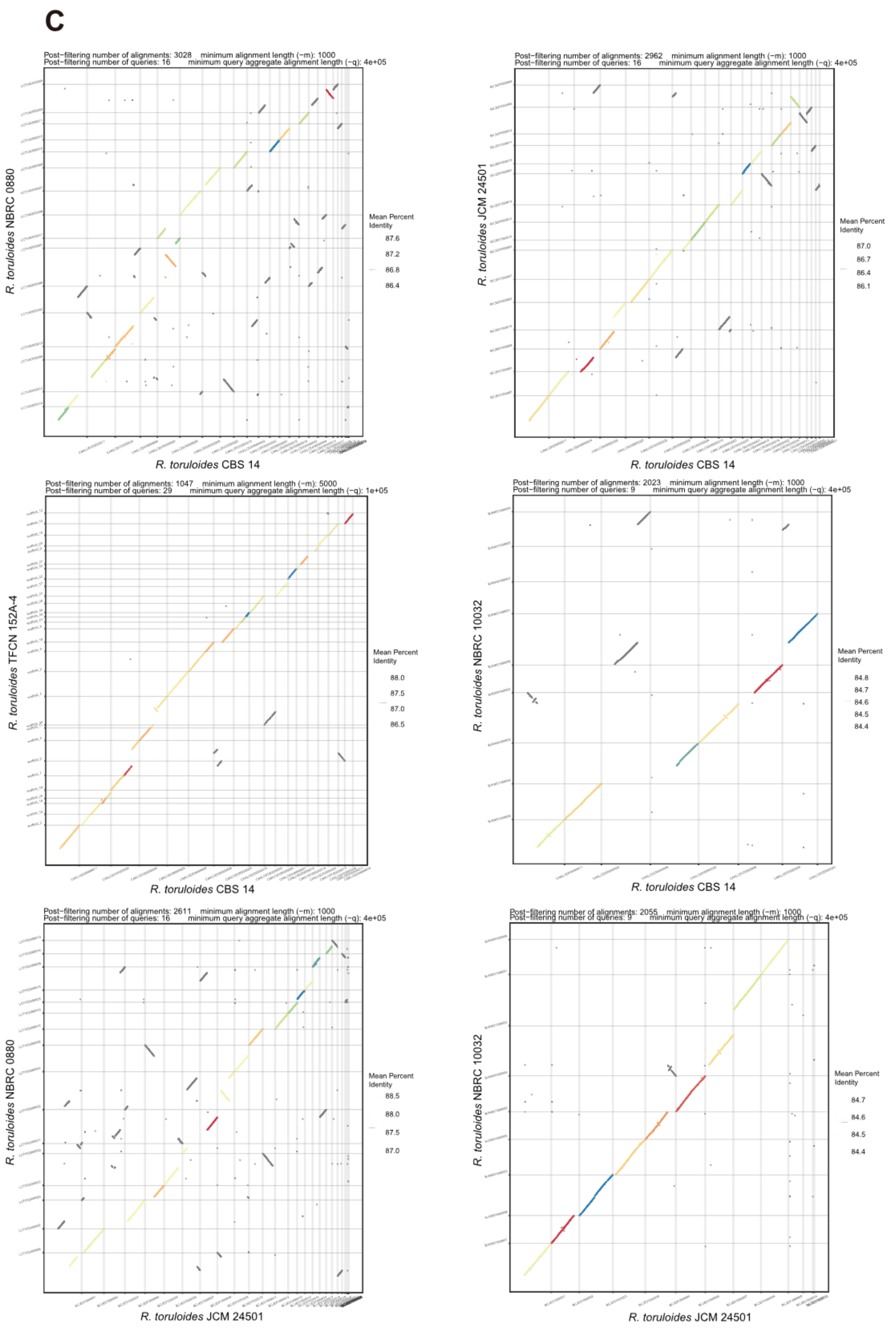
C
Post−filtering number of alignments: 3028 minimum alignment length (−m): 1000
Post−filtering number of queries: 16 minimum query aggregate alignment length (−q): 4e+05
R. toruloides NBRC 0880
R. toruloides CBS 14
Mean Percent Identity
87.6
87.2
86.8
86.4
Post−filtering number of alignments: 2962 minimum alignment length (−m): 1000
Post−filtering number of queries: 16 minimum query aggregate alignment length (−q): 4e+05
R. toruloides JCM 24501
R. toruloides CBS 14
Mean Percent Identity
87.0
86.7
86.4
86.1
Post−filtering number of alignments: 1047 minimum alignment length (−m): 5000
Post−filtering number of queries: 29 minimum query aggregate alignment length (−q): 1e+05
R. toruloides TFCN 152A-4
R. toruloides CBS 14
Mean Percent Identity
88.0
87.5
87.0
86.5
Post−filtering number of alignments: 2023 minimum alignment length (−m): 1000
Post−filtering number of queries: 9 minimum query aggregate alignment length (−q): 4e+05
R. toruloides NBRC 10032
R. toruloides CBS 14
Mean Percent Identity
84.8
84.7
84.6
84.5
84.4
Post−filtering number of alignments: 2611 minimum alignment length (−m): 1000
Post−filtering number of queries: 16 minimum query aggregate alignment length (−q): 4e+05
R. toruloides NBRC 0880
R. toruloides JCM 24501
Mean Percent Identity
88.5
88.0
87.5
87.0
Post−filtering number of alignments: 2055 minimum alignment length (−m): 1000
Post−filtering number of queries: 9 minimum query aggregate alignment length (−q): 4e+05
R. toruloides NBRC 10032
R. toruloides JCM 24501
Mean Percent Identity
84.7
84.6
84.5
84.4

**D**

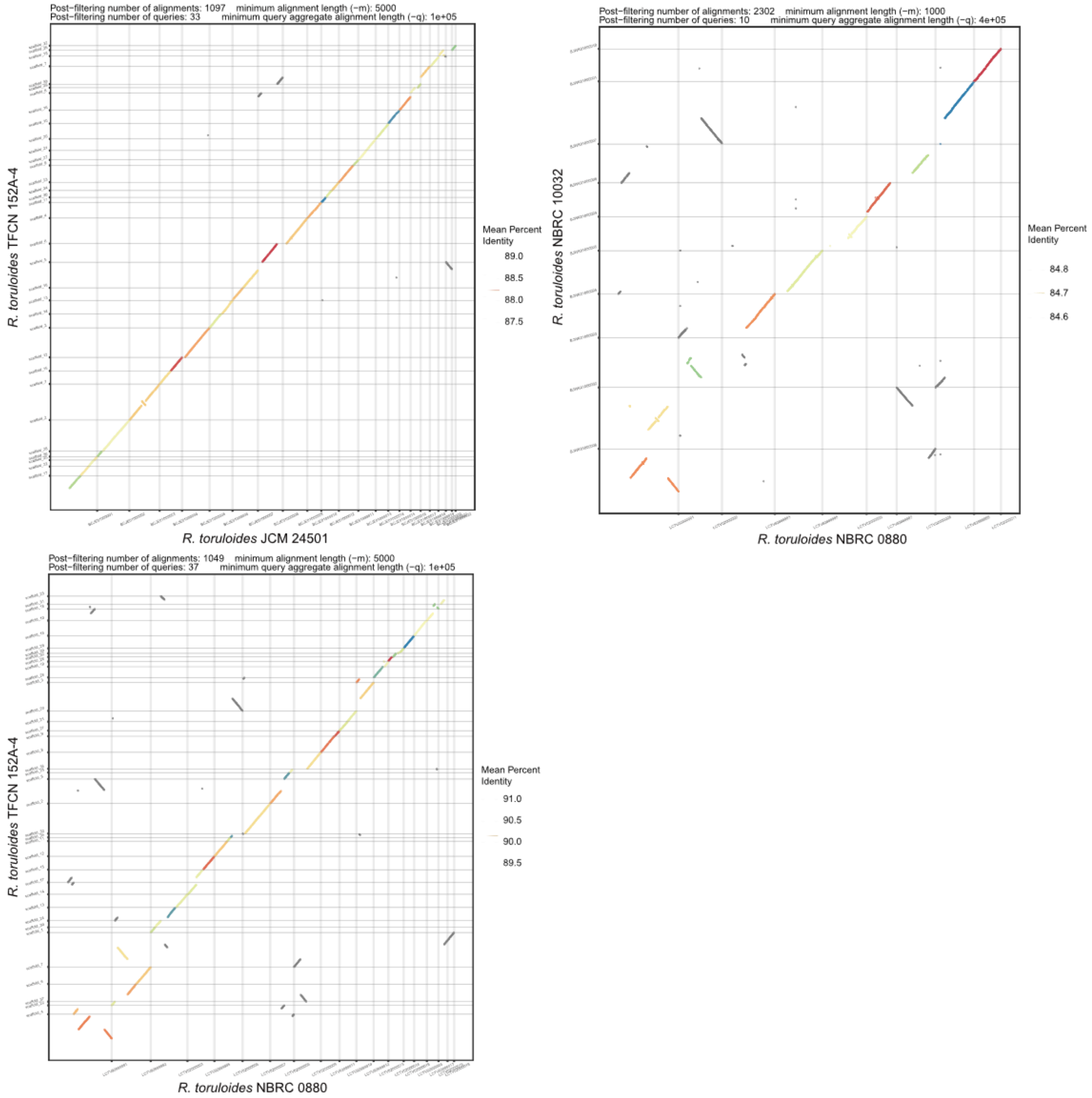


**Supplementary Figure 4. Genome-wide synteny and sequence divergence across *Rhodotorula* species complexes.** Whole-genome dot plots comparing representative species pairs within **(A)** the *R. mucilaginosa* species complex (*R. mucilaginosa*, *R. aff. mucilaginosa*, and *R. frigidialcoholis*), **(B)** the complex in Clade C (*R. aff. babjevae*, *R. babjevae*, *R. graminis*, and *R. glutinis*), and **(C, D)** the *R. toruloides* species complex. Pairwise nucleotide alignments were generated using nucmer in MUMmer v4.0 and visualized with mummerCoordsDotPlotly.R.

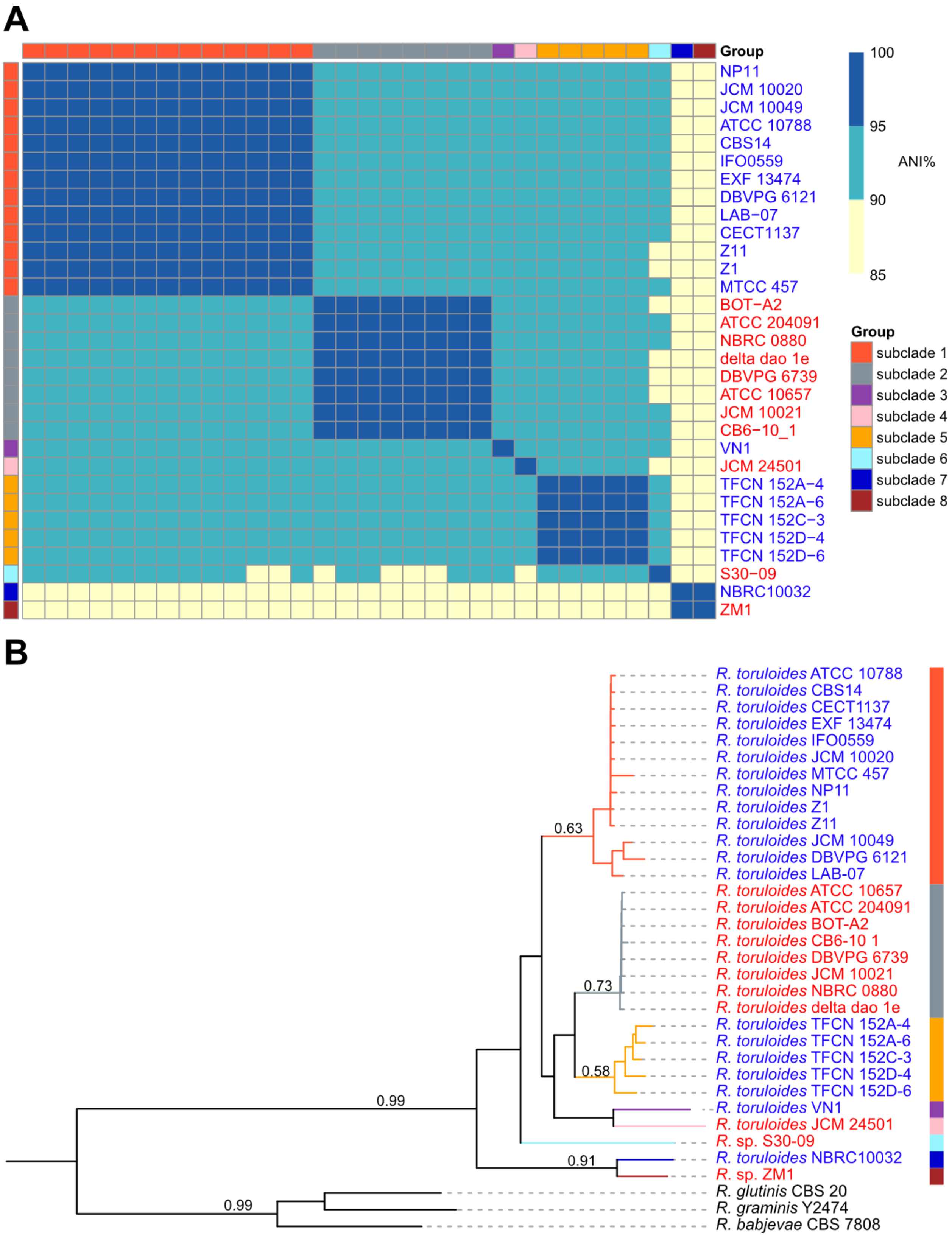
A
Group
NP11
JCM 10020
JCM 10049
ATCC 10788
CBS14
IFO0559
EXF 13474
DBVPG 6121
LAB−07
CECT1137
Z11
Z1
MTCC 457
BOT−A2
ATCC 204091
NBRC 0880
delta dao 1e
DBVPG 6739
ATCC 10657
JCM 10021
CB6−10_1
VN1
JCM 24501
TFCN 152A−4
TFCN 152A−6
TFCN 152C−3
TFCN 152D−4
TFCN 152D−6
S30−09
NBRC10032
ZM1
100
95
90
85
ANI%
Group
subclade 1
subclade 2
subclade 3
subclade 4
subclade 5
subclade 6
subclade 7
subclade 8
B
0.63
0.73
0.58
0.99
0.91
0.99
R. toruloides ATCC 10788
R. toruloides CBS14
R. toruloides CECT1137
R. toruloides EXF 13474
R. toruloides IFO0559
R. toruloides JCM 10020
R. toruloides MTCC 457
R. toruloides NP11
R. toruloides Z1
R. toruloides Z11
R. toruloides JCM 10049
R. toruloides DBVPG 6121
R. toruloides LAB-07
R. toruloides ATCC 10657
R. toruloides ATCC 204091
R. toruloides BOT-A2
R. toruloides CB6-10 1
R. toruloides DBVPG 6739
R. toruloides JCM 10021
R. toruloides NBRC 0880
R. toruloides delta dao 1e
R. toruloides TFCN 152A-4
R. toruloides TFCN 152A-6
R. toruloides TFCN 152C-3
R. toruloides TFCN 152D-4
R. toruloides TFCN 152D-6
R. toruloides VN1
R. toruloides JCM 24501
R. sp. S30-09
R. toruloides NBRC10032
R. sp. ZM1
R. glutinis CBS 20
R. graminis Y2474
R. babjevae CBS 7808

**Supplementary Figure 5. Comparative genomic analysis of the *Rhodotorula toruloides* species complex. (A)** Pairwise Average Nucleotide Identity (ANI) heatmap calculated using Mash. Colors indicate ANI thresholds: dark blue (> 95%, standard species boundary), cyan (90–95%), and yellow (< 90%). **(B)** Coalescent species tree inferred using OrthoFinder. Numbers on branches indicate support values > 0.50. Strain labels are color-coded by mating type: A1 (blue) and A2 (red). Subclades are defined based on the 95% ANI threshold. Note that three aneuploid or diploid strains (PYCC 5615, CGMCC 2.1609, and CCT 0783) were excluded from downstream subdataset analyses.

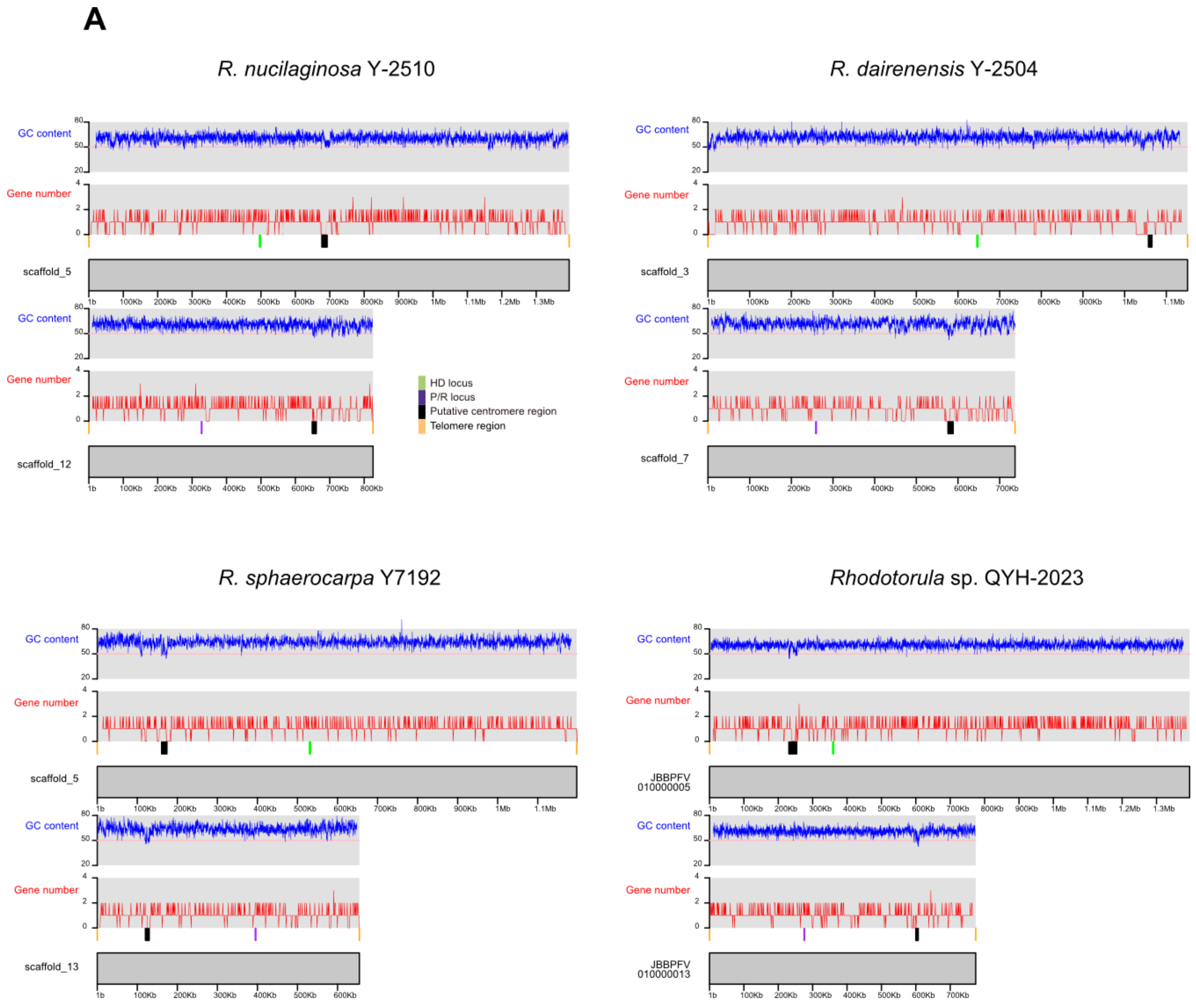
A
R. nucilaginosa Y-2510
R. dairenensis Y-2504
R. sphaerocarpa Y7192
Rhodotorula sp. QYH-2023
GC content
Gene number
scaffold_5
scaffold_12
scaffold_3
scaffold_7
scaffold_13
JBBPFV 010000005
JBBPFV 010000013
HD locus
P/R locus
Putative centromere region
Telomere region

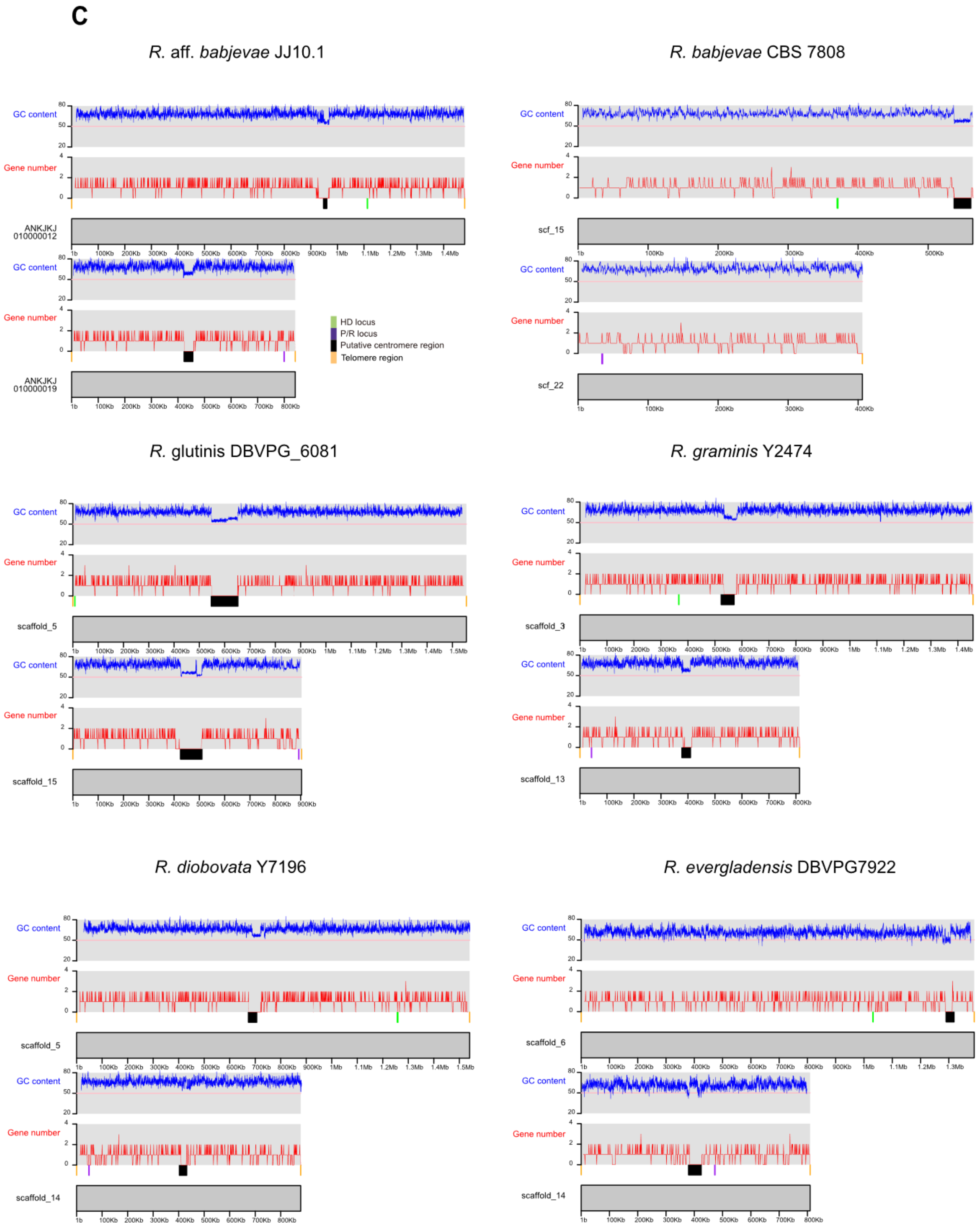
C
R. aff. babjevae JJ10.1
R. babjevae CBS 7808
GC content
Gene number
ANKJKJ 010000012
ANKJKJ 010000019
scf_15
scf_22
HD locus
P/R locus
Putative centromere region
Telomere region
R. glutinis DBVPG_6081
R. graminis Y2474
scaffold_5
scaffold_15
scaffold_3
scaffold_13
R. diobovata Y7196
R. evergladensis DBVPG7922
scaffold_5
scaffold_14
scaffold_6
scaffold_14

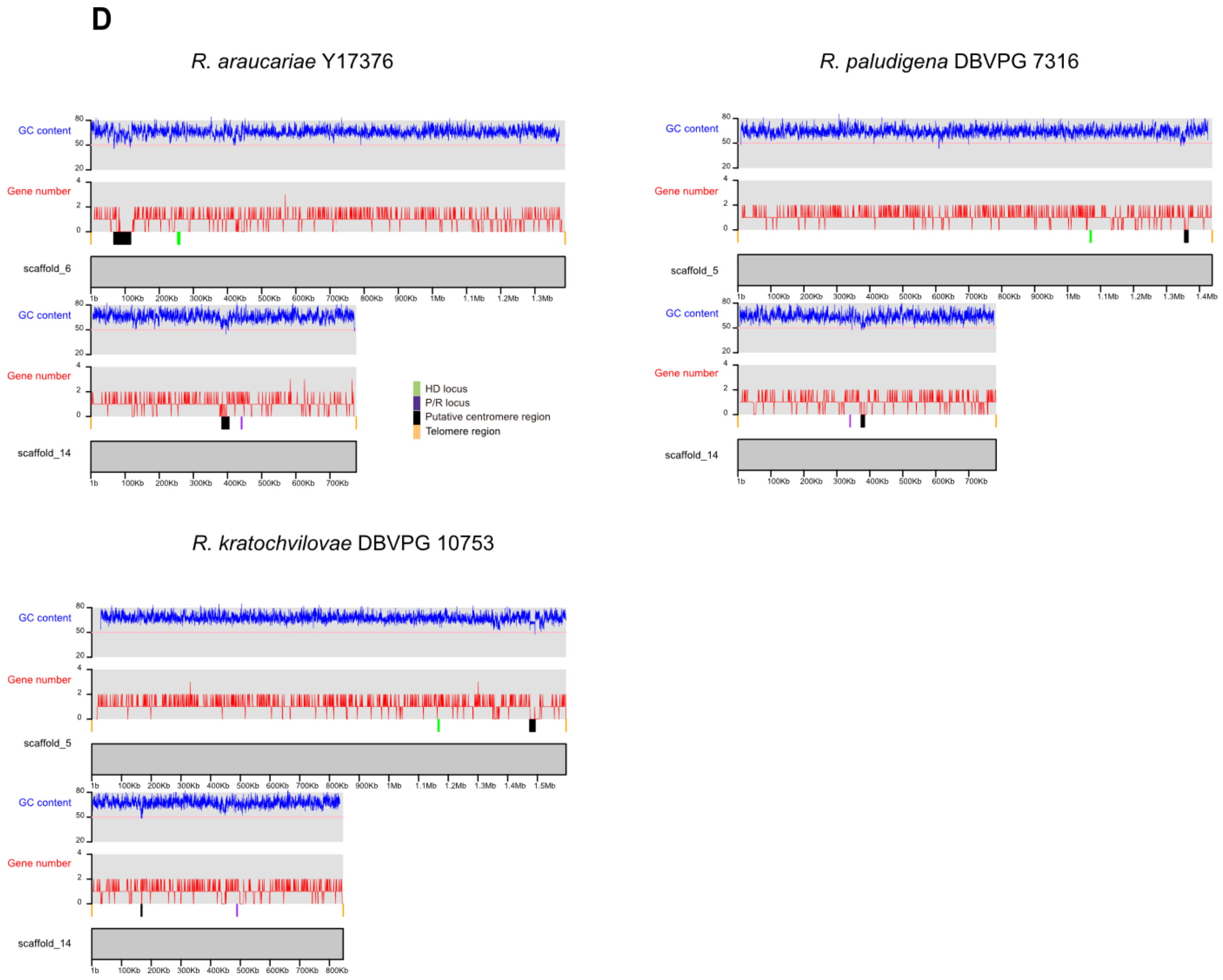


**Supplementary Figure 6. Chromosomal organization of the *MAT* locus in representative *Rhodotorula* species.** Chromosomal plots showing the locations of the *P/R* locus and *HD* locus relative to centromeres and telomeres across representative species from **(A)** Clade A, **(B)** Clade B, and **(C,D)** Clade C. Tracks above each chromosome or contig display GC content and gene density distributions.

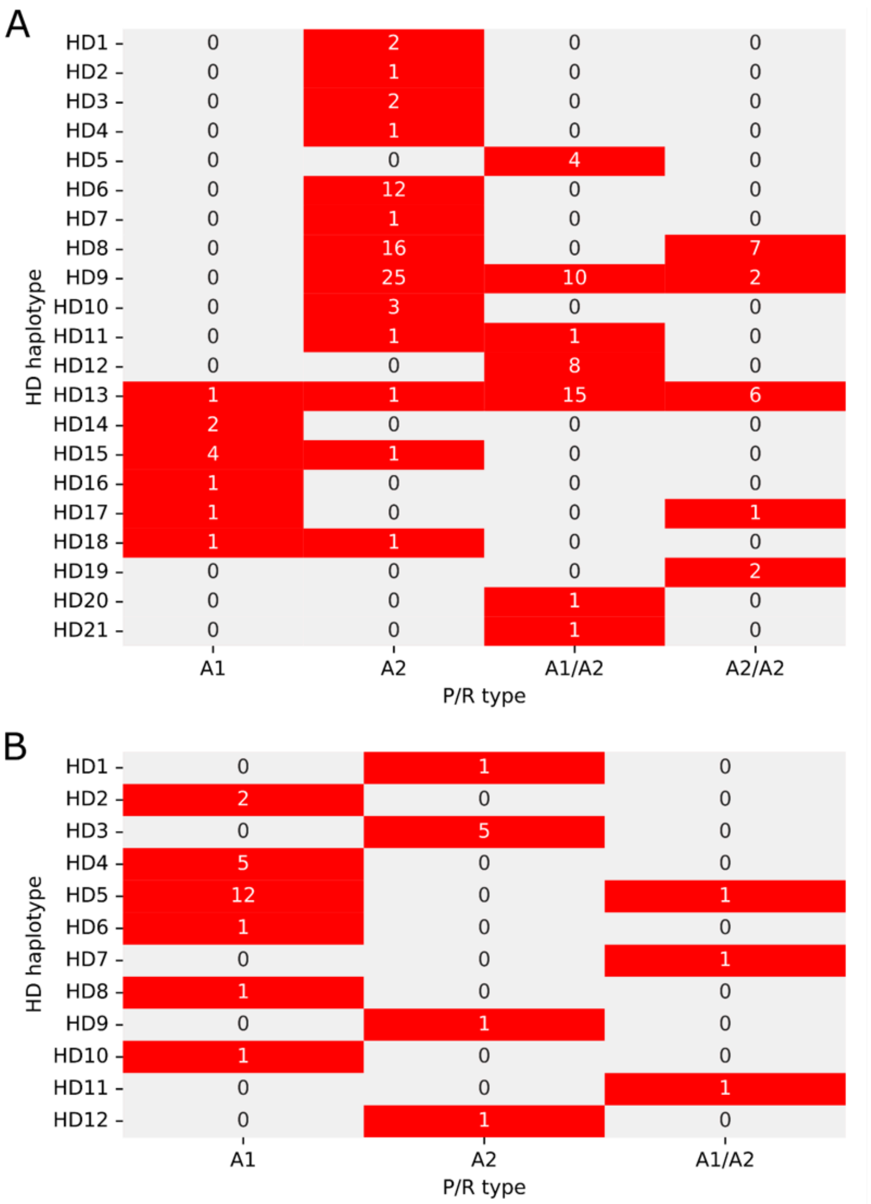


**Supplementary Figure 7. Observed combinations of the *P/R* and *HD* alleles in (A)** *R. mucilaginosa* species complex and **(B)** *R. toruloides* species complex.

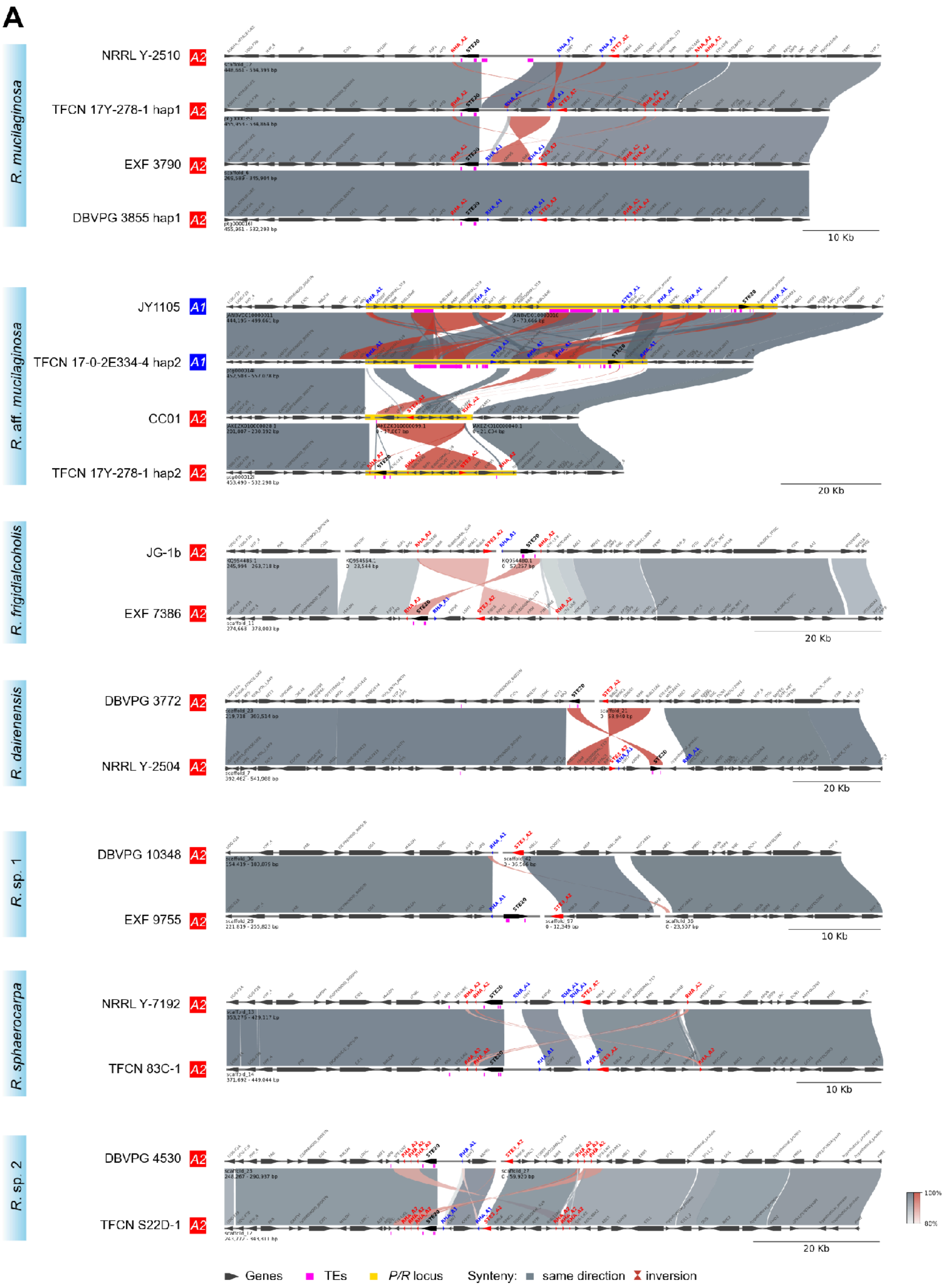
A
R. mucilaginosa
NRRL Y-2510 A2
TFCN 17Y-278-1 hap1 A2
EXF 3790 A2
DBVPG 3855 hap1 A2
10 Kb
R. aff. mucilaginosa
JY1105 A1
TFCN 17-0-2E334-4 hap2 A1
CC01 A2
TFCN 17Y-278-1 hap2 A2
20 Kb
R. frigidialcoholis
JG-1b A2
EXF 7386 A2
20 Kb
R. dairenensis
DBVPG 3772 A2
NRRL Y-2504 A2
20 Kb
R. sp. 1
DBVPG 10348 A2
EXF 9755 A2
10 Kb
R. sphaerocarpa
NRRL Y-7192 A2
TFCN 83C-1 A2
10 Kb
R. sp. 2
DBVPG 4530 A2
TFCN S22D-1 A2
20 Kb
100%
80%
Genes
TEs
P/R locus
Synteny: same direction inversion

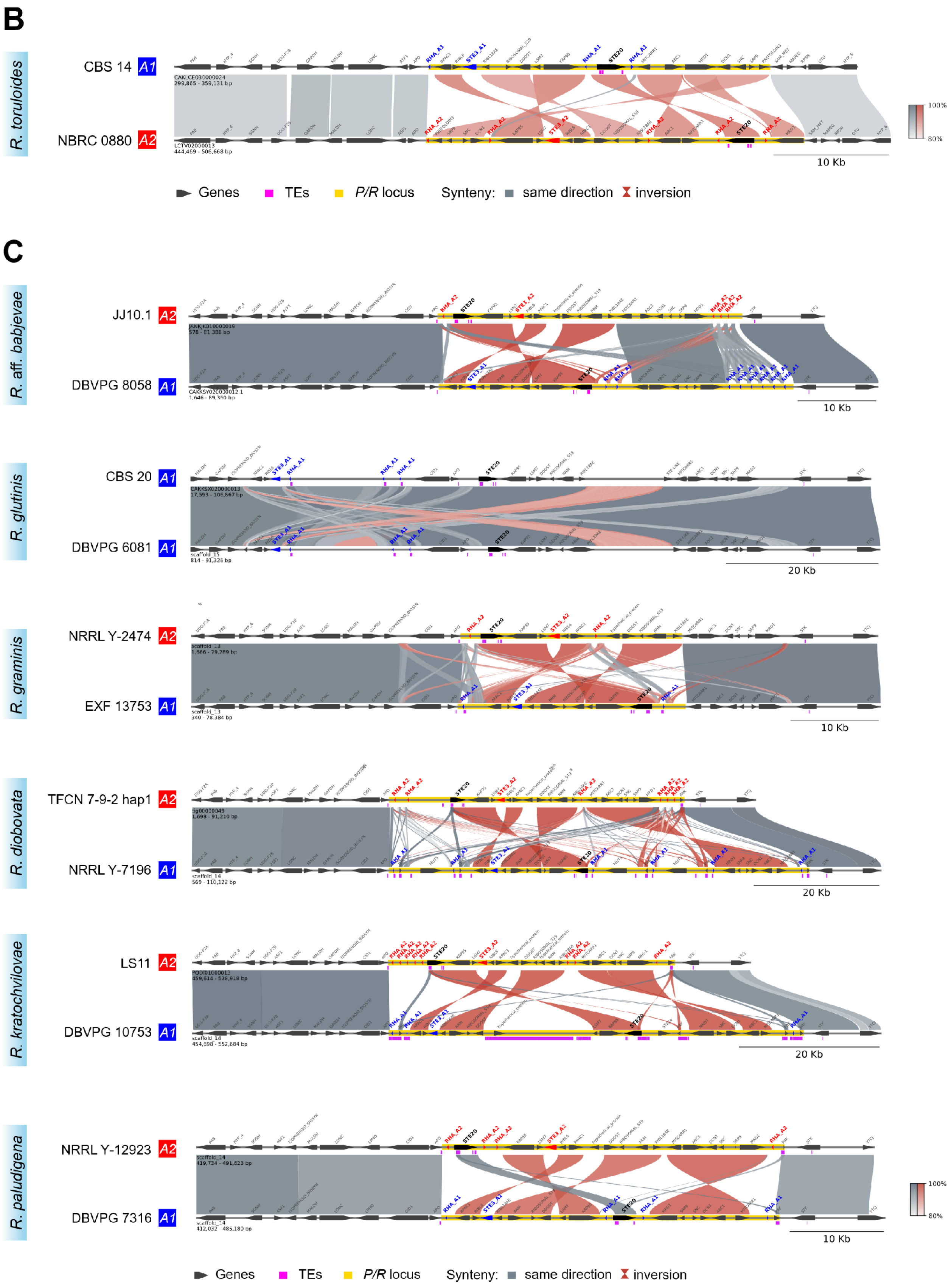
B
R. toruloides
CBS 14 A1
NBRC 0880 A2
10 Kb
100%
80%
Genes
TEs
P/R locus
Synteny:
same direction
inversion
C
R. aff. babjevae
JJ10.1 A2
DBVPG 8058 A1
10 Kb
R. glutinis
CBS 20 A1
DBVPG 6081 A1
20 Kb
R. graminis
NRRL Y-2474 A2
EXF 13753 A1
10 Kb
R. diobovata
TFCN 7-9-2 hap1 A2
NRRL Y-7196 A1
20 Kb
R. kratochvilovae
LS11 A2
DBVPG 10753 A1
20 Kb
R. paludigena
NRRL Y-12923 A2
DBVPG 7316 A1
10 Kb
100%
80%
Genes
TEs
P/R locus
Synteny:
same direction
inversion

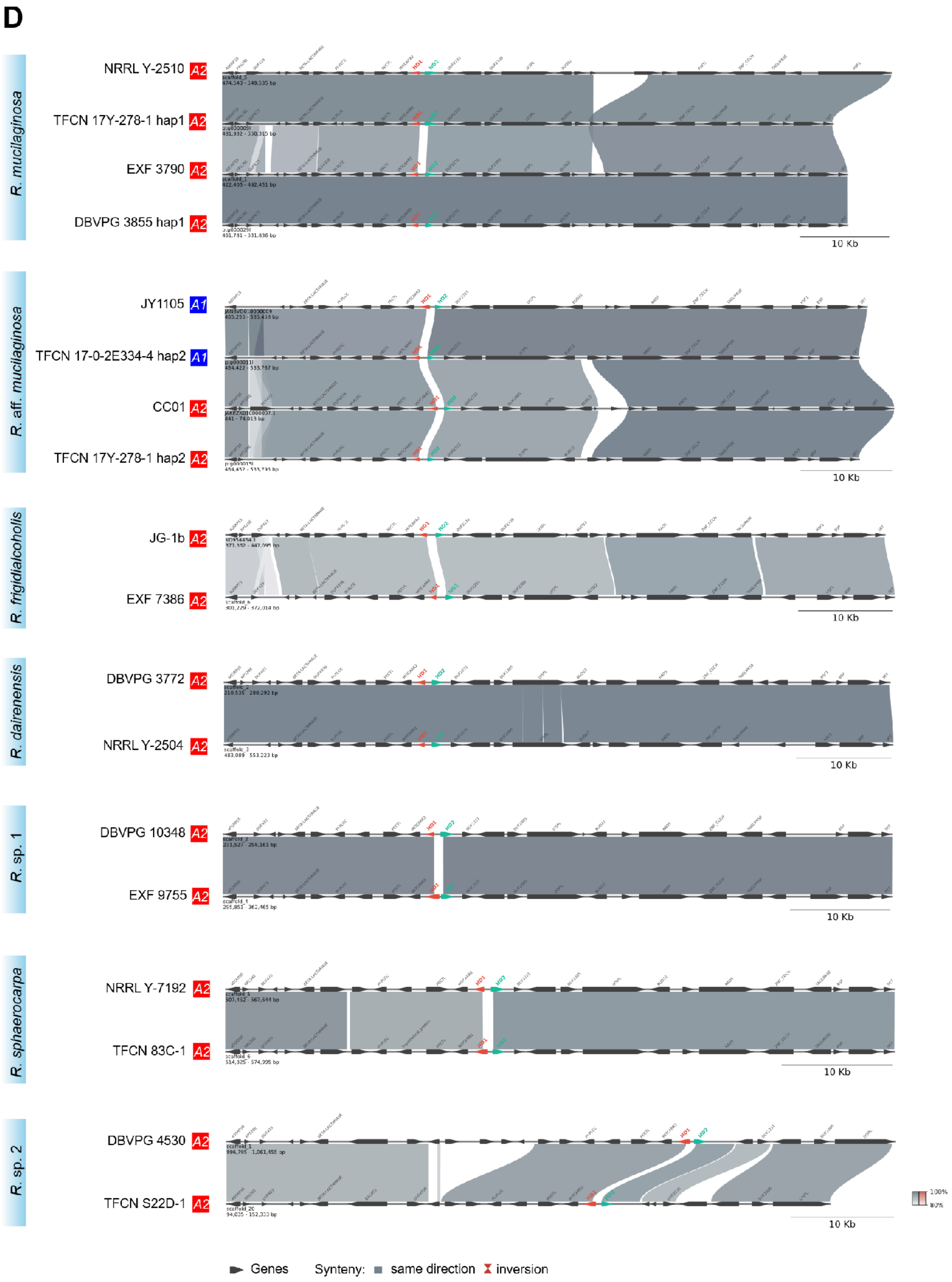
D
R. mucilaginosa
NRRL Y-2510 A2
TFCN 17Y-278-1 hap1 A2
EXF 3790 A2
DBVPG 3855 hap1 A2
R. aff. mucilaginosa
JY1105 A1
TFCN 17-0-2E334-4 hap2 A1
CC01 A2
TFCN 17Y-278-1 hap2 A2
R. frigidialcoholis
JG-1b A2
EXF 7386 A2
R. dairenensis
DBVPG 3772 A2
NRRL Y-2504 A2
R. sp. 1
DBVPG 10348 A2
EXF 9755 A2
R. sphaerocarpa
NRRL Y-7192 A2
TFCN 83C-1 A2
R. sp. 2
DBVPG 4530 A2
TFCN S22D-1 A2
10 Kb
HD1
HD2
100%
80%
Genes
Synteny: same direction inversion

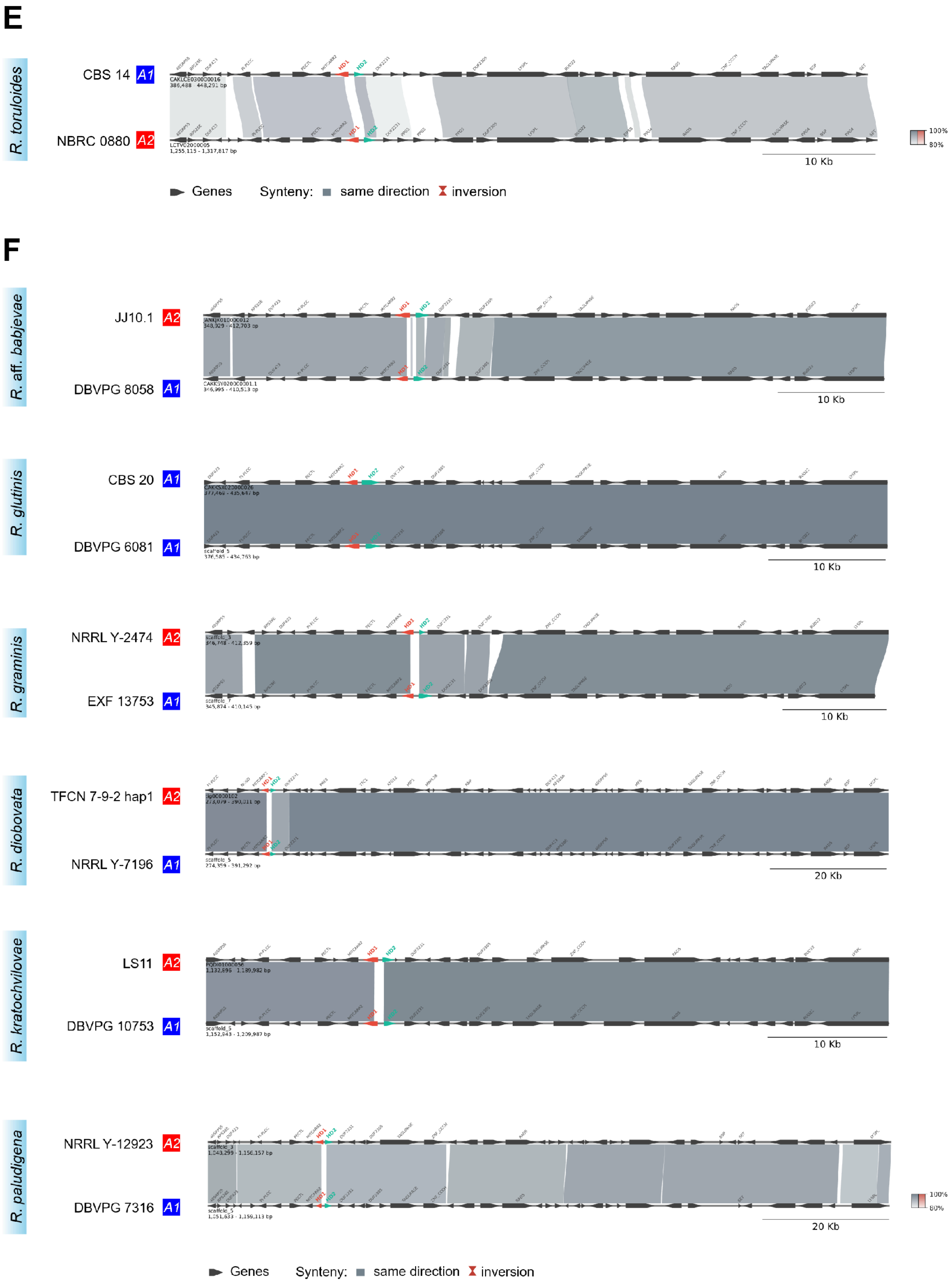

E
R. toruloides
CBS 14 A1
NBRC 0880 A2
10 Kb
Genes
Synteny: same direction
inversion
F
R. aff. babjevae
JJ10.1 A2
DBVPG 8058 A1
10 Kb
R. glutinis
CBS 20 A1
DBVPG 6081 A1
10 Kb
R. graminis
NRRL Y-2474 A2
EXF 13753 A1
10 Kb
R. diobovata
TFCN 7-9-2 hap1 A2
NRRL Y-7196 A1
20 Kb
R. kratochvilovae
LS11 A2
DBVPG 10753 A1
10 Kb
R. paludigena
NRRL Y-12923 A2
DBVPG 7316 A1
20 Kb
Genes
Synteny: same direction
inversion

**Supplementary Figure 8. Synteny and genomic architecture of *P/R* and *HD* loci across *Rhodotorula* species. (A–C)** Comparative synteny of the *P/R* locus in Clades A, B, and C, respectively (Pages 1–3). **(D–F)** Comparative synteny of the *HD* locus in Clades A, B, and C, respectively (Pages 4–6). Genes are represented by block arrows indicating transcription orientation. Transposable elements (TEs) and repeat regions are shown in pink; predicted *P/R* locus boundaries are highlighted in yellow. Connecting colored shades between adjacent genomes represent syntenic collinear regions (gray) and structural inversions (red).

A

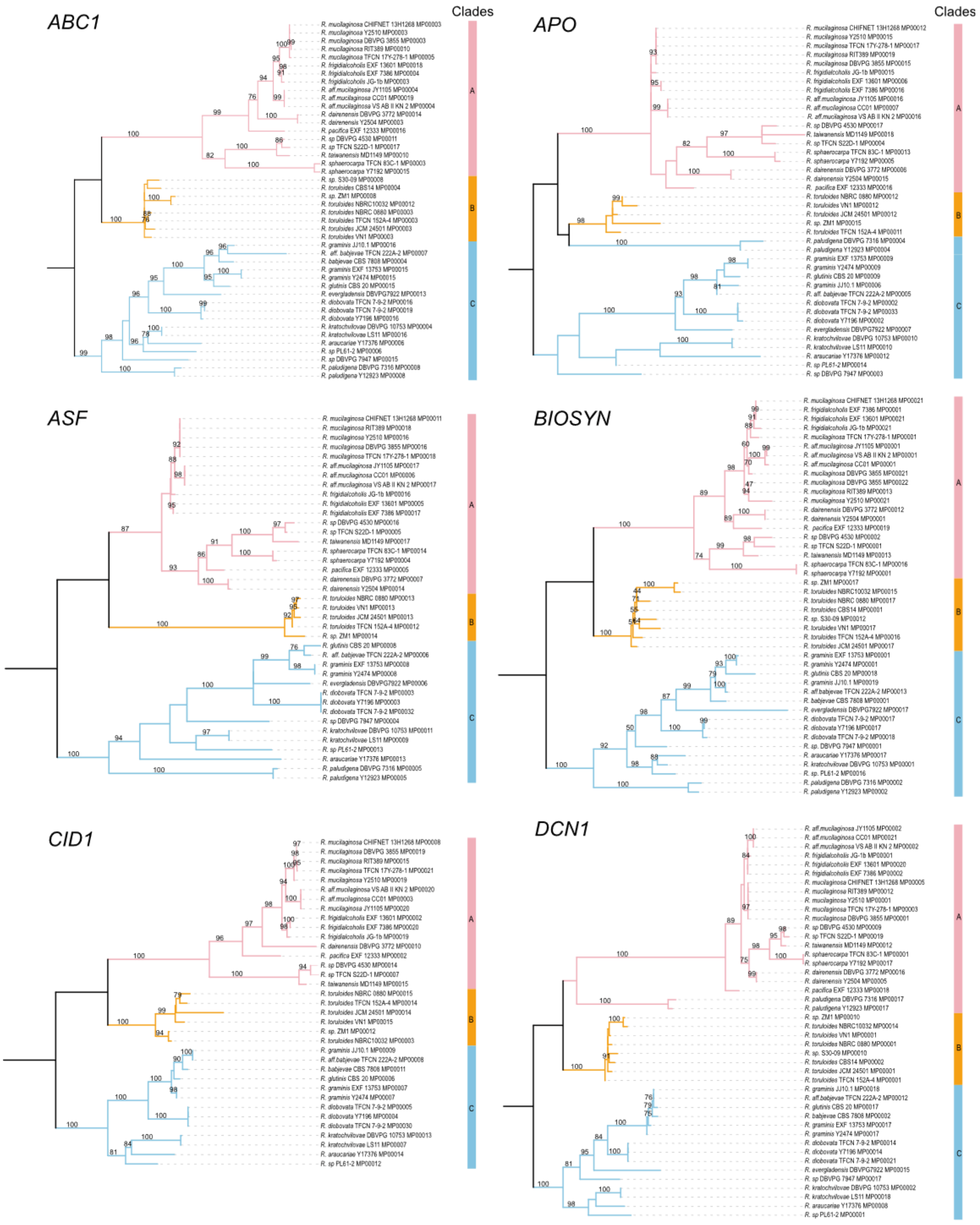

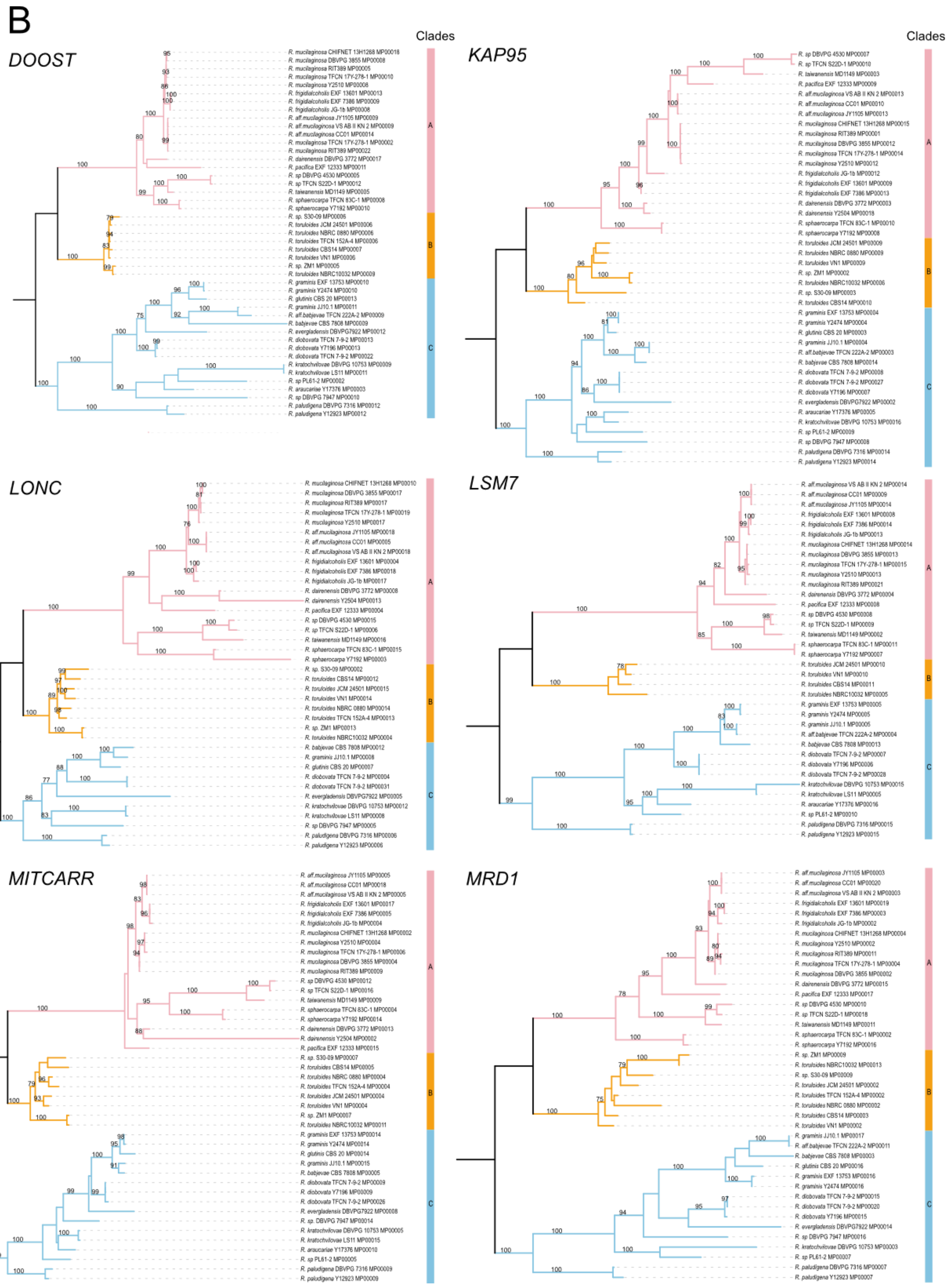
B
DOOST
KAP95
LONC
LSM7
MITCARR
MRD1
Clades
Clades

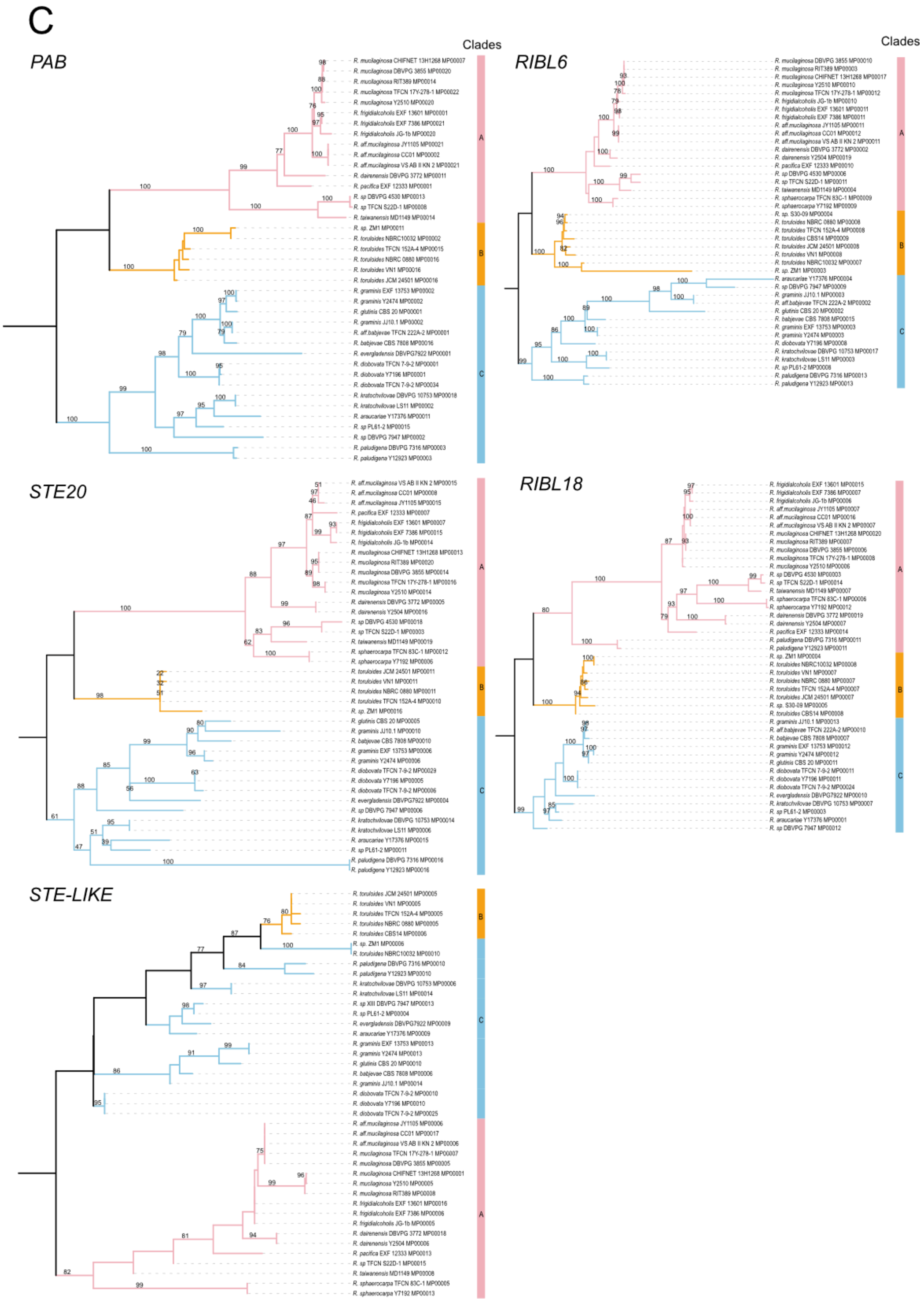
C
PAB
Clades
RIBL6
Clades
STE20
RIBL18
STE-LIKE

**Supplementary Figure 9. Phylogenetic analysis of genes within and adjacent to the *MAT* locus across *Rhodotorula* species.**

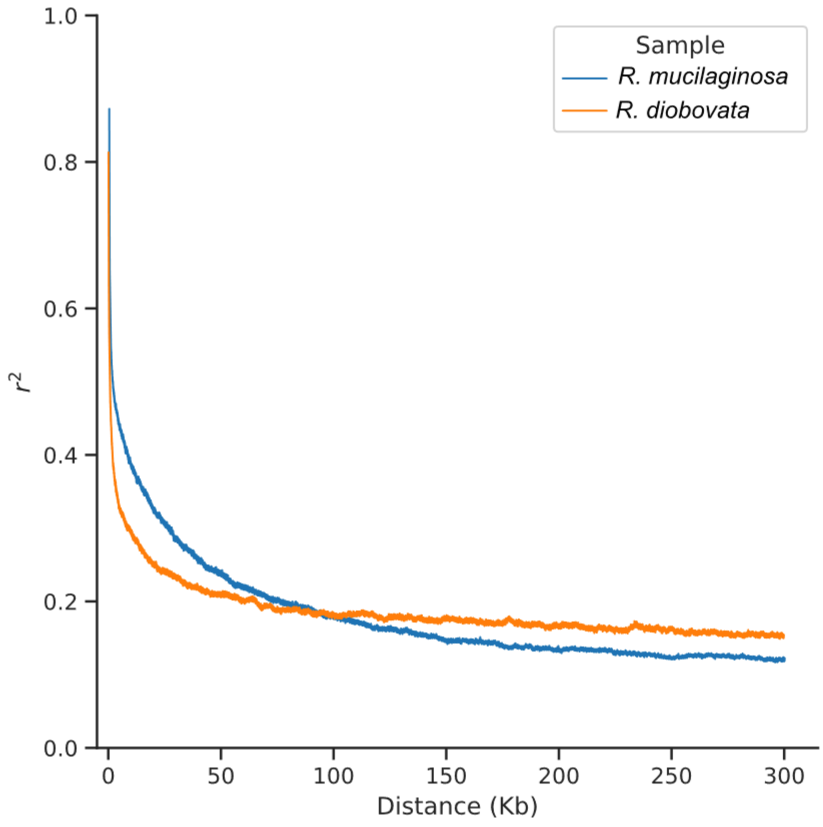


**Supplementary Figure 10. Linkage disequilibrium (LD) decay in *R. mucilaginosa* and *R. diobovata* populations.** Pairwise linkage disequilibrium ($r^2$) plotted across a 300 bp genomic window for *R. mucilaginosa* and *R. diobovata* populations.

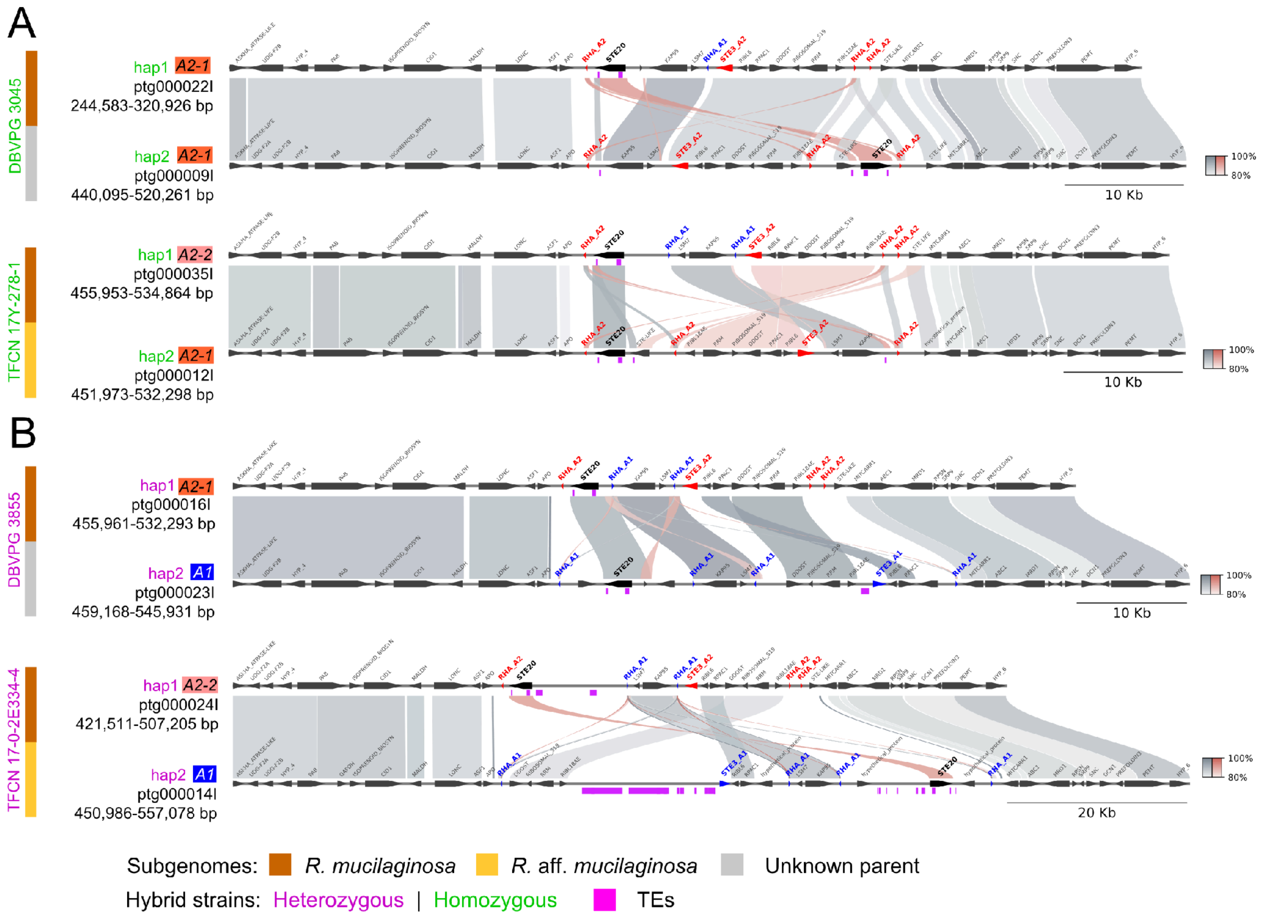


**Supplementary Figure 11. The *P/R* locus architecture in *Rhodotorula mucilaginosa* hybrid strains.** Synteny analysis of the *P/R* locus between phased subgenomes (hap1 and hap2) of **(A)** homozygous (A2+A2; green) and **(B)** heterozygous (A1+A2; purple) *R. mucilaginosa* hybrid strains. Colored blocks on the left indicate inferred parental origin.

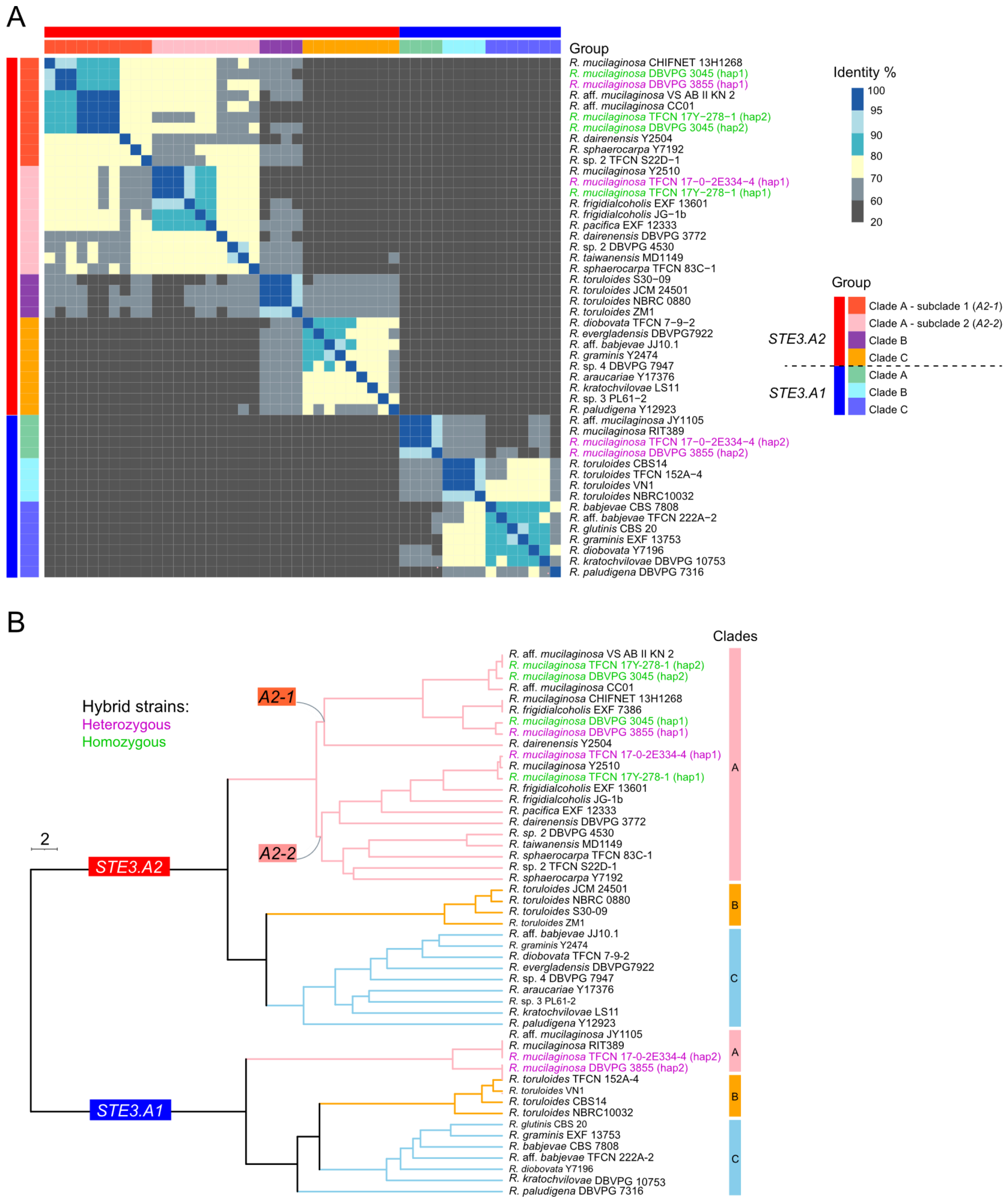


**Supplementary Figure 12. Sequence divergence of *STE3* alleles across *Rhodotorula*. (A)** Heatmap of pairwise sequence identities among *STE3* alleles. **(B)** UPGMA dendrogram depicting hierarchical clustering based on pairwise nucleotide distance.

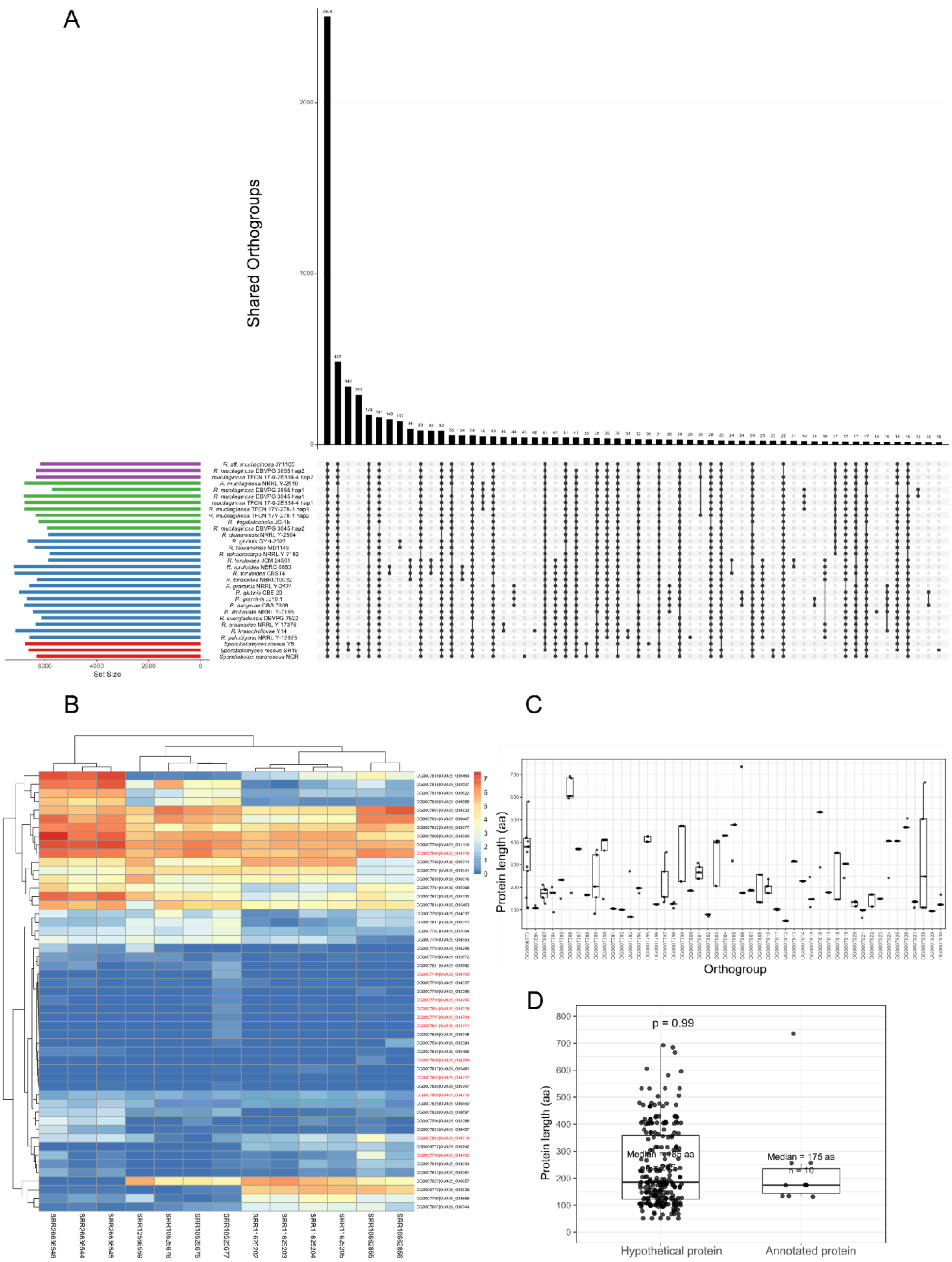
A
Shared Orthogroups
Set Size
B
C
Protein length (aa)
Orthogroup
D
p = 0.99
Median = 175 aa
Protein length (aa)
Hypothetical protein
Annotated protein

**Supplementary Figure 13. Identification, expression, and structural characteristics of A2-associated orthogroups in the *Rhodotorula mucilaginosa* species complex. (A)** UpSet plot showing 49 orthogroups exclusively shared by *R. mucilaginosa* Y-2510 and A2-type hap1 subgenomes of four hybrids, but absent in hap2 subgenomes and other *Rhodotorula* species. **(B)** Express profiles of genes across these orthogroups across 13 public RNA-seq datasets, with genes within the 11-orthogroup accessory block highlighted in red. **(C)** Protein-length distributions across 49 orthogroups. **(D)** Comparison of protein lengths between hypothetical and functionally annotated proteins.

A

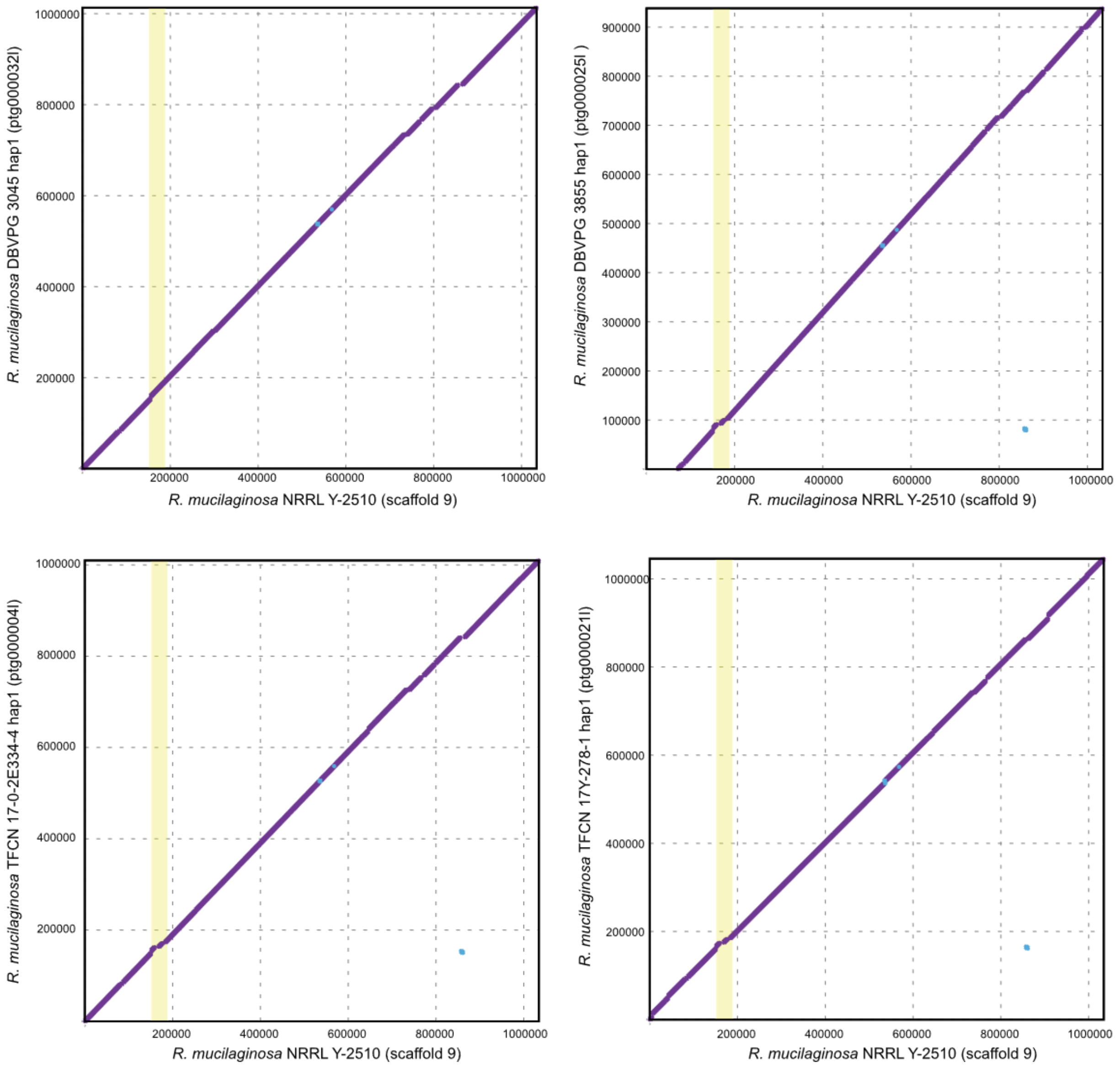
R. mucilaginosa DBVPG 3045 hap1 (ptg000032l)
R. mucilaginosa NRRL Y-2510 (scaffold 9)
R. mucilaginosa DBVPG 3855 hap1 (ptg000025l )
R. mucilaginosa NRRL Y-2510 (scaffold 9)
R. mucilaginosa TFCN 17-0-2E334-4 hap1 (ptg000004l)
R. mucilaginosa NRRL Y-2510 (scaffold 9)
R. mucilaginosa TFCN 17Y-278-1 hap1 (ptg000021l)
R. mucilaginosa NRRL Y-2510 (scaffold 9)

B

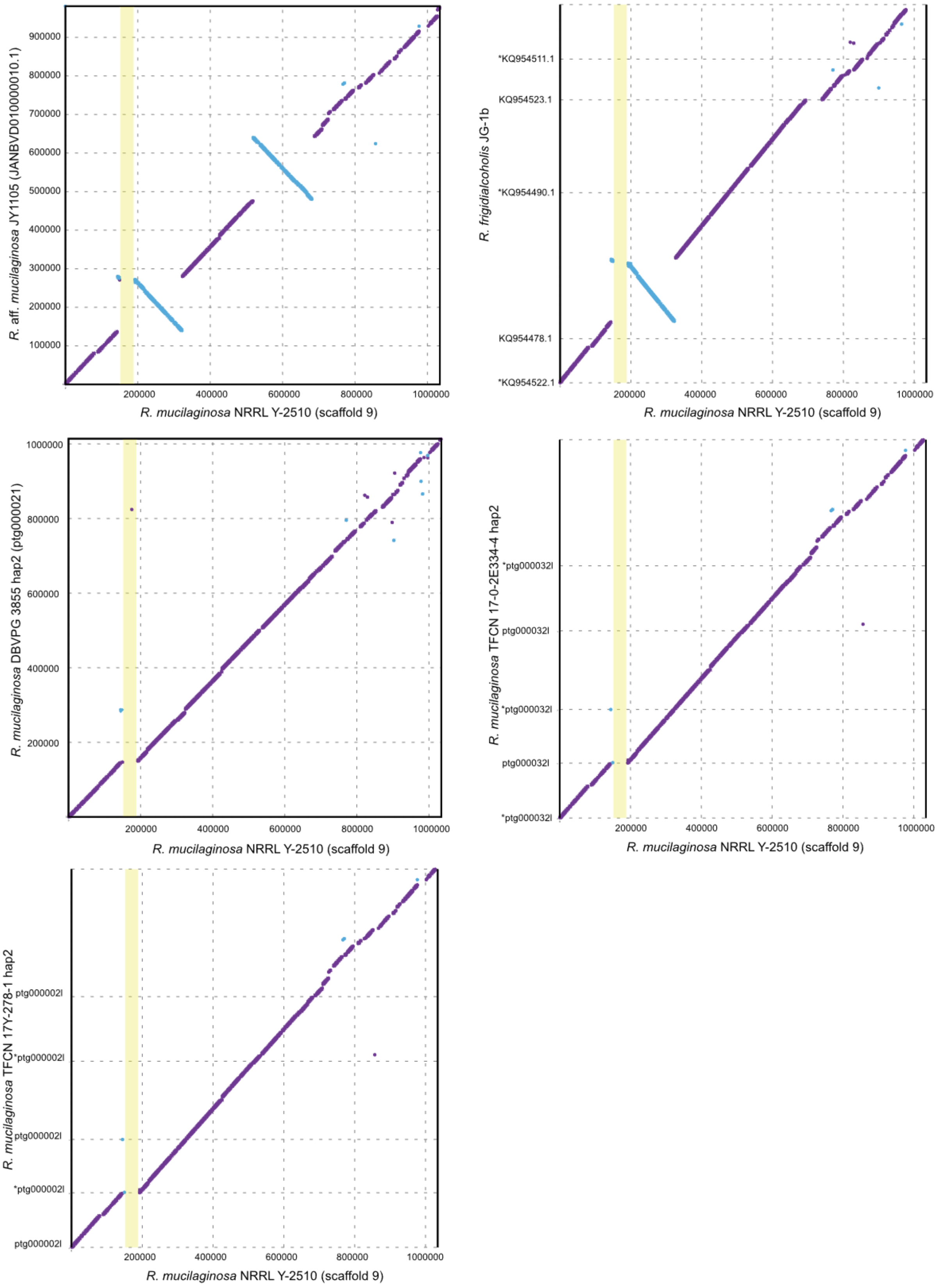

R. aff. mucilaginosa JY1105 (JANBVD010000010.1)
900000
800000
700000
600000
500000
400000
300000
200000
100000
200000
400000
600000
800000
1000000
R. mucilaginosa NRRL Y-2510 (scaffold 9)
R. frigidialcoholis JG-1b
*KQ954511.1
KQ954523.1
*KQ954490.1
KQ954478.1
*KQ954522.1
R. mucilaginosa NRRL Y-2510 (scaffold 9)
R. mucilaginosa DBVPG 3855 hap2 (ptg000021)
1000000
800000
600000
400000
200000
R. mucilaginosa NRRL Y-2510 (scaffold 9)
R. mucilaginosa TFCN 17-0-2E334-4 hap2
*ptg000032l
ptg000032l
*ptg000032l
ptg000032l
*ptg000032l
R. mucilaginosa NRRL Y-2510 (scaffold 9)
R. mucilaginosa TFCN 17Y-278-1 hap2
ptg000002l
*ptg000002l
ptg000002l
*ptg000002l
ptg000002l
R. mucilaginosa NRRL Y-2510 (scaffold 9)

C

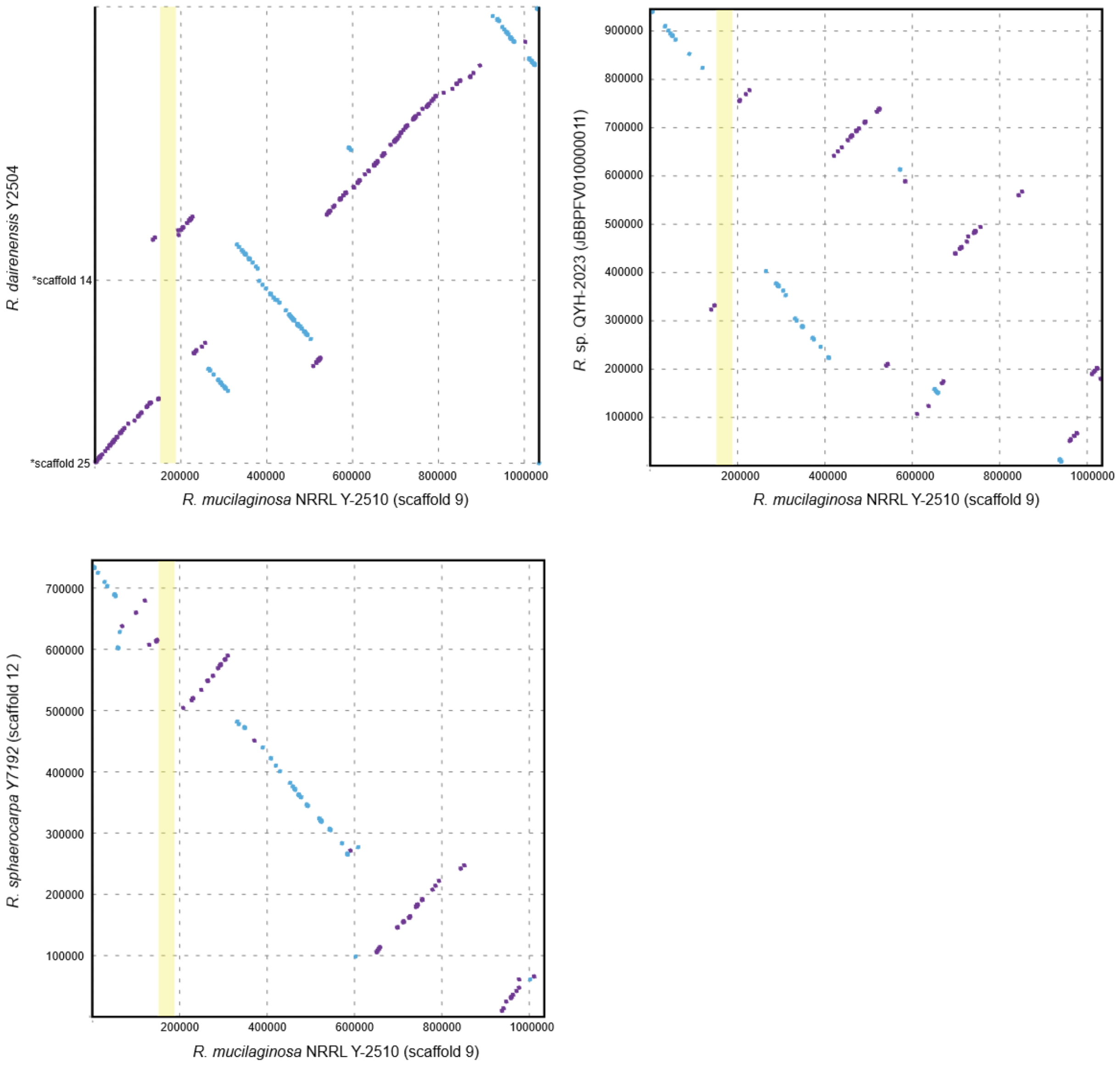


**Supplementary Figure 14. Conservation and lineage-specific distribution of the 11-orthogroup accessory genomic block.** Synteny analysis shows that the intact collinear 11-orthogroup block is restricted to the A2-type *R. mucilaginosa* genome and A2-type hap1 hybrid subgenomes, while absent from hap2 subgenomes and other Clade A species. **(A)** Synteny comparison across A2-type subgenomes from hybrids, illustrating conserved gene order and orientation. **(B)** Alignment with syntenic regions in hybrid hap2 subgenomes and closely related species (*R. frigidialcoholis* and *R.* aff. *mucilaginosa*). **(C)** Alignment with homologous regions across other Clade A species.

### Supplementary text S1

#### Genomic dataset composition and assembly quality

To improve taxonomic representation across *Rhodotorula* species, we assembled a high-quality dataset of 20 genomes. At the onset of this study, only four recognized species had publicly available type-strain assemblies (*R. frigidialcoholis* JG-1b, *R. toruloides* CBS14, *R. babjevae* CBS 7808 and *R. glutinis* CBS 20). To fill this gap, we newly sequenced eight type strains using a hybrid Oxford Nanopore (long-read) and Illumina (short-read) sequencing strategy. The remaining 12 public genomes comprise the four previously available type strains, three representatives of the *R. toruloides* species complex (NBRC 0880, NBRC 10032, and JCM 24501), two strains corrected for misidentification in this study—*R. "glutinis"* QYH-2023 (*Rhodotorula sp. 2*) and *R. "graminis"* JJ10.1 (*R. aff. babjevae*)—which represent putative novel species, two genomes retrieved directly from NCBI without affiliated publications (*R. aff. Mucilaginosa* JY1105 and *R. kratochvilovae* Y14), and *R. taiwanensis* MD1149. To maintain analytical rigor, dataset inclusion was limited to genomes assembled via multi-platform or long-read technology combinations (ONT, PacBio, Illumina mate-pair, 454, or Illumina). All 20 genomes demonstrated high assembly quality, with BUSCO completeness exceeding 95% and scaffold N50 values surpassing 1 Mb for all strains except *R. taiwanensis* MD1149 and *R. frigidialcoholis* JG-1b (Table S1).

The 249 *Rhodotorula* genome assemblies exhibited high completeness, with BUSCO scores exceeding 90% for all strains except *R. kratochvilovae* LS11, *R. kratochvilovae* YM25235, and *R. mucilaginosa* TFCN 3M-1-1 (84.5–87.8%). The mean scaffold N50 across the dataset was 0.43 Mb. Genome assemblies spanned 17.4–42.7 Mb in size with GC content ranging from 58.7% to 68.5% (Table S2).

**Phylogenomic reconstruction, concordance, and localized discordance in *Rhodotorula***

We reconstructed *Rhodotorula* phylogenetic relationships using two complementary approaches: STAG species-tree inference and Phyling concatenated alignment analysis. For the STAG approach, 7,999 gene trees were constructed, of which 3,152 species-ubiquitous trees were used for consensus species-tree inference. For Phyling, single-copy orthologs were identified against the BUSCO fungi_odb10 marker set, resulting in a concatenated alignment of 746 single-copy orthologs used for partitioned maximum-likelihood phylogenetic inference. Both approaches yielded congruent backbone topologies that support the three-clade framework of the genus (Fig. 1).

However, localized method-dependent placements occurred within recently diverged lineages. For example, *R. mucilaginosa* Y-2510 was placed sister to *R. frigidialcoholis* JG-1b in the STAG tree but placed sister to *R.* aff. *mucilaginosa* JY1105 in the Phyling tree. In the expanded 249-strain dataset, these three species could not be clearly resolved due to strain-level placement instability (Fig. S1). While Phyling supported all branches with > 96% bootstrap confidence, STAG yielded lower bipartition support within the *R. mucilaginosa* complex. These metrics reflect distinct properties—STAG measures gene-tree concordance across loci, whereas Phyling bootstrap values assess statistical confidence from alignment site resampling.

#### Genome size, GC content, ANI, and dDDH

We compared genome size and GC content across Clades A, B, and C using the full set of 249 strains. ANOVA followed by Tukey's HSD post hoc tests revealed significant differences in both genome size and GC content among the three clades at a confidence level of $p < 0.05$. At a confidence level of $p < 0.01$, Clade A had significantly smaller genome sizes than Clades B and C, whereas Clade C had significantly higher GC content than Clades A and B (Fig. S2).

To complement the pairwise ANI patterns observed across species complexes, we calculated digital DNA-DNA hybridization (dDDH) values for representative strains of each species. All interspecific dDDH values fell below 30%, well beneath the standard 70% species-delimitation threshold in yeast (Libkind et al. 2020). However, within the three species complexes exhibited a higher interspecific average dDDH value (25.7%) compared to non-complex interspecific pairs (20.1%) (Table S3). These results support that taxa within these complexes represent distinct, recently diverged species.

#### Composition of core *P/R*-locus genes

In addition to the core mating-related genes such as the pheromone receptor gene *STE3* and the pheromone precursor gene *RHA*, several genes involved in mating and filamentation have been anciently recruited to the *P/R* locus across diverse taxa, including *STE11*, *STE12*, and *STE20* (Fraser et al. 2004; Coelho et al. 2010; Sun et al. 2019). Among these genes, only *STE20* was consistently found within the *P/R* locus of *Rhodotorula* species, except in the fragmented assembly of *R*. aff. *mucilaginosa* A2-type strain CC01. A *STE12*-like gene was also present within the core *P/R* region across the genus, but its mating-type association varied among clades. It was linked to the *P/R* A2 allele in Clade A and B, whereas associated with the *P/R* A1 allele in Clade C (Table S5). Despite substantial sequence divergence across clades, these genes retained characteristic domain of *STE*-like transcription factor. We therefore designated them as *STE*-like in the synteny plots (Fig. S8).

#### Structural divergence between alternative *P/R* alleles

Comparisons between alternative A1 and A2 *P/R* alleles revealed multiple rearranged modules spanning the entire *P/R* locus. In *R. paludigena*, for example, the A1 and A2 *P/R* alleles differed by four inverted modules, including regions containing *KAP95-LSM7*, *RIBL6-RPAC1*, *DDOST-RIBOSOMAL_S19-RRM-RIBL18AE* and *MITCARR1-ABC1-DCN1-SNC-SRP9-MRD1* (Fig. 3 and

Fig. S8C). These rearrangements provided a representative example of the large-scale structural divergence observed between compatible mating-type alleles.

#### Lineage- and strain-specific rearrangements within *P/R* allele classes

Within otherwise conserved *P/R* allele classes, several lineage- or strain-specific rearrangements were observed. Among A2 *P/R* alleles, these included repositioning of *STE20* in *R. toruloides* NBRC 0880 and inversion of the *LSM7-KAP95* module in *R. sphaerocarpa* NRRL Y-7192 and *Rhodotorula* sp. 2 TFCN S22D-1. In Clade C species, *STE20* remained in the same relative position within the *P/R* locus but was oriented oppositely to its Clade A counterparts. Among A1 *P/R* alleles, *STE20* inversions were observed in *R. kratochvilovae* DBVPG 10753, *R. diobovata* NRRL Y-7196, *R. babjevae* CBS 7808, and *R.* aff. *babjevae* DBVPG 8058. An inverted module containing *MRD1*, *SRP9*, *SNC*, *DCN1*, and A*BC1* was observed in *R. graminis* EXF 13753, *R. glutinis* CBS 20, and *R.* aff. *babjevae* DBVPG 8058.

Strain-specific rearrangements were also detected among A2 mating-type strains within the same Clade A species (Fig. S8A). A short inversion involving *KAP95*, *LSM7*, and *RHA* was identified in *R. mucilaginosa*, whereas larger inversions were detected in *R.* aff. *mucilaginosa*, *R. frigidialcoholis*, and *R. dairenensis*. These inversions involved different combinations of *STE20*, *RHA*, *RIBL6*, *RPAC1*, *DDOST*, *RRM*, *RIBL18AE*, *LSM7*, and *KAP95*. In the A1 strain JY1105 of *R.* aff. *mucilaginosa*, two regions, *ASF1-APO-LONC* and *DDOST-RIBOSOMAL_S19-RRM-RIBL18AE*, were duplicated relative to the A1-type subgenome of TFCN 17-0-2E334-4 and to A2 counterparts including CC01 and the hap2 subgenome of TFCN 17Y-278-1, with one copy undergoing inversion (Fig. S8A).
